# Diversity and functional specialization of oyster immune cells uncovered by integrative single cell level investigations

**DOI:** 10.1101/2024.07.19.604245

**Authors:** Sébastien de La Forest Divonne, Juliette Pouzadoux, Océane Romatif, Caroline Montagnani, Guillaume Mitta, Delphine Destoumieux-Garzon, Benjamin Gourbal, Guillaume M. Charrière, Emmanuel Vignal

## Abstract

Mollusks are a major component of animal biodiversity and play a critical role in ecosystems and global food security. The Pacific oyster, *Crassostrea (Magallana) gigas*, is the most farmed bivalve mollusk in the world and is becoming a model species for invertebrate biology. Despite the extensive research on hemocytes, the immune cells of bivalves, their characterization remains elusive. Here we were able to extensively characterize the diverse hemocytes and identified at least seven functionally distinct cell types and three hematopoietic lineages. A combination of single-cell RNA sequencing, quantitative cytology, cell sorting, functional assays and pseudo-time analyses was used to deliver a comprehensive view of the distinct hemocyte types. This integrative analysis enabled us to reconcile molecular and cellular data and identify distinct cell types performing specialized immune functions, such as phagocytosis, reactive oxygen species production, copper accumulation, and expression of antimicrobial peptides. This study emphasized the need for more in depth studies of cellular immunity in mollusks and non-model invertebrates and set the ground for further comparative immunology studies at the cellular level.

## Introduction

*Mollusca* is the second largest invertebrate phylum, after Arthropoda, and the largest marine phylum, comprising approximately 23 % of all known marine organisms (*1*). Among them, bivalves exhibit a high diversity and a rich evolutionary history (*2*). The Pacific oyster *Crassostrea (Magallana) gigas* (*C. gigas* - Thunberg, 1793) (NCBI:txid29159) is a sessile filter-feeding bivalve that thrives in a variety of stressful environments ranging from intertidal to deep-sea conditions (*3*). It is a key species for the aquaculture industry worldwide (*4*). Several infectious diseases affect *C. gigas* at different life stages, which impacts its production. Given the significant socio-economic value of this species, there has been an increased focus on understanding and mitigating these diseases (*5*). The causes of these mortalities can involve a variety of pathogens, including viruses, bacteria and parasites that can be responsible for the mortality events affecting *C. gigas* (*6*). One of the most extensively researched infectious diseases is POMS (Pacific Oyster Mortality Syndrome), a polymicrobial disease responsible for mass mortalities of juvenile oysters (*7*). The disease is triggered by the OsHV-1 μVar herpesvirus, which alters the immune defenses of oysters, allowing the colonization of opportunistic bacteria, including *Vibrio*, that cause hemocyte lysis and bacteremia, ultimately leading to animal death (*7*). Other bacterial pathogens have also been identified as a contributing factor in mass mortalities of adult Pacific oysters in several countries. The most notable is *Vibrio aestuarianus* which affects adult oysters in Europe (*8*, *9*). To date, the majority of oyster pathogens or opportunistic pathogens that have been characterized in detail have been found to subvert hemocyte defenses for their own benefit. These include the OsHV-1 μVar herpesvirus (*10*) and virulent *Vibrio* strains of the species *Vibrio crassostreae* and *Vibrio tasmaniensis* (*11*), *Vibrio aestuarianus* and *Vibrio harveyi* (*12*), highlighting the critical role of hemocytes in oyster immunity. The development of immune-based prophylactic treatments, such as immune-priming and immune-shaping, represents a promising avenue for enhancing the natural defenses of oysters against pathogens and increasing their survival rate. Nevertheless, the advancement of such therapies is still constrained by a dearth of knowledge regarding the underlying molecular and cellular mechanisms (*13*, *14*).

The study of hemocytes has a long history, dating back to the 1970s (for review see (*15*)). Hemocytes are cellular effectors of the immune system. They engage in phagocytosis to engulf and destroy potential pathogens, neutralizing parasites by encapsulation, or preventing pathogen dissemination by cell aggregation and the release of extracellular DNA traps (*16*). Furthermore, they engage in the humoral response by releasing cytokines, antimicrobial peptides, and reactive oxygen species (ROS), which enable them to combat pathogens (*17*). In addition to their role in oyster immunity, hemocytes have been implicated in numerous physiological processes, including shell repair (*18*), wound healing, nutrient transport, and environmental contaminant removal (*19*). Despite this acquired knowledge, hemocytes remain an under-characterized population of circulating immune cells. The lack of a unified classification and of molecular and functional genetic tools hinders our understanding of lineage ontogeny and functional specialization. Several studies have proposed different classifications of hemocytes in the *Ostreidae* family, with 3 to 4 hemocyte types reported (*15*). These classifications are primarily based on either microscopic or flow cytometry analyses. In *C. gigas*, three primary hemocyte cell types have been classically identified : blast, hyalinocyte, and granulocyte cells. Blast-like cells are considered as undifferentiated hemocyte types (*20*), hyalinocytes (*21*) seem to be more involved in wound repair, and granulocytes, more implicated in immune surveillance. The latter are considered as the main immunocompetent hemocyte types (*22*). While the immune response of *C. gigas* has been extensively studied using classical transcriptomics (*23–25*) at the whole animal, tissue, or circulating hemocyte levels, these approaches have failed to consider the diversity of hemocyte cell types and lineages that underpin these responses (*26*). However, it is still imperative to accurately describe the diversity of these cells, understand their ontogeny, and delineate cell lineages to comprehend their specific roles. The advent of single-cell RNA sequencing (scRNA-seq) techniques has enabled the monitoring of global gene expression at the single-cell level with thousands of individual cells in a single experiment. This provides a unique opportunity to overcome these limitations and deepen our understanding of hemocyte diversity and function in bivalves. Recently scRNA-seq was used to provide a first molecular description of a hemocyte population in the oyster *Crassostrea hongkongensis* (*27*). However, in the absence of morphological and/or functional characterization studies, the authors could not deduce the hemocyte cell types to be matched to the transcriptomic profiles generated by scRNA-seq.

While numerous transcriptomic analyses have been conducted on *C. gigas* hemocytes, none have adopted a single-cell approach. In this study, we present an integrative analysis of the diversity of *C. gigas* hemocytes at the single-cell level. To this end, we combined scRNA-seq from a pathogen-free adult oyster combined with cytological, cell fractionation and functional assays. This approach allowed us to create a comprehensive transcriptomic, cytological and functional atlas of hemocyte cell types. Our scRNA-seq analyses identified 7 distinct transcriptomic populations and functional annotation revealed distinct populations with specific functions, including phagocytosis, oxidative burst, energetic metabolism, enhanced transcription, translation and cell division. Quantitative cytology enabled the identification of 7 morphologically distinct hemocyte cell types, which allowed us to reconcile molecular and cytological data. Density gradients were used to separate hemocyte cell types and qPCR or functional assay analyses were performed to validate cell type-specific markers. By employing this integrated approach, we could identify 1 type of hyalinocyte, 2 types of blasts and 4 types of granular cells. Furthermore, we identified cell types that perform antimicrobial functions through phagocytosis, ROS production, copper accumulation, and expression of antimicrobial peptides. Finally, trajectory analysis of scRNA-seq data combined with functional analysis revealed distinct differentiation pathways that may control hemocyte ontology and differentiation processes. Based on these findings, we propose a more comprehensive and up-to-date classification of *C. gigas* hemocytes, with a more accurate description of the different cell types, their potential ontology and a precise description of their sub-functionalization.

## Results

### Single-cell RNA sequencing reveals 7 distinct transcriptomic clusters of circulating immune cells in oysters

Oysters are known to exhibit a high degree of individual genetic polymorphism, including Copy Number Variation (CNV) and Presence Absence Variation (PAV) (*28*). To prevent misinterpretation of the single-cell transcriptomic data, and to characterize the hemocyte cell types and their heterogeneity, we sampled hemocytes from a unique known pathogen-free animal (Ifremer Standardized Animal, 18-month-old) and applied single-cell drop-seq technology. A total of 4,950 cells were loaded, with a target recovery of 3,000 single hemocytes for analysis. **(Fig. 1A)**. The scRNA-seq library was generated and sequenced, resulting in 127,959,215 high-quality filtered reads available for single-cell analysis (ENA project accession number PRJEB74031). Primary bioinformatics analysis was performed using the STAR solo aligner software (*29*) against the *C. gigas* genome (Genbank reference GCA_902806645.1) from the Roslin Institute (*30*). Of the 127,959,215 reads, 97 % showed a valid barcode and 89.2 % were successfully mapped to the genome with a saturation of 75.6 %. A total of 2,937 cells were profiled, yielding a median of 1,578 genes and 4,412 unique molecular identifiers (UMIs) per cell among the 23,841 total genes detected, with a sequencing saturation of 75.6 % **(Supp. Table S1)**. Secondary bioinformatic data processing was conducted using the Seurat R package (version 4.3.0) (*31*). The data set was filtered to remove data corresponding to empty droplets or cell doublets. Cells with a gene number between 750 and 4,000 and less than 5 % mitochondrial genes were retained. After quality control processing, 120 cells corresponding to empty droplets and cell doublets were removed, and 2,817 cells were processed for data normalization. Finally, we performed linear dimensional reduction and clustering on the 3,000 most variable genes from 2,817 cells **(Supp. Fig. S1)**. Dimension reduction and clustering led to identifying 7 transcriptomic clusters, within which hemocytes were distributed. These 7 different clusters represented 27.6, 23.1, 17.8, 16.9, 7, 4.6 and 3 % of the total cells **(Fig. 1B)**. For each transcriptomic cluster, the heatmap displays the top 10 positively enriched marker genes identified by Seurat’s differential expression analysis, highlighting cluster-specific overexpression **(Fig. 1C)**.

**Fig. 1.**
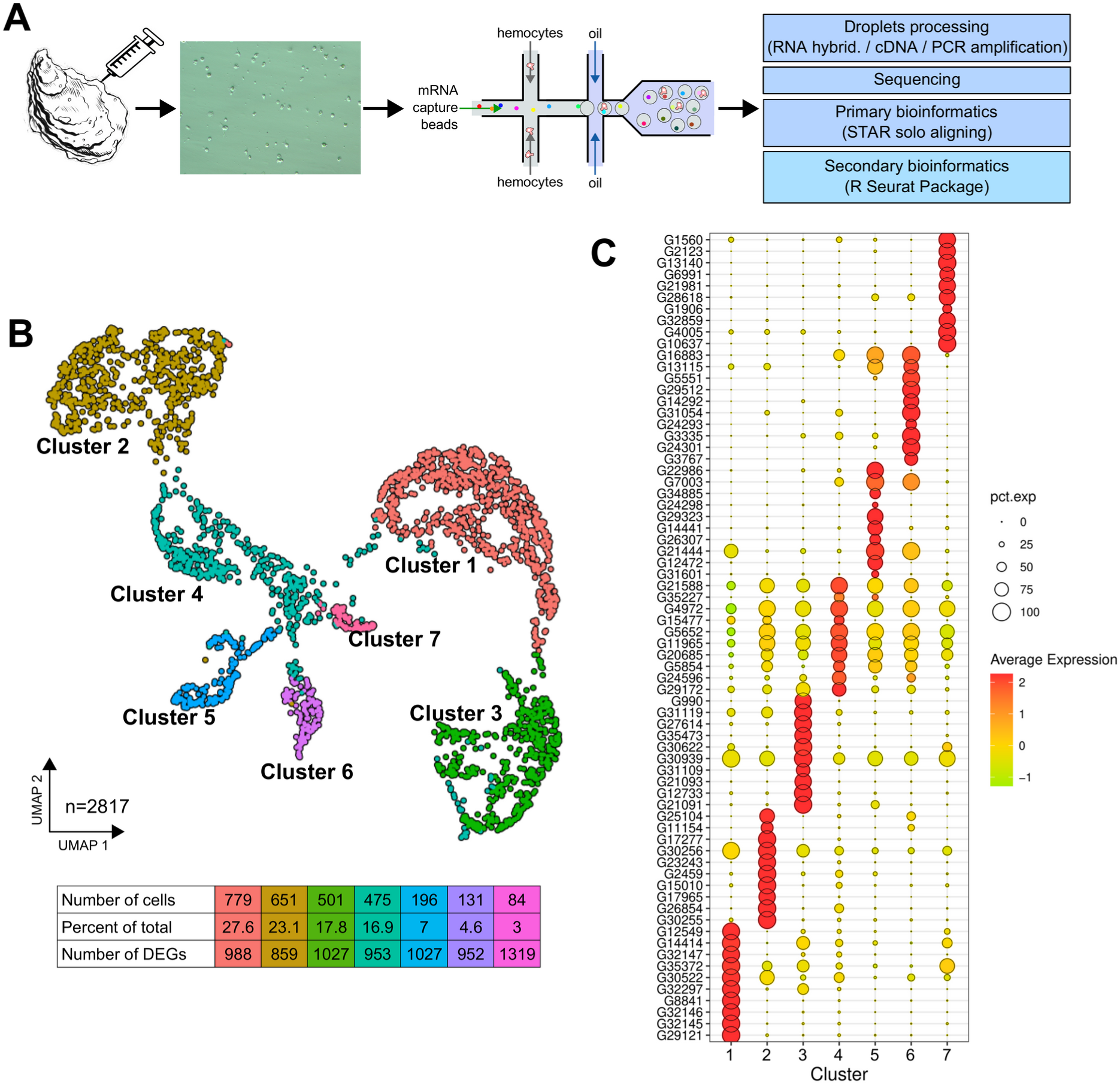
scRNA-seq analysis of *C. gigas* circulating hemocytes reveals 7 transcriptomic cell clusters. **(A)** Schematic of the scRNA-seq 10X Genomics Chromium microfluidic technology and bioinformatics processing workflow used. Dissociated hemocytes were collected from a pathogen-free oyster and encapsulated in droplets for processing. After sequencing, the data were processed bioinformatically. **(B)** Uniform Manifold Approximation and Projection (UMAP) plot for dimensional reduction of the data set and summary of cells and the number of Differentially Expressed Genes (DEGs) in each cluster. The table shows the characteristics (number of cells, percentage of total cells and number of Differentially Expressed Genes in each cluster) of the seven clusters identified. **(C)** Dot plot representing the ten most enriched DEGs per cluster based on average expression (avg_log2FC). The color gradient of the dot represents the expression level, while the size represents the percentage of cells expressing each gene per cluster.

Average Log2FC values and percentage of expression in each cluster relative to all other clusters were calculated for the ten most differentially expressed genes **(Fig. 1D and Table 1).** Clear transcriptomic cell clusters were detected as well as specific gene markers for each cluster **(Supp. Data 1)**. Only the transcriptomic signature of cluster 4 was less contrasting than that of the other clusters **(Fig. 1D)**.

**Table 1.**
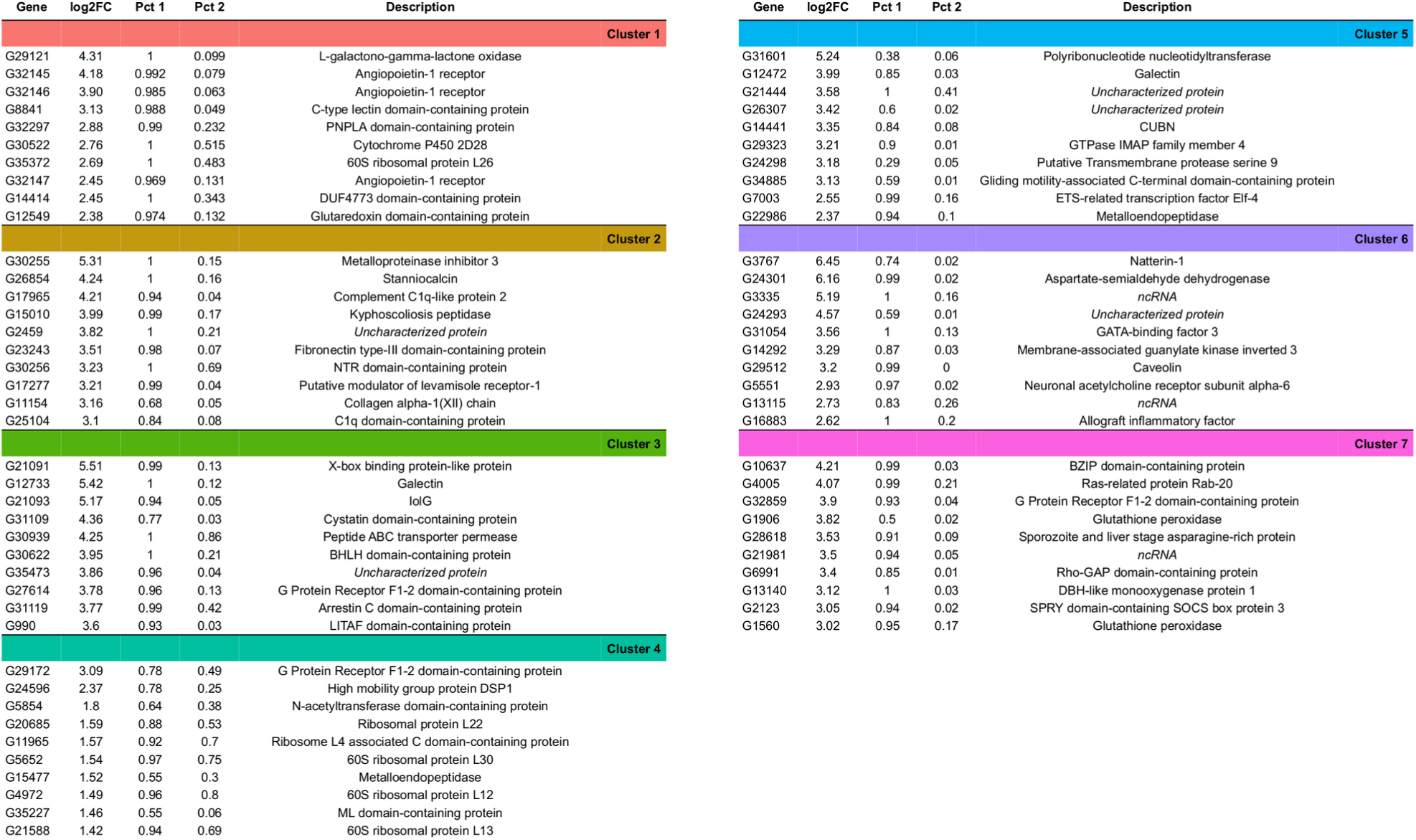
Top 10 overexpressed genes identified in each transcriptomic cluster. The first column indicates the gene number according to the annotation. ‘log2FC’ represents the log2 fold change of the gene in the cluster compared to all other cells. ‘Pct1’ is the percentage of cells expressing the gene in the cluster and ‘Pct2’ is the fraction of cells expressing the gene in all other clusters. The description is the annotation of the expressed gene. (adjusted p-value < 0.05).

### KEGG pathways and GO-terms analyses reveal functional diversity in *C. gigas* hemocytes

The ScRNA-seq data demonstrated that specific functions are carried out by the hemocyte cell types that comprise the seven transcriptomic clusters. A preliminary overview of the functions over-represented in each cluster was obtained through a KEGG pathway analysis on the overrepresented transcripts (Log2FC > 0.25) using the *C. gigas* annotation provided by the DAVID consortium (*32*), thereby identifying pathways specifically enriched in each cluster. Cluster 1 demonstrated enrichment in viral processing and endocytosis **(Figure 2A and Supplemental Table S2)**. Additionally, it demonstrated enrichment in pyruvate metabolism, glycolysis/gluconeogenesis and the pentose phosphate pathways. Clusters 1 and 3 exhibited a distinctive enrichment in carbohydrate metabolism and endocytosis. In particular, cluster 3 exhibited enriched transcripts associated with glycolysis/gluconeogenesis and TCA cycle activities. Cluster 4 was enriched in protein synthesis, including transcription (spliceosome, ribosome, nucleocytoplasmic transport and mRNA surveillance pathway) folding, sorting and degradation of protein pathways. Clusters 2, 5 and 6 exhibited a shared signature of enrichment in ribosome-related genes. Cluster 2 demonstrated a specific enrichment in motor protein-coding genes responsible for cell motility and xenobiotic metabolism. Remarkably, clusters 1, 3, 4 and 5 exhibited enriched oxidative phosphorylation transcripts, whereas the transcripts of cluster 7 were enriched in vesicular trafficking and endo-lysosomal pathways (endocytosis, endosome, phagosome, lysosome, auto and mitophagy). Cluster 7 displayed also the signature of Wnt pathway components.

**Fig. 2.**
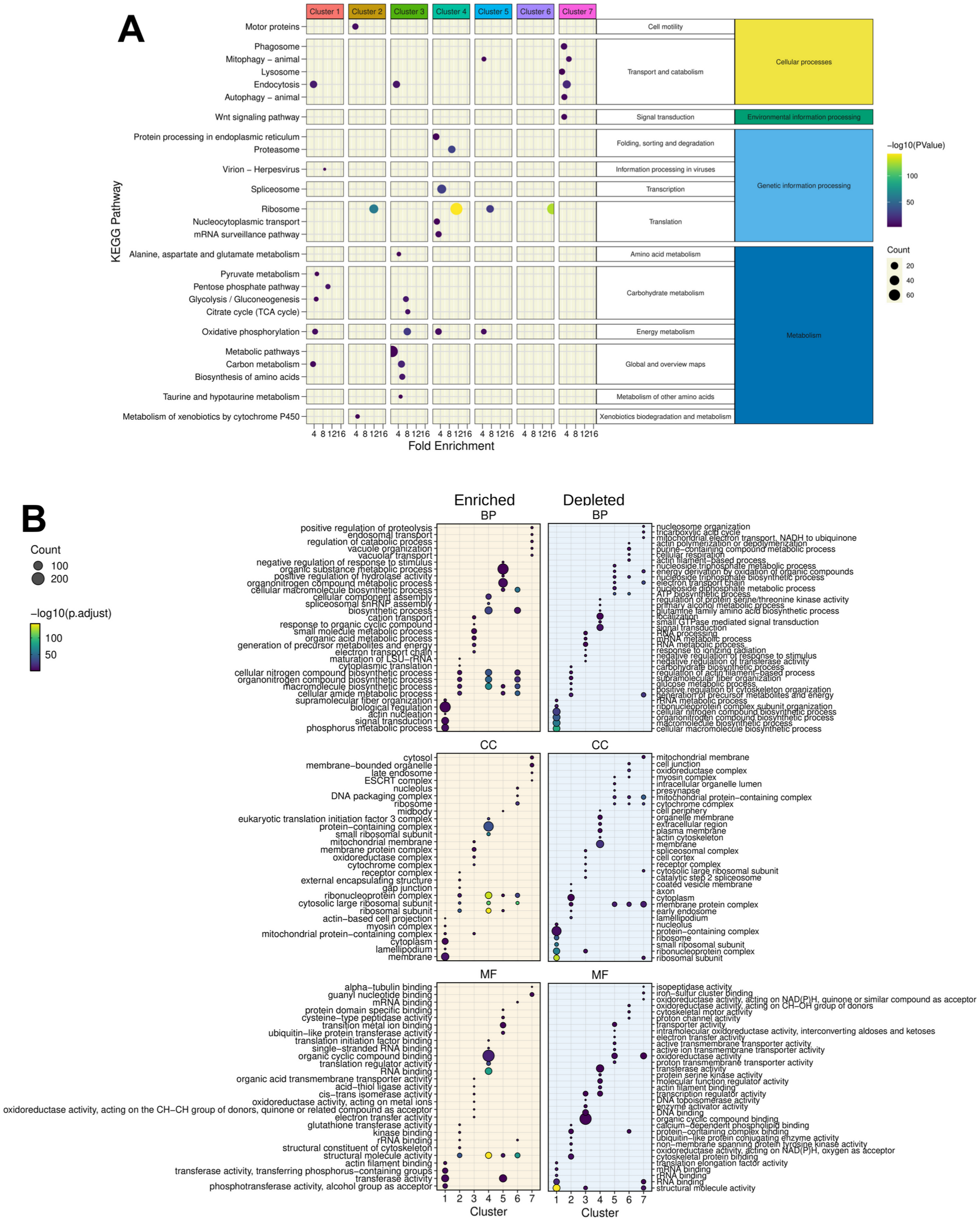
KEGG and Gene Ontology analysis of the gene signature in each cluster. **(A)** A synthetic representation of the KEGG pathway analysis is shown. Colored columns represent the 7 transcriptomic clusters. Each row is a KEGG pathway, the colored dot represents the - log(p-value) and the dot size represents the number (count) of enriched genes in each pathway category. The fold enrichment is shown on the x-axis. Panels **(B)** show the results of Gene Ontology terms (GO-terms) for Biological Processes (BP), Cellular Components (CC) and Molecular Functions (MF) respectively, obtained with the genes of each cluster (Absolute value Log2FC > 0.25 and significant p-value < 0.001) using RBGOA analysis (p-value <= 0.001) for three different ontology universes. Each panel corresponds to one ontology universe, and the analysis highlights enriched and depleted terms. The dot size indicate the proportion of significant GO-terms and the gradient scale the p-adjusted value.

For a more detailed functional characterization of each transcriptomic cluster, we performed a re-annotation of the *C. gigas* genome using a combination of tools to enhance the GO term richness of the existing annotation prior to functional GO term analysis. The original annotation for *C. gigas*, based on the work of Penaloza et al. (*30*), provided GO annotations for 18,750 out of 30,724 genes (61%). In order to improve upon these baseline annotations, we used the *C. gigas* genome (Genbank reference GCA_902806645.1) and the associated gff3 annotation file from the Roslin Institute (*30*). Using the Orson pipeline (see Materials and Methods), these files were used to extract and process the longest CDSs for GO-term annotation, and we then re-annotated each predicted protein by sequence homology, assigning putative functions and improving downstream GO-term analyses. Of the 30,724 extracted CDSs, 22,462 were annotated (GO-terms and sequence description), yielding an annotation percentage of 73.1 %. Of the 30,724 CDSs, 22,391 were annotated with Molecular Functions (MF), Biological Processes (BP), and Cellular Components (CC) GO-terms (**Supp. Fig. S2 and Supp. Data 2**). Overall, this improved annotation coverage significantly enhances the resolution of our downstream functional analyses. Using the GO-term annotation and the Log2FC of genes calculated after scRNA-seq processing in each cluster, GO enrichment analysis was performed by rank-based gene ontology analysis (RBGOA) (*33*).

RBGOA analysis was performed on the GO-terms identified in each cluster (**Supp. Data 3**). The results are presented in **Figure 2B**. The scRNA-seq-based analysis identified seven distinct transcriptomic profiles for each cell cluster, thereby shedding light on greater heterogeneity and functional diversity of *C. gigas* hemocytes than previously described. Cluster 1, comprising 27.6 % of cells, is characterized by GO-terms related to myosin complex, lamellipodium, membrane and actin cytoskeleton remodelling, as well as phosphotransferase activity. This is evidenced by an enrichment in actin nucleation, phosphorus metabolic process and signal transduction BP, actin filament binding and phosphotransferase activity MF and lamellipodium and actin-based cell projection CC. Cluster 2, comprising 23.1 % of cells, has an increased translation activity, as indicated by enrichment in BP, MF and CC terms related to rRNA binding, cytoplasmic translation, maturation of LSU-rRNA, and ribosomes respectively. Cluster 3, representing 17.8 % of cells, shows enrichment in cellular oxidation and actin nucleation, as evidenced by an enrichment in BP and MF related to oxidoreductase activities acting on metal ions, electron transfer and cellular oxidation, and CC related to cytochromes, oxidoreductase complex, and mitochondrial protein-containing complexes. Cluster 4, representing 16.9 % of cells, has transcriptomic signatures reminiscent of spliceosome assembly, RNA maturation, and macromolecules biosynthesis (BP related to biosynthetic process, macromolecules biosynthesis process, and spliceosome assembly, MF related to RNA and ssRNA binding and translation initiation, and CC related to ribosomes and ribonucleoprotein complexes). Cluster 5, comprising 7 % of cells, demonstrates enrichment in structural molecule activity, hydrolase regulation and ubiquitin-like protein transferase activity, as evidenced by its BP, MF, and CC related to ribosome and midbody localization. Cluster 6, comprising 4.6 % of cells, is characterized by a specific nucleosome organization, translation, and biosynthetic processes, with related BP, MF, and CC tied to nucleolus, ribosomes, ribonucleoprotein complex and rRNA binding. Cluster 7, representing a mere 3 % of cells, is characterized by multivesicular body sorting, vacuolar transport, vacuole organization, and positive regulation of proteolysis, with related BP, MF, and CC of the ESCRT machinery.

### Seven morphologically distinct immune cell types are identified by quantitative cytology and transcriptomic markers

Cytological studies were conducted using MCDH staining to better characterize the diversity of circulating hemocytes in *C. gigas*. Seven distinct hemocyte morphotypes were identified: three non-granular (acidophilic blasts, basophilic blasts and hyalinocytes) and four granular (small granule cells, big granule cells, vesicular cells and macrophage-like) **(Fig. 3 and Supp. Fig. S3)**. Hyalinocytes (30 % of total hemocytes) are large cells with an irregular spreading membrane. They contain an azurophilic cytoplasm without granulations and their irregular nucleus varies in size (**Fig. 3A panel H**). Basophilic blasts characterized by a basophil cytoplasm (**Fig. 3A panel BBL**) and acidophilic blasts with acidophil cytoplasm (**Fig. 3A panel ABL**) accounted for 17 % and 14 %, respectively, of the total hemocytes. They are rounded cells without granulation, with a uniform and regular dense nucleus and a high nucleo-cytoplasmic ratio. Macrophage-like cells (20 % of the hemocytes) present an irregular membrane punctuated by rare pseudopodia with a polylobed nucleus and a basophilic cytoplasm where polychromatic inclusions of various sizes could be observed (**Fig 3A. panel ML**). Small granule cells (13 % of total hemocytes) have an irregular membrane punctuated by rare pseudopodia, an acidophilic cytoplasm, and numerous homogeneous purple granules (**Fig. 3A panel SGC**). Big granule cells (4 % of total hemocytes) are rounded cells with a basophilic cytoplasm containing large, slightly dark purple to black vesicles of heterogeneous size (**Fig. 3A panel BGC**). Vesicular cells (2 % of total hemocytes) are rounded cells with acidophilic cytoplasm, rich in homogeneous transparent and fluorescent to UV light vesicles with an irregular nucleus (**Fig. 3A panel VC**). Only blasts (BBL, ABL) have a nucleo-cytoplasmic ratio greater than 0.6, whereas the other hemocytes described have a nucleo-cytoplasmic ratio less than 0.3.

**Fig 3.**
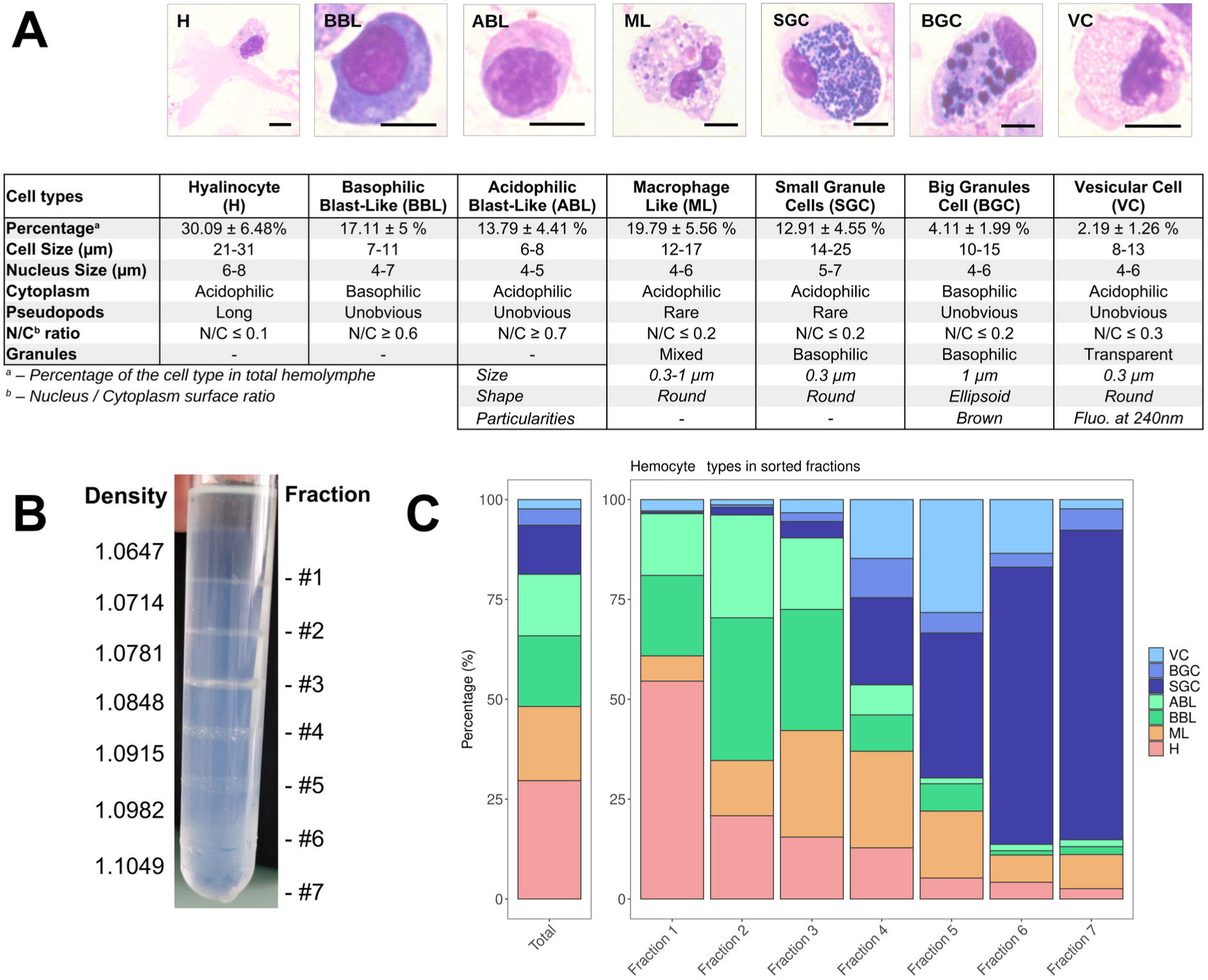
*C. gigas* naive hemocyte formula and Percoll gradient hemolymph fractionation. **(A)** Morphology, percentages and characteristics of the 7 cell types identified by MCDH staining. **H** : Hyalinocyte, **ML** : Macrophage Like, **BBL** : Basophilic Blast Like cell, **ABL** : Acidophilic Blast Like cell, **SGC** : Small Granule Cell, **BGC** : Big Granule Cell, **VC** : Vesicular Cell. <u>Scale bar :</u> 5 µm. For the ML, SGC, BGC, and VC hemocyte types, size refers to the average granule diameter, shape describes the morphology of the granules (e.g., round, ellipsoid), and particularities highlight distinguishing features such as granule color or fluorescence properties observed under specific staining or imaging conditions. **(B)** Sorting of hemocytes on a discontinuous Percoll gradient. 7 fractions were identified along the gradient at the top of each density cushion (from d=1.0647 at #1 to d= 1.1049 at #7). **(C)** Representation of the average values (from 5 different fractionation experiments) of the different hemocyte types in the seven percoll gradient fractions compared to the average hemolymph composition of a naive oyster (Total). **VC** : Vesicular Cells, **BGC** : Big Granule Cells, **SGC** : Small Granule Cells, **ABL** : Acidophilic Blast Like cells, **BBL** : Basophilic Blast Like cells, **ML** : Macrophage Like cells and **H** : Hyalinocytes respectively. (**Supp. Fig. S4** for statistics).

To gain further insight into the functional characterization of these diverse hemocyte morphotypes and to associate them with the transcriptomic clusters identified in the scRNA-seq, we enhanced an existing hemocyte fractionation approach using an isopycnic Percoll density gradient to sort the hemocytes (*34*). Cell sorting was performed using a discontinuous Percoll gradient with densities ranging from 1.0647 to 1.1049. Seven distinct density fractions were established (**Fig. 3B**) and the hemocyte composition was then characterized by cytological analysis of each fraction (**Fig. 3C**; see also **Supp. Fig. S4** for statistical significance). The uneven distribution of hemocyte morphotypes along the density gradient enabled a relative separation of the different cell populations (**Supp. Fig S4**). In summary, hyalinocytes were significantly enriched in the first fraction, while macrophage-like cells were significantly enriched in fraction 3 and fraction 4. Acidophilic blasts were significantly enriched in fraction 2. Basophilic blasts were significantly enriched in fractions 2 and 3. Vesicular cells were enriched in fraction 5, big granule cells were enriched in fraction 4 and small granule cells in fractions 6 and 7 (**Fig. 3C and Supp. Fig. S4**).

To identify the transcriptomic cell clusters corresponding to the different hemocyte morphotypes, we used RT-qPCR to detect the expression of different cluster-specific marker genes in the different hemocytes in the Percoll density fractions. The marker genes were selected without any a priori assumptions based solely on their enrichment levels (Log2C) and their percentage of expression (Pct1 / Pct2 ratio)F in the cluster of interest relative to all other cell clusters (**Table 2).** The expression level of each marker in the seven different clusters confirmed that the 14 selected marker genes were differentially expressed in each scRNA-seq transcriptomic cluster (**Supp. Fig. S5**).

**Table 2.** Table of the 14 marker genes specific to the different transcriptomic clusters. Gene number, average Log2FC, pct1/pct2 ratio (percentage of cells expressing this transcript in the cluster divided by the percentage of all other cells expressing this transcript) and cluster number are reported. The description is taken from our annotation and the marker name is derived from the description.

| Gene Number | Avg. Log2FC | Pct.1 | Pct.2 | Pct.1/Pct.2 Ratio | Cluster | Description | Marker Name |
| --- | --- | --- | --- | --- | --- | --- | --- |
| G8994 | 2.05 | 0.917 | 0.024 | 38.20 | 1 | Laccase-24 | LACC24 |
| G8841 | 3.13 | 0.988 | 0.049 | 20.16 |  | C-type lectin domain-containing protein | CLEC |
| G24846 | 2.31 | 0.914 | 0.029 | 31.52 | 2 | EGF-like domain-containing protein 8 | EGFL |
| G17277 | 3.21 | 0.989 | 0.043 | 23.00 |  | Putative modulator of levamisole receptor-1 | LEVAR |
| G22387 | 3.16 | 0.972 | 0.029 | 16.76 | 3 | TGc domain-containing protein | TGC |
| G21091 | 5.51 | 0.992 | 0.129 | 7.69 |  | X-box binding protein-like protein | XBOX |
| G35227 | 1.46 | 0.549 | 0.055 | 9.98 | 4 | ML domain-containing protein | MLDP |
| G31074 | 0.94 | 0.501 | 0.077 | 6.51 |  | High mobility group protein B1 | HMGB1 |
| G14441 | 3.35 | 0.837 | 0.076 | 11.01 | 5 | Cubilin | CUBN |
| G12472 | 3.99 | 0.847 | 0.034 | 24.91 |  | Galectin | GAL |
| G29512 | 3.20 | 0.985 | 0.004 | 246.25 | 6 | Caveolin | CAV |
| G3767 | 6.45 | 0.740 | 0.018 | 41.11 |  | Natterin-1 | NAT1 |
| G13140 | 3.12 | 1.000 | 0.025 | 40.00 | 7 | DBH-like monooxygenase protein 1 | MOX |
| G32859 | 3.90 | 0.929 | 0.039 | 23.82 |  | G protein receptor F1-2 domain-containing protein | GPROT |

RT-qPCR expression profiles in the hemocyte fractions obtained from the Percoll density gradient revealed distinct patterns according to the different transcriptomic cluster marker genes (**Fig. 4A**). Cluster 1 markers (LACC24 and CLEC) were overexpressed in fractions 1, 2, 3 and underexpressed in the remaining fractions relative to total hemocytes. Cluster 2 markers (EGFL and LEVAR) showed a decreasing pattern of expression from fraction 1 to fraction 7. Cluster 3 markers (TGC and XBOX) showed a significant increase in expression in fractions 4 to 7. Cluster 4 markers (MLDP and HMGB1) showed a gradually decreasing expression pattern from fraction 2 to fraction 7. Cluster 5 marker genes (GAL and CUBN) were underexpressed in fractions 4 to 7, but not differentially expressed in fractions 1 to 3 compared to total hemocytes. Cluster 6 marker genes (CAV and NAT1) were overexpressed in fraction 3, expressed similarly to total hemocytes in fraction 2, and underexpressed in fractions 1, and 4 to 7. Finally, cluster 7 marker genes (MOX and GPROT) were overexpressed only in fractions 3 to 7. Correlation analysis and statistical validation demonstrated a clear association between hemocyte morphotypes and 14 gene markers (**Fig. 4B**, **Fig. 4C and Supp. Table S4**). Cluster 3 marker genes (TGC and XBOX) correlated positively (r = 0.74 & 0.75) and specifically with small granule cells (SGC). Cluster 7 markers (MOX and GPROT) correlated (r = 0.68 and r = 0.56) with vesicular cells (VC). Cluster 2 markers (EGFL and LEVAR) correlated positively with hyalinocytes (H) (r=0.85 and r=0.81). No specific markers could be identified for blasts and macrophage-like cells.

**Fig 4.**
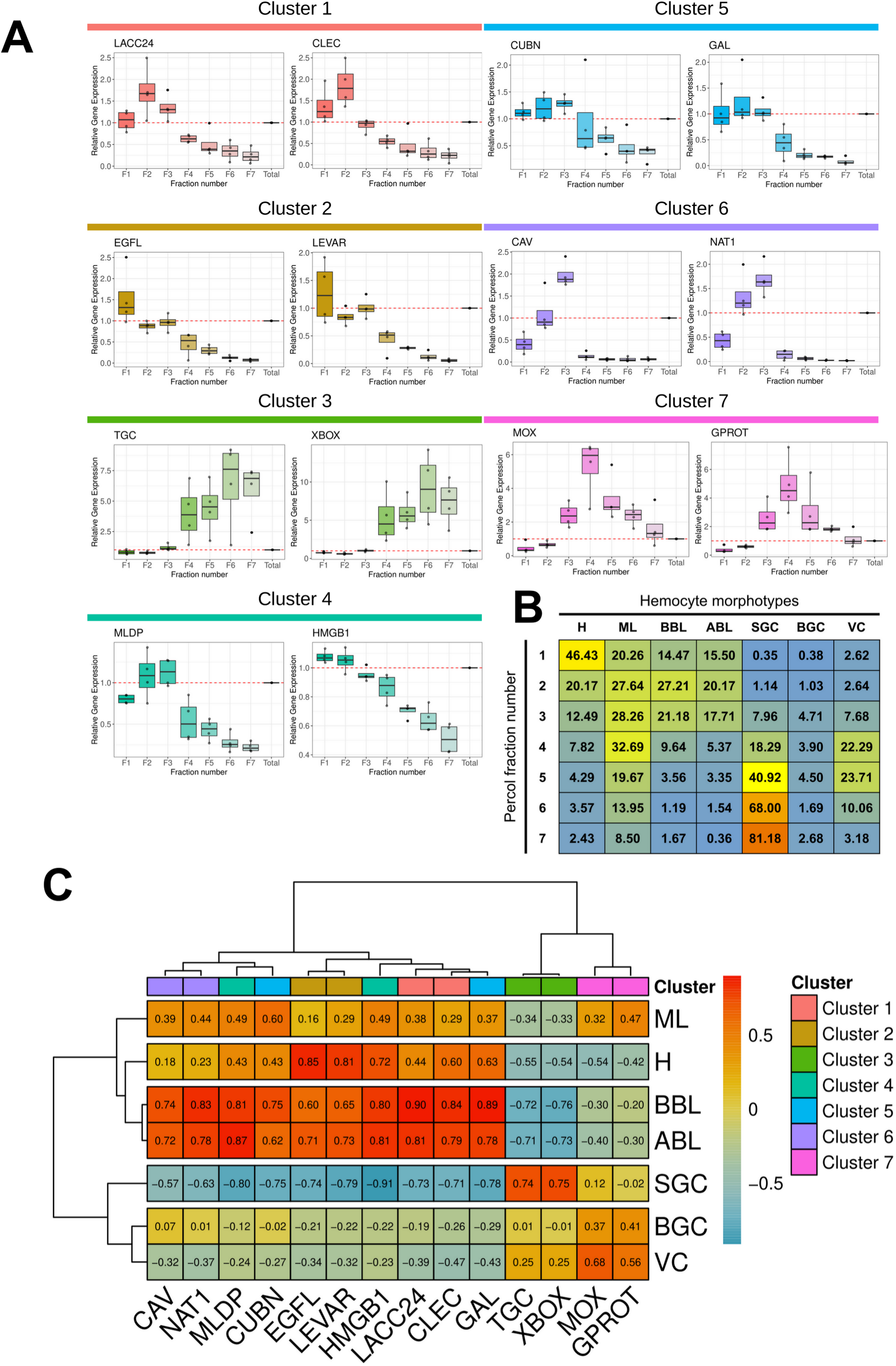
Characterization of molecular markers specific to the different hemocyte morphotypes. **(A)** Relative gene expression level of the 14 markers in the various fractions after gradient density sorting. The graphs show the relative expression of genes compared to their expression in total hemocytes in the various fractions (red dotted line). Relative gene expression levels were normalized to the reference gene Cg-rps6 and the 2^-ΔCt^ method was used to calculate relative expression levels, where ΔCt represents the difference between the target gene’s Ct value and the reference gene’s Ct value. Standard deviations were calculated based on four independent experiments. **(B)** Average percentage of each hemocyte type in the 7 Percoll gradient fractions used to quantify marker gene expression by qPCR. **(C)** Correlation matrix between the relative gene expression of each marker gene in each fraction and the percentage of each hemocyte type in the same fractions. Values and color scale represent the Pearson correlation coefficient (r) ranging from -1 (inverse correlation) to +1 (full correlation). **H** : Hyalinocyte, **ML** : Macrophage Like, **BBL** : Basophilic Blast Like cell, **ABL** : Acidophilic Blast Like cell, **SGC** : Small Granule Cell, **BGC** : Big Granule Cell, **VC** : Vesicular Cell. **LACC24** : Laccase 24, **CLEC** : C-type lectin domain-containing protein, **EGFL** : EGF-like domain-containing protein 8, **LEVAR** : Putative regulator of levamisole receptor-1, **TGC** : TGc domain-containing protein, **XBOX** : X-box binding protein-like protein, **MLDP** : ML domain containing protein, **HMGB1** : High mobility group protein B1, **CUBN** : Cubilin, **GAL** : Galectin, **CAV** : Caveolin, **NAT1** : Natterin-1, **MOX** : DBH-like monooxygenase protein 1, **GPROT** : G protein receptor F1-2 domain-containing protein.

By employing RT-qPCR, cell sorting on Percoll gradients, and scRNA-seq analysis, we could identify cell types corresponding to three transcriptomic clusters. Cluster 3 corresponds to small granule cells (SGC), cluster 2 to hyalinocytes (H), and cluster 7 to vesicular cells (VC). Cluster 3, cluster 2 and cluster 7 represent 17.8 %, 23 % and 3 % of the total cells analyzed by scRNA-seq, respectively. This is consistent with cytological data, which indicated the presence of 12.5 % +/-5 % Small Granule Cells, 30 % +/-9.3 % Hyalinocytes and 2.3 % +/-1.6 % Vesicular Cells (**Fig. 1B and Fig. 3A**). Furthermore, the molecular and cellular functions derived from GO-term and KEGG analyses (**Fig. 2**) were in alignment with the anticipated functions for these three cell types.

### Only macrophage-like and small granule cells behave as professional phagocytes

The hemocytes separated on the Percoll gradient were characterized functionally to gain further insight into their functional specialization. Two hemocyte functions, namely phagocytosis and production of Reactive Oxygen Species (ROS), are known to carry out major cellular antimicrobial activities. Granular cells have been suggested to be the professional phagocytes specialized for these functions (*22*). Phagocytosis and oxidative burst were studied using cell response toward zymosan particles (*35*).

The phagocytic activity of hemocytes was first tested on a sample of total oyster hemolymph. Cells were incubated for 1 hour with either zymosan particles or the bacterial strain *Vibrio tasmaniensis* LMG20012^T^. Only the small granule cells and the macrophage-like cells exhibited efficient phagocytosis for both zymosan or vibrios, as observed after MCDH staining (**Fig. 5A, Supp. Fig. 6 and Supp. Fig. 7A**). Macrophage-like cells and small granule cells showed a phagocytic activity of 49 % and 55 %, respectively, and a phagocytosis index of 3.5 and 5.2 particles per cell respectively (**Fig. 5B and Supp. Fig. 7B**), as confirmed in 3 independent experiments examining a total of 2,807 cells. Very limited phagocytic activity was observed for hyalinocytes (1.7 %), basophilic blasts (0.18 %), and big granule cells (2.7 %) with a phagocytosis index of 2, 1, and 2.5 particles per cell, respectively (**Fig. 5B and Supp. Fig. 7B**). These results confirmed that only small granule cells and macrophage-like cells behave as professional phagocytes demonstrating robust phagocytic activity.

**Fig 5.**
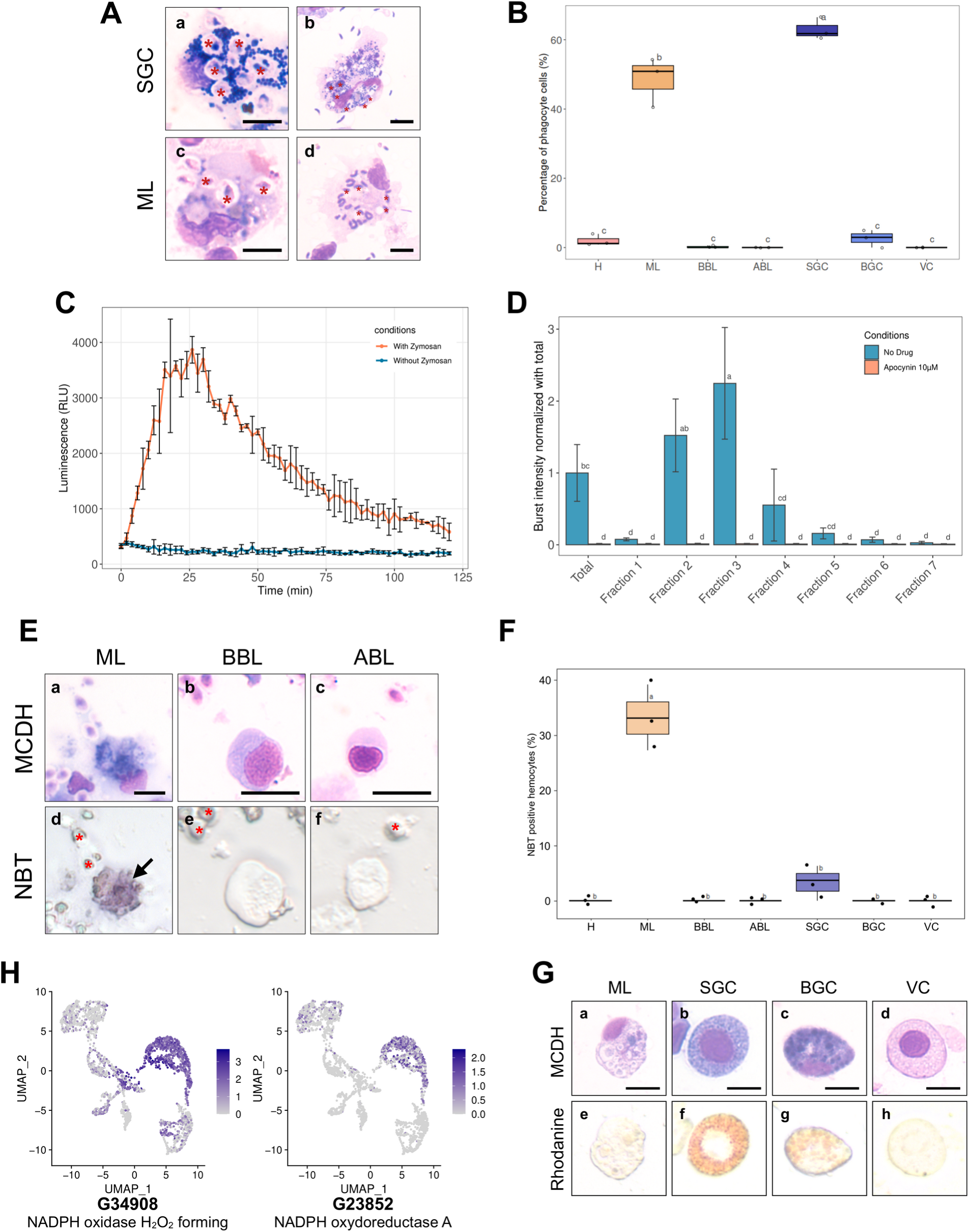
Phagocytosis, Reactive Oxygen Species production capacity and copper storage of hemocytes. **(A)** Images of small granule cells (**SGC**) and macrophage-like (**ML**) cells with phagocytosed zymosan particles (**panels a & c - red stars**) and *Vibrio tasmaniensis* LMG20012^T^ bacteria (**panels b & d - red stars**) from whole hemolymph sample. <u>Scale bar :</u> 5 µm. **(B)** Quantification of the phagocytic activity of zymosan particles by each cell type. The graph shows the result of 3 independent experiments. **(C)** Luminescence recording to detect the production of Reactive Oxygen Species (ROS). In orange, a biphasic curve was obtained on naive oyster hemolymph after zymosan addition at t = 0 min. In blue, the control condition corresponds to hemocytes without zymosan addition. **(D)** Graph showing the intensity of ROS production in each Percoll fraction. Normalized burst intensity was calculated from the luminescence peak obtained from each fraction. In blue, no drug was added to the experiment, in orange, ROS production was impaired by the addition of apocynin. **(E)** NBT (NitroBlueTetrazolium) staining of hemocytes exposed to zymosan particles. Hemocytes morphology after MCDH staining: Macrophage Like **(a**), Basophilic **(b)** and Acidophilic **(c)** Blast cells. NBT staining of the different types of hemocytes **(d-f)**. Red stars show zymosan and bacteria particles. Black arrows indicate Macrophage-Like cells. <u>Scale bar :</u> 10 µm **(F)** Quantification of NBT-positive cells present in the total hemolymph of oysters exposed to zymosan. **(H)** UMAP plots showing cells expressing NADPH oxidase found in the scRNA-seq dataset and their expression level. **(G)** Labeling of intracellular copper stores in *C.gigas* hemocytes. MCDH (upper panels) and rhodanine (lower panels) staining of oyster hemocytes to reveal copper accumulation. <u>Scale bar :</u> 10µm. For panels **(B)**, **(D)** and **(F)** the alphabetic characters displayed above the data points in each plot represent statistically significant differences among the groups, as determined by Tukey’s test following ANOVA. Groups denoted by different letters differ significantly at the p < 0.05 level of statistical significance. **H** : Hyalinocytes, **ABL** : Acidophilic Blast-Like cells, **BBL** : Basophilic Blast-Like cells, **ML** : Macrophage-Like cells, **SGC** : Small Granule Cells, **VC** : Vesicular Cells and **BGC** : Big Granule Cells.

### Only macrophage-like cells produce Reactive Oxygen Species

The next step was to assess the capacity of *C. gigas* hemocytes in each Percoll fraction to produce Reactive Oxygen Species (ROS) upon stimulation by zymosan, utilizing the Luminol oxidation assay. Luminol luminescence peaked 25 minutes after exposure of hemocytes (isolated from total hemolymph) to zymosan, indicating a robust oxidative burst (**Fig. 5C**). The production of ROS was dependent on a NADPH oxidase, as it was completely inhibited by the NADPH oxidase-specific inhibitor apocynin (**Fig. 5D**). To ascertain which hemocyte types were involved in ROS production we tested Percoll density-sorted hemocytes for oxidative burst activity. Fractions 2 and 3 displayed higher oxidative burst activity than the other fractions. The burst intensity of fraction 3 was twice that of total hemocytes, and fraction 2 also exhibited significantly increased burst activity. In contrast, fraction 4 showed a significant decrease in oxidative burst, and no activity was observed for fractions 1, 6 and 7 (**Fig. 5D**). These results indicate that NADPH-dependent oxidative burst activity is carried out by fractions enriched in macrophage-like (ML) cells and blast-like cells (ABL and BBL).

Previous studies have shown that granular cells can produce ROS (*22*). In contrast, we found that small granule cells collected in fraction 7 could not produce ROS through an oxidative burst. To identify which type of hemocyte from blast-like cells and macrophage-like cells produces ROS within a few minutes after exposure to zymosan, ROS production was investigated using NitroBlueTetrazolium (NBT) reduction to stain hemocytes directly. Correlative microscopic analysis using MCDH staining can then be conducted after the NBT reduction reaction. Macrophage-like cells were strongly and significantly stained by NBT reduction (33.2 %) (**Fig. 5E, panels a & d**). In contrast, some small granule cells were lightly stained by NBT reduction (**Fig. 5F**). Blast-like cells, big granule cells, vesicular cells, and hyalinocytes were never NBT stained, confirming that these cell types were not involved in ROS production (**Supp. Fig. S7**). Taken together, these observations demonstrate that small granule cells and macrophage-like cells are the two professional phagocytes among hemocytes. However, only macrophage-like cells are capable of oxidative burst upon exposure to zymosan. In light of these new functional data, we further analyzed the expression level of NADPH-oxidase-related enzymes in the scRNA-seq dataset. The cells in cluster 1 predominantly expressed two NADPH oxidase isoforms (gene numbers G34908 and G23852) compared to other clusters (**Fig. 5H**). Furthermore, this cluster expresses macrophage-related genes, including the macrophage-expressed gene 1 protein (G30226) (**Supp. Data S1**), along with maturation factors for dual oxidase, an enzyme involved in peroxide formation (**Supp. Fig. S8**), supporting its designation as macrophage-like based on functional characteristics. Collectively, these data indicated that cells of transcriptomic cluster 1 corresponded to macrophage-like cells.

### Small granule cells and big granule cells accumulate intracellular copper

The effects of copper on oyster hemocytes have been studied extensively due to its abundance in polluted marine environments (*19*). In addition, several studies have shown that *Vibrio* species pathogenic for *C. gigas* possess copper resistance genes, which are crucial for their survival within hemocytes (*9*). This prompted us to investigate which hemocyte types are involved in copper metabolism. To this end, total hemocytes were isolated from naive oysters and stained with rhodanine to reveal copper storage in cells, and a total of 1,562 cells were examined across three independent experiments. Rhodanine staining revealed that 33 % +/-2 % of small granule cells and 30 % +/-10 % of big granule cells exhibited a specific reddish/brown staining indicative of a high concentration of copper in their granules (**Fig. 5G and Supp. Fig. S9**). These results provide functional evidence that small granule cells (SGCs) are specialized in metal homeostasis in addition to phagocytosis, as suggested by the scRNA-seq data identifying cluster 3. Specifically, single-cell RNA sequencing revealed an upregulation of copper transport-related genes, including G4790 (a copper transporter) with a 2.7 log2FC and a pct ratio of 42, reinforcing the role of SGCs in copper homeostasis (see Supp. File S1).

### Antimicrobial peptides are expressed by agranular cells, blasts and hyalinocytes

Antimicrobial peptides (AMPs) have long been studied for their role in the invertebrate humoral immune response. They are expressed by hemocytes, including those in *C. gigas* (*36*). The aim was to ascertain whether different hemocyte cell types expressed distinct AMPs. However, due to the limited sequencing depth, the scRNA-seq analysis was not sensitive enough to reveal AMP expression. This limitation was addressed by investigating AMP expression through RT-qPCR on Percoll density-fractionated hemocytes. The results indicated that Cg-Bigdefs 1-2, Cg-BPI and hemocyte defensin were predominantly expressed by agranular cells, blasts and hyalinocytes (ABL, BBL and H) **(Fig. 6A, B and C)**. The expression of these AMPs was associated with Blasts abundance, while the expression of Cg-BigDefs 1-2 was only associated with hyalinocytes **(Fig. 6D)**. Granular cells (VC, ML, BGC and SGC) did not seem to express any of the analyzed AMPs. These data suggest that some of the agranular cells are specialized in the production of humoral effectors.

**Fig 6.**
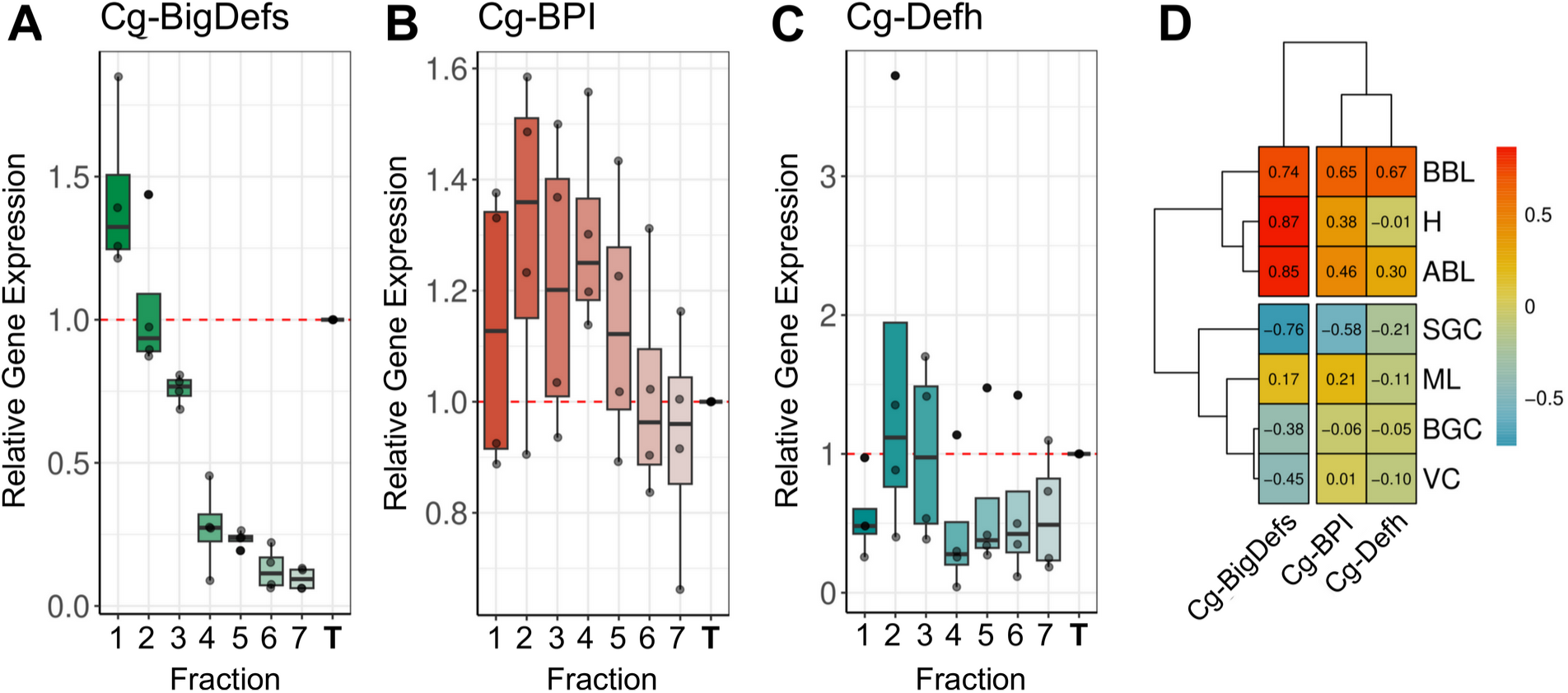
Hemocyte expression profiles of some antimicrobial peptides. **(A) (B)** and **(C)** Relative gene expression in the 7 Percoll hemocyte fractions of Big-Defensin1 & 2 (Cg-BigDefs), BPI (Cg-BPI) and hemocyte defensin (Cg-Defh), respectively, in comparison to the gene expression level in unfractionated hemolymph. Relative gene expression levels were normalized to the reference gene Cg-rps6. The 2^-ΔCt^ method was used to calculate relative expression levels, where ΔCt represents the difference between the target gene’s Ct value and the reference gene’s Ct value. **(D)** Correlation matrix between the relative gene expression of BigDefensin1 & 2, BPI and hemocyte defensin gene in each fraction and the percentage of each hemocyte type in each fraction (**H** : Hyalinocytes, **ABL** : Acidophilic Blast Like, **BBL** : Basophilic Blast Like, **SGC** : Small Granule Cell, **ML** : Macrophage Like, **BGC** : Big Granule Cell, **VC** : Vesicular Cell. Values and color scale represent the Pearson correlation coefficient (r) ranging from -1 (inverse correlation) to +1 (full correlation).

### Tentative model of hemocyte lineages and differentiation pathways in *C. gigas*

The ontogeny, lineage and differentiation pathways of bivalves remain largely unknown (*37*). However, there are some indications of circulating and proliferating hemocyte progenitors in the hemolymph of *C. gigas* (*38*). GO-terms analysis of the 7 transcriptomic clusters revealed different functional signatures, including the transcriptomic signature of cluster 4, which showed a high expression level of ribosomal proteins (**Supp. Fig. S1J**) . This particularity has been observed in hematopoietic stem cells in vertebrates (*39–41*). Furthermore, scRNA-seq approaches using bioinformatic tools like Monocle3 (*42*) can now be used to deduce differentiation pathways from trajectory inference using gene expression profiles analysis. This revealed an overrepresentation of functions involved in the splicing, transcription and translation continuum in the same fourth cluster (see cluster 4 in **Fig. 2A** and **Fig. 2B**), thereby supporting the hypothesis of a pool of quiescent or immature cells that can differentiate upon stimulation.

Cluster 4 was chosen to enroot the pseudotime analysis to deduce differentiation pathways and cell lineages using Monocle3. (**Fig. 7A**). By temporally ordering the 2817 cells analyzed by scRNA-seq, 6 cell lineages could be defined (**Fig. 7B**). Differentiation pathway 1 leads to hyalinocytes (H) (**Fig. 7C**). This transition is characterized by the downregulation of 11 genes (**Supp. Fig. S10A**), two of which are transcription factors (G31522 and G7003). The hyalinocyte cluster was also characterized by the overexpression of genes involved in cell contractility (G11418, G22824, G153 and G718). Differentiation pathway 2 leads to cells of cluster 5 and is characterized by the upregulation of about 26 genes (**Supp. Fig. S10B**), one gene related to metalloendopeptidase activity (G15477) was downregulated, a GTPase IMAP gene (G29323) and three transcription factors were upregulated (G7003, G11013 and G4646) (**Fig. 7D**). Differentiation pathway 3 leads to cells of cluster 6 and is characterized by the upregulation of 22 genes (**Supp. Fig. S10C**). Three fibrinogen-related protein genes (G20176, G20226 and G20227) and a GATA-binding factor (G31054) were up-regulated whereas a metalloendopeptidase gene (G15477) was downregulated. The severin gene displayed a transient increased expression (G28864) (**Fig. 7E**). The pathways between clusters 4, 5 and 6 found by Monocle3 analysis were pseudo temporally short and few specific markers were identified, suggesting that they were transcriptionally close. Differentiation pathway 4 leads to vesicular cells (VC) (**Fig. 7F**) where a large number of genes were activated to give rise to these cells (**Supp. Fig. S10D**). Four genes were specifically downexpressed in VC (G15477, G25512, G2213 and G29172). Interestingly, 6 genes encoding potential transcription factors were upregulated in this lineage (G30997, G10637, G1067, G13555, G2123 and G27827). Differentiation pathway 5 leads to macrophage-like cells (ML), and is characterized by the underexpression of 41 genes and the overexpression of 20 genes (**Supp. Fig. S10E**). Among the 41 genes, 6 are putative transcription factors (G2123, G10637, G13555, G27827, G1067 and G11196) (**Fig. 7G**). Finally, we can outline a lineage ending in small granule cells (SGC) (**Fig. 7H**). This pathway involved cluster 4 (immature cells), VC, ML and SGC and was characterized by the downregulation of 27 genes (**Supp. Fig. S10F**), including 5 potential transcription factors (G10637, G13555, G27827, G1067 and G2123). SGC showed a distinct transcriptomic signature with 61 overexpressed genes, including 3 transcription factors (G1708, G21091 and G30622). Based on these findings, we postulate that immature cluster 4 cells possess pluripotent potential and can give rise to four terminally differentiated cell types : cluster 5 and 6 cells, hyalinocytes and small granule cells, and two other transient hemocyte cell types : vesicular cells and macrophage-like cells. All data from the Monocle3 analyses are available as Supplementary Data (**Supp. Fig. S10**).

**Figure 7.**
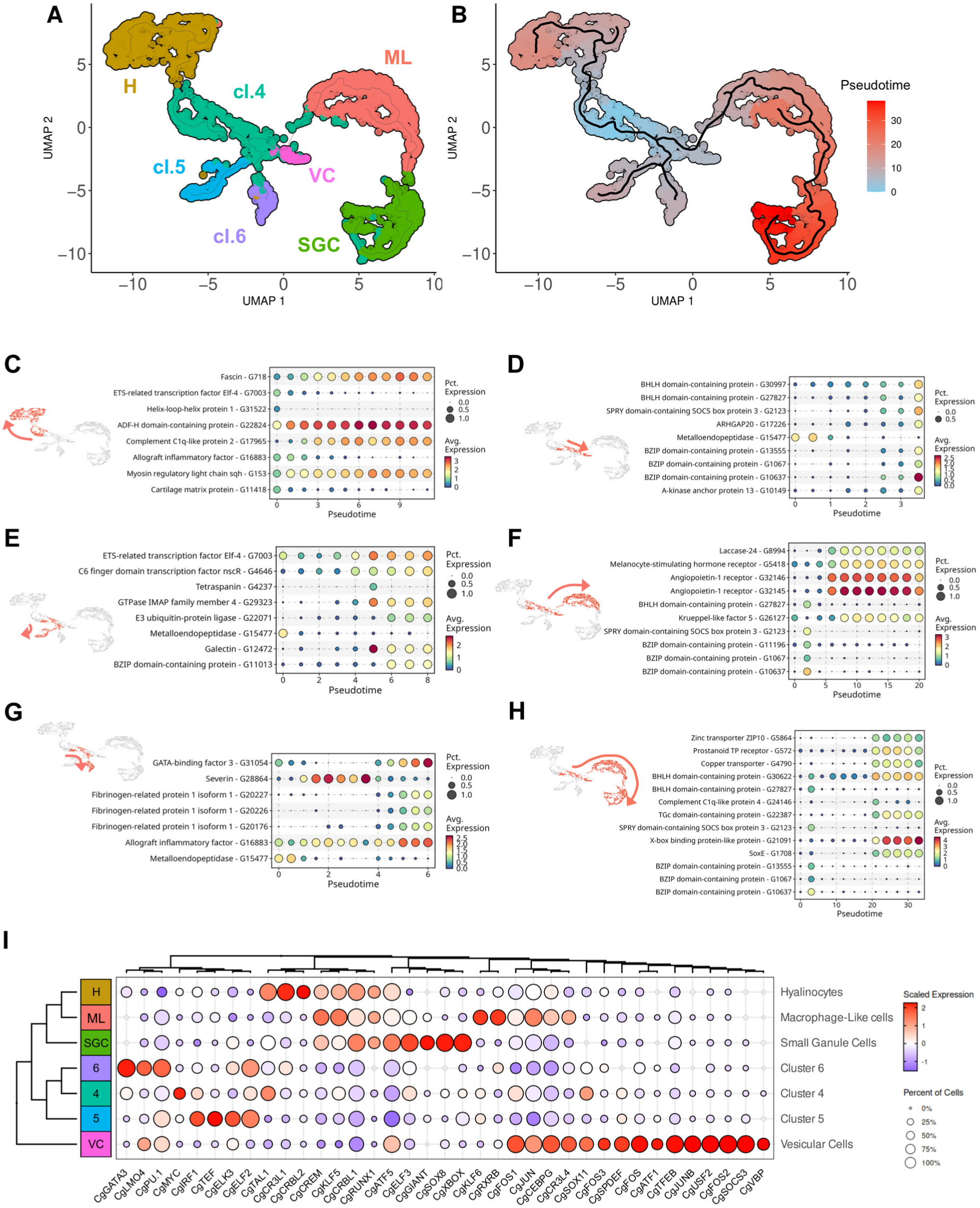
Pseudotime ordering of cells revealed 6 potential differentiation pathways of hemocytes. **(A)** UMAP plot of scRNA-seq analysis showing the 7 transcriptomic clusters used for pseudotime analysis. 4 clusters were identified cytologically (SGC for small granule cells - cluster 3, H for hyalinocytes - cluster 2, ML for Macrophage Like - cluster 1 and VC for vesicular cells - cluster 7), cl.4, cl.5, and cl.6 represent clusters 4, 5, and 6, respectively. **(B)** Graphical representation (UMAP projection) of the Monocle 3 pseudo-time order of the clustered cells. Cluster 4 (cl.4) was used as the origin for the pseudotime analysis. **(C) (D) (E) (F) (G) and (H)** show the gene expression level of selected marker genes obtained from the monocle3 trajectory analysis at the beginning and end of the modelized differentiation pathways (in red on the UMAP plot) from cluster 4 to hyalinocytes, to Vesicular Cells (VC), to cluster 5 cells, to Macrophage-Like cells (ML), to cluster 6 cells, and to Small Granule Cells (SGC) respectively. The color scale represents the normalized expression level of each gene. **(I)** Dot plot showing the average expression and the percentage of cells expressing identified transcripts encoding for transcription factors in the scRNA-seq dataset. The scale represents the normalized expression level, while the dot diameter indicates the percentage of cells expressing the gene.

Differentiation pathway analysis thus revealed the over- or under-expression of various transcription factors in the identified pathways. Given the established role of transcription factors as master regulators of cell differentiation and their utility in delineating cell lineages, we investigated the combinatorial expression patterns of transcription factors among the different transcriptomic cell clusters. Based on GO-terms annotation, 28 different sequences corresponding to transcription factors were isolated from the scRNA-seq dataset. The transcription factor function was confirmed by manual annotation (**Supp. Table S5**) and **Figure 7I** shows the average expression profiles of these factors in the different clusters. **Supplementary Figure S11** illustrates their expression levels in single cells. Two transcription factors, CgATF5 and CgCRBL1, exhibited a contrasting expression profile with an increased average expression in macrophage-like cells, hyalinocytes and small granule cells versus a low expression profile in cells in clusters 4, 5, 6 and vesicular cells. We also identified transcription factors that were specifically expressed in the different transcriptomic clusters : CgSPDEF, CgSOCS3, CgFOS and CgTFEB were specific for vesicular cells, CgSOX8, CgXBOX and CgELF3 were specific for small granule cells, CgCR3L1 and CgTAL1 for hyalinocytes, and CgJUN, CgKLF5, CgKLF6 and CgCREM for macrophage-like cells. Eight additional transcription factors were specifically identified in cluster 6 (CgGATA3, CgPU.1 and CgELF2), cluster 5 (CgELF2, CgELK3 and CgIRF1) and cluster 4 (CgTAL1 and CgFOS1). These data potentially define four distinct hematopoietic lineages originating from one type of immature blast cells and give rise to hyalinocytes, SGC (via VC and ML), or two distinct differentiated blast-like cells. We also identified a combination of transcription factors specific to lineages and cell types that are potential master regulators of cell fate during hematopoiesis.

## Discussion

The findings of our study represent a significant advancement in our understanding of the functional diversity and lineages of *C. gigas* hemocytes. Single-cell RNA-seq and cytology were combined to identify seven distinct hemocyte transcriptomic cell clusters and an equivalent number of morphotypes. These include four granular cell types (big granule cells, macrophage-like cells, small granule cells and vesicular cells), two distinct blast-like cells (basophilic and acidophilic blast-like cells), and one agranular epithelial-like cell type (hyalinocytes). A significant challenge was to identify correlations between transcriptomic and cytological data to fully define each cytological cell type. This challenge was overcome by combining multiple approaches, including isopycnic Percoll density gradient fractionation combined with the analysis of transcriptomic markers expression, and functional assays including phagocytosis, oxidative burst, copper accumulation, AMP expression and finally pseudotime analysis of gene expression. These results confirmed the historical classification of the 3 main cell groups : blasts, hyalinocytes, and granular cells (*15*) and deepened our understanding of the functional specificities of poorly characterized cell types. In particular, we identified distinct transcriptional and functional subtypes among blasts and granular cells with complementary immune specialization and lineage relationships between cells.

One significant outcome of the present study is the identification of cell types involved in antimicrobial activities, including phagocytosis, intracellular copper accumulation, an oxidative burst and antimicrobial peptide production. These cell types have been extensively studied for their role in antibacterial and antiparasitic defenses, as they are found in the large majority of invertebrates (*17*). Nevertheless, there has been considerable debate surrounding the cell types specialized for these critical immune functions in bivalves, particularly oysters. Moreover, the involvement of the different hemocyte subpopulations in immune functions is not yet fully understood.

Our findings reveal that the macrophage-like (ML) cells and small granule cells (SGC) are the sole hemocyte cell types that function as professional phagocytes, as demonstrated against Zymosan or *Vibrio*. These two distinct cell types could be distinguished functionally. First, two distinct transcriptomic clusters were identified for each cell type (Cluster 1 for ML / BGC and Cluster 3 for SGC). Secondly, only ML induced a measurable oxidative burst. Thirdly, only SGC accumulated intracellular copper in specific granules. Interestingly, our scRNA-seq analysis indicates that SGC (cluster 3) expresses the scavenger receptor cysteine-rich (SRCR) gene G3876, annotated as an Low-density lipoprotein receptor-related protein with a Log2 fold change (Log2FC) of 0.77 linking them to scavenger receptor-mediated pathogen recognition and clearance. This aligns with findings by Wang et al. (*43*), who demonstrated significant expansion and dynamic regulation of SRCR genes in response to pathogen-associated molecular patterns. The pseudotime trajectory linking macrophage-like (ML) and small granule-like (SGC) cells suggests a potential developmental relationship within the granular cell lineage; however, this hypothesis requires further validation. The notion that ML could serve as a precursor for SGC may seem counterintuitive. However, this is not an isolated phenomenon among invertebrate hemocytes. For instance, in *Drosophila* larvae (*44*), some populations of professional phagocytes, the sessile plasmatocytes, give rise to crystal cells or lamellocytes that are morphologically and functionally distinct from plasmatocytes (*45*). Similarly, the existence of multiple professional phagocytes in the oyster is reminiscent of vertebrate macrophages, polynuclear neutrophils and dendritic cells, which possess distinct functional specializations, including efferocytosis (*46*), oxidative burst, NETosis, or antigen presentation (*47*).

The characterization of professional phagocytes in oysters is of particular importance for a deeper understanding of oyster-*Vibrio* interactions during pathogenesis. Some of the most extensively studied oyster pathogens, including strains of *V. tasmaniensis* and *V. aestuarianus francensis,* harbor virulence traits that enable them to disrupt the phagocytic activity of hemocytes during pathogenesis. For instance, *V. tasmaniensis* behaves as a facultative intracellular pathogen with phagocytosis-dependent cytotoxicity (*11*). Additionally, both *V. tasmaniensis* and *V. aestuarianus* display resistance to copper toxicity through CopA and CusA/B transporters. This trait is essential for the survival and virulence of these pathogens in oysters (*9*, *48*). Since SGCs have been demonstrated to be professional phagocytes with copper-rich granules, the cellular interactions between SGCs and these vibrios are likely to be critical during the antibacterial host response and pathogenesis. ML are phagocytes that possess a very potent NADPH-oxidase-dependent oxidative burst. The oxidative burst is a rapid and potent antimicrobial response observed in professional phagocytes, such as polynuclear neutrophils in mammals. It is worth noting that NADPH-dependent ETosis has been observed in *C. gigas* in a manner analogous to that observed in human neutrophils (*49*). This cell death is characterized by the projection of DNA extracellular traps that capture and kill some pathogens like *Vibrio* (*49*). Therefore, it is reasonable to hypothesize that ML may also be involved in ETosis.

The specialized functions of the two other types of granular cells, the BGCs and the VCs, remain unclear. Despite the difficulty in identifying a specific scRNAseq transcriptomic cluster for BGCs, the level of expression of laccase 24 was found to be higher in a particular subcluster among ML (**Supp. Fig. S8**) and pseudo-time analysis highlighted the same subcluster as an alternative differentiation state among ML (**Fig. 7B**). The enrichment of transcripts involved in oxido-reduction pathways, particularly laccase 24, aligns with their potential role in melanization and response to oxidative stress (*50*). While melanin-like deposits have been observed in some cases of infestation by the parasites *Martelia* or B*onamia* in the Sydney rock oyster (*51*), this mechanism is not as robust as that described in arthropods, which perform melanization through a prophenoloxidase activation cascade (*52*, *53*). In many marine invertebrates (*50*), a type of hemocyte known as Brown Cells could be related to the BGCs described here. When observed without any staining (as in **Fig. 5F**), their big granules with a yellow to dark brown content appear to align with the historical description of brown cells that often infiltrate tissues (like gills) in animals exposed to polluted waters (*50*). It has therefore been theorized that these cells are involved in detoxification processes. Our pseudotime analysis indicates that they likely originate from ML, with limited phagocytosis activity and a specialized role in melanization and potentially heavy metal detoxification, as evidenced by rhodanine staining showing copper accumulation in some of their granules (**Fig. 5G**). Further studies are recommended to clarify the role of BGCs, particularly in the context of parasitic infestation or exposure to toxic stresses. Lastly, VC are granular cells that remain to be functionally characterized. Their transcriptomic profile suggests strong intracellular vesicular trafficking and autophagy activity. However, our functional assays did not reveal any particular immune-related function. Their clear granules appear auto-fluorescent when illuminated with UV light (**Supp. Fig. S12**) but the biochemical nature of the content of these granules remains to be characterized. As they also appear as cell intermediates along the granular cell differentiation pathway in pseudotime analysis, they could represent functionally immature precursors of the other three granular cell types, much like promyelocytes which possess specific azurophilic granules but are functionally immature precursors during granulocyte differentiation in humans. However, the enrichment of autophagy-related transcripts in VC calls for further investigation into a potential role in antiviral immunity, as autophagy has been suggested to play a role in the response to OsHV-1 virus (*54*).

Hyalinocytes are a homogenous cell type, with only one morphotype matching one transcriptomic cluster. It can be deduced from pseudotime analysis that they originate from a specific and very different differentiation pathway than the granular cells. According to the literature, hyalinocytes are involved in the early stages of inflammation and can infiltrate wounds and interact with foreign particles. In the flat oyster *Ostrea edulis*, they contribute to shell production and wound healing (*55*). In the Sydney rock oyster they play a role in cell aggregation (*56*), while in *C. virginica* (*57*) they contribute to encapsulation, reminiscent of lamellocytes in *Drosophila*. Our results suggest an important role of AMP expression in the immune response. Indeed, *Cg-*BigDefs, which participate in the control of oyster microbiota (*58*), were found to be expressed in both hyalinocytes and blast-like cells. Moreover, hyalinocytes from the oyster *O. edulis* have been shown to express the AMP Myticin C (*59*), which lends further support to this immune function. Among the AMPs, we also found that Cg-BPI and *Cg-*Defh, are more expressed in BBL, ABL than in other cell types. These results are somewhat unexpected, given the prevailing assumption that AMPs are stored in granules of granular cells, rather than agranular cells (*16*). These results highlight the necessity to reassess the role of specific agranular cell types in the active production of humoral immune effectors. Nevertheless, further staining approaches would be necessary to confirm these transcriptomic results. Our findings suggest that hyalinocytes and/or the blast-like cells may be a cellular target of the OsHV-1 virus, the causal agent of POMS, which dampens the expression of certain AMPs (*7*), thereby inducing bacterial dysbiosis. It is still unclear whether this is due to a decreased expression of AMPs and/or inhibition of immature blast cell differentiation involved in the renewal of agranular cell types.

It should be noted that the complexity of blast-like cells could not be fully elucidated in this study, as 3 clusters and only 2 morphotypes were identified. Cell fractionation using Percoll gradient failed to yield pure blast-enriched fractions (ABL and BBL), preventing precise functional characterization. The enrichment in transcripts of the transcription synthesis degradation continuum aligns with the definition of undifferentiated blast-type cells and with the basophilic staining obtained in MCDH for BBL cells, as immature blast cells are characterized by a basophilic cytoplasmic staining in humans. However, our results show that certain blast populations can produce AMPs, suggesting that these cells may also play a role in the production of humoral effectors. Ultimately, it remains unclear whether these circulating immature cells are indeed the stem cells from which all hemocytes originate.

In the animal kingdom, the innate immune system relies on specialized cells derived from pluripotent precursors through hematopoiesis. Transcription factors in particular have been found to exhibit a high degree of conservation throughout the animal kingdom, from invertebrates to vertebrates. However, the mechanisms underlying the functional differentiation of bivalves are only partially available or understood (*60*). Recent research has indicated the potential existence of hemocyte progenitors, also known as blast-like cells in several bivalve species. These include clams, mussels, scallops, marine mussels, freshwater mussels, oysters, pearl oysters, and wing shells (*60*). However, various models for hematopoiesis in bivalves have been proposed and extensively debated without a clear consensus or definitive proof. Single-cell RNA sequencing and pseudotime analysis have enabled us to propose a refined model of hematopoiesis at an unprecedented level of detail. In this model, the different types of hemocytes are likely produced through four differentiation pathways that originate from a common progenitor. One pathway results in the formation of agranular hyalinocytes, while another independent pathway gives rise to the granular cells, including VC, ML, BGC, and SGC. Furthermore, two additional differentiation pathways have been identified that may lead to terminally differentiated blast-like cells. The differentiation of mature hemocytes involves the establishment of lineage-specific gene expression profiles, which rely on transcription factors to modulate the expression of their target genes. Our study identified a combination of transcription factors that are differentially expressed during *C. gigas* hematopoiesis and are specific to differentiation stages and cell lineages, including many well-known hematopoietic transcription factors such as GATA, PU-1, TAL1 and SOX factors, which are positioned in this model of *C. gigas* hematopoiesis (*61–63*) (**Fig. 8**). Nevertheless, we did not detect any canonical stem or progenitor cell populations in our dataset, underscoring the need for future investigations - potentially involving immunological challenges and lineage-tracing assays - to clarify whether proliferative cells circulate in the hemolymph or instead reside primarily in tissue compartments.

**Figure 8.**
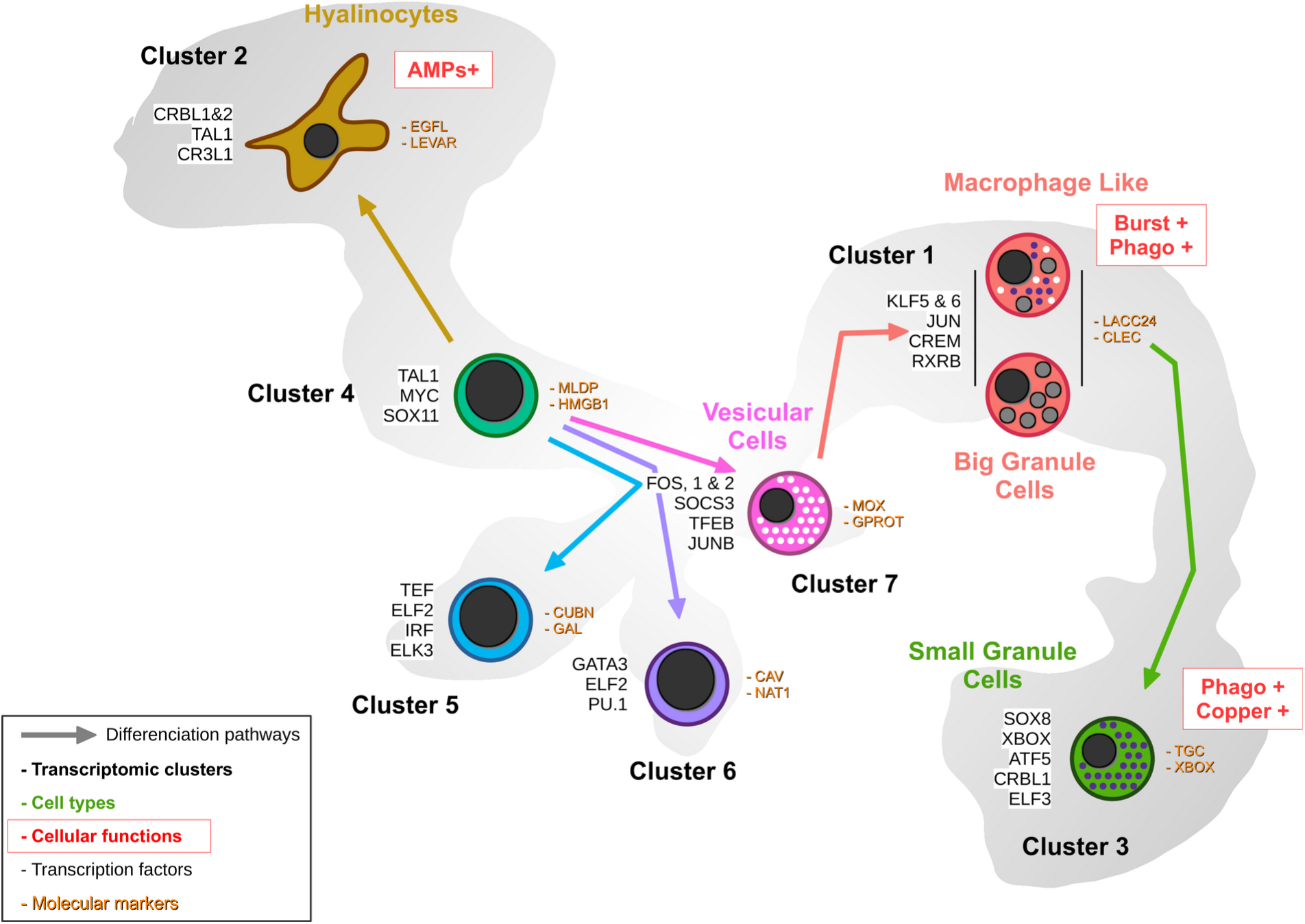
Proposed hemocyte ontology in *Crassostrea gigas* based on the transcriptomic, cytological and functional results obtained. Cells are colored according to the same color code as the transcriptomic clusters. Cluster numbers and cell types are indicated. To the left of the cells are the overexpressed transcription factors and to the right are the identified marker genes in each cluster. Functional characteristics of hyalinocytes, macrophage-like cells and small granule cells are marked in red. (**AMP** : AntiMicrobial Peptide, **Burst** : ROS production, **Phago** : phagocytosis)

The comparison of hemocyte populations in *C. hongkongensis* and *C. gigas* highlights a shared reliance on granulocytes as primary immune effectors, with *C. hongkongensis* (*27*). Our study emphasizes functional specialization in *C. gigas*, focusing on phagocytosis, oxidative stress responses, and vesicular trafficking, likely reflecting differences in transcriptomic and functional characterization methodologies. Moreover, the identification of conserved transcription factors such as Tal1, Sox, Runx and GATA in both datasets underscores the evolutionary conservation of hematopoiesis pathways across metazoans. While direct comparisons between scRNA-seq datasets are hindered by differing annotation frameworks, parallels can still be drawn with studies in other invertebrates. For example, shrimp and crayfish scRNAseq datasets (*64*) describe hemocytes with immune-activated states and integrin-related markers, respectively, while *Anopheles gambiae* (*65*)exhibits clusters with high ribosomal activity and transcription factors such as bHLH and myc. These findings highlight both the shared molecular foundations and species-specific differences that define hemocyte diversity and specialization.

This study significantly advances our understanding of bivalve immunity, especially in comparison to arthropods. By introducing a standardized reference hemocytogram for oysters using MCDH staining, defining cell type-specific markers and key transcription factors likely involved in cell fate determination, as well as clarifying the functions of different hemocytes, we have paved the way for future in-depth studies. Moreover, the identified clusters represent functionally and transcriptionally distinct cell populations under the conditions studied, yet they may reflect dynamic states rather than fixed subtypes, thereby acknowledging the potential for hemocyte populations to transition between states in response to environmental or physiological changes—even under homeostatic conditions. This will facilitate further studies of the oyster’s immune response to various biotic and abiotic stress at the cellular level. Improved comprehension of antiviral and antibacterial responses in bivalves, along with an enhanced understanding of immune priming and immune memory at the cellular level in bivalves, will benefit health and population management practices for sustainable aquaculture production. These findings will also contribute to the broader field of evolutionary immunology by enabling comparative studies and elucidating the diversification of immune cells and immunity-related genes in a protostome.

## Materials and Methods

### Conservation of oysters

The work described here was performed using two different sources of oysters of the same species *Crassostrea (Magallana) gigas*. ISA (Ifremer Standardized Oysters - La tremblade - France) oysters for the scRNA-seq experiment and oysters provided by a local supplier (https://www.huitres-bouzigues.com). Animals were washed and kept in 10 L tanks containing seawater and a bubbler to oxygenate and homogenize the water. Water was changed daily and oyster health was monitored. All animals used in this study were 18 months of age.

### Hemocyte collection and processing

*Crassostrea gigas* hemocytes were collected from live animals by puncture of the adductor muscle. The oyster shell was incised on the posterior side with forceps. Hemolymph was collected using a 23Gx1” needle mounted on a 5 mL syringe prefilled with 2 mL of ice-cold Alsever modified medium (20.8 g glucose – 8 g trisodium citrate - 22.5 g sodium chloride - 0.4 g BSA - pH=7.5). Samples were centrifuged for 4 minutes at 200 g – 4 °C and the supernatant was removed and replaced with 1 mL of fresh Alsever modified medium. Each hemocyte sample was thoroughly checked for quality, counted under a microscope using KOVA (Kova International, USA) slides, and stored on ice prior to processing. For scRNA-seq analysis, resuspended hemocytes were filtered on 30 µm filters, counted, the solution was adjusted to 1.10^6^ cells per mL and stored on ice prior to 10X genomic library preparation.

### *C. gigas* genome annotation

The *C. gigas* genome (Genbank GCA_902806645.1) (*30*) was used as a reference. Prior to annotation, the longest CDS sequences were extracted from the gff3 file. Annotation was realized using the ORSON script (https://gitlab.ifremer.fr/bioinfo/workflows/orson). ORSON combines cutting-edge tools for annotation processes within a Nextflow pipeline. The ORSON script performs a sequence similarity search with BLAST (*66*) against the Uniprot-Swissprot and Uniprot-trEMBL databases, and functional prediction with InterProScan (*67*) and eggNOG (*68*) orthogroup annotation. Interproscan analysis was performed against Pfam, Prosite, CDD, TIGR, SMART, SuperFamily, PRINTS and Hamap databases. Results were collected and processed using Blast2GO (*69*) for annotation mapping and validation.

### Drop Seq-based scRNA-seq library generation

The 10X Genomics protocol, Single Cell 3’ Reagent Kits v2 User Guide from the manufacturer (10X Genomics, USA) was followed to prepare gel in emulsion beads (GEM) containing single cells, hydrogel beads, and reverse transcription reagents, perform barcoded cDNA synthesis, and generate sequencing libraries from pooled cDNAs. The concentration of single-cell suspensions was approximately 1000 cells / μL, as estimated by manual counting, and cells were loaded according to the 10X protocol to capture approximately 3000 cells per reaction. Library construction (after GEM digestion) was performed using 10X reagents according to the manufacturer’s instructions. Libraries (paired-end reads 75 bp) were sequenced on an Illumina NovaSeq (Illumina, USA) using two sequencing lanes per sample.

### scRNA-seq analysis

Reads were aligned to the *C. gigas* reference genome (Genbank GCA_902806645.1) (*27*) using STAR solo software (v 2.7.10) (*29*). Unique molecular identifiers (UMIs) were extracted and counted for each cell, and an expression matrix was generated for further analysis. Single-cell RNA sequencing (scRNA-seq) data analysis was performed using the R programming language (version 4.2.1) (R Core Team, 2018) and the Seurat package (version 4.3.0) (*70*). The data were then pre-processed to remove unwanted sources of variation and to normalize gene expression. Cells with small library sizes and a high proportion of mitochondrial genes were excluded. Data normalization was performed using the SCTransform method. After normalization, highly variable genes were identified using the FindVariableFeatures function and the top variable genes were selected for downstream analyses. Dimensionality reduction was performed using Principal Component Analysis (PCA), followed by Uniform Manifold Approximation and Projection (UMAP) to visualize the data. Cell clustering was performed using the ‘FindClusters’ function, using the previously identified significant principal components (dims = 6) and a resolution parameter (r = 0.1) to define cluster granularity. Differential expression analysis was performed using the ‘FindAllMarkers’ function (pct.min = 0.25) to identify genes differentially expressed between clusters, with statistical significance determined using the Wilcoxon rank sum test. Functional enrichment analysis of differentially expressed genes was performed using gene set enrichment analysis (RBGOA) (*33*).

### KEGG pathway analysis

KEGG analysis was performed using DAVID Bioinformatics Resources (NIAID/NIH) (*32*). Gene lists of specifically overexpressed genes in each cluster were obtained after scRNA-seq processing (genes with Log2FC > 0.25 and significant p-value < 0.001) and used for KEGG annotation. The *C. gigas* reference genome from the DAVID bioinformatics resource was used for this analysis, with thresholds of 2 for counts and 0.05 for EASE value (p-value). KEGG annotation results were post-processed and presented as a heatmap showing the KEGG pathway, fold enrichment, p-value significance and number of positive terms.

### Rank-Based Gene Ontology Analysis

RBGOA first clusters GOs according to their representative genes to identify the most significant GOs, and then ranks the identified biological processes according to their average expression levels (over all representative genes). Finally, biological processes, molecular functions and cellular components significantly enriched in DEGs are identified by a Mann-Whitney rank test with a strict FDR correction. The files generated by the GO-MWU scripts (https://github.com/z0on/GO_MWU) were post-processed to extract the category names and the fraction indicating the number of “good candidates” relative to the total number of genes belonging to that category. The “good candidates” were the genes that exceeded an arbitrary ‘absValue’ cutoff (defined as 0.001) in their significance measure. The results were presented as a heatmap.

### Percoll Density Gradient Separation of Hemocytes10.3390/cells13242106

A concentration series of Percoll (Cytiva, Sweden) diluted in Alsever modified medium was prepared as follows: 10, 20, 30, 40, 50, 60 and 70 % (vol/vol). Discontinuous density gradients (from 10 % Percoll with a density of 1.0647 to 70 % Percoll with d=1.1049) were made using 1.5 mL of each concentration and loaded with 1 mL of the hemocyte suspension corresponding to approximately 2.10^7^ cells. Centrifugation was performed (30 min, 800 g, 4 °C) in a swinging bucket on a Beckman Coulter JS-13.1 rotor (Beckman Coulter, USA). Hemocytes concentrated at each density interface were collected separately with a 70 mm long 20Gx2.75” needle mounted on a 1 mL syringe. The hemocytes were then washed from the Percoll by adding 10 mL of ice-cold Alsever modified medium, pelleted by centrifugation (10 min, 200 g, 4°C) and resuspended in Alsever modified medium or filtered seawater.

### Cytological description of the hemocyte populations

200,000 fresh hemocytes were seeded onto a slide using a Cytospin 4 centrifuge (Thermo Scientific, USA). The samples were then stained using the panoptic MCDH (Micro Chromatic Detection for Hematology) (Cellavision, Sweden) staining protocol. This protocol produces purple hues typical of Romanowsky-Giemsa staining results. After staining, the samples were observed using a LEICA DMR (Leica AG, Germany) transmitted light microscope with a 40x magnification objective. Each slide was imaged and the hemocytes were counted and characterized based on their morphology.

### Real Time-quantitative Polymerase Chain Reaction (RT-qPCR)

Total RNA was extracted using the RNeasy kit (Qiagen, the Netherlands) and cDNA was synthesized from 1 μg total RNA using the Superscript IV kit (ThermoFisher Scientific, USA) with oligo(dT) primers. RT-PCR was performed on LightCycler© 480 thermocycler using the SYBR Green 1 Master kit (Roche, Switzerland). Primers were used at 200 nM. Primer sequences are listed in **Supplementary Table S3**. Expression was normalized to *Cg-rps6* reference gene. The standard cycling conditions were 95°C pre-incubation for 3 minutes followed by 40 cycles of 95°C for 10 seconds, 60°C for 20 seconds, and 72°C for 25 seconds. Each pair of primers was first tested and validated on a total hemocyte RNA preparation to control melting curves and establish calibration lines.

### Oxidative Burst Assay

The production of reactive oxygen species was quantified by luminescence assay. Briefly, hemocytes from hemolymph puncture or Percoll density gradient were washed once with filtered sterile water. 50 µL of hemocytes were plated in triplicate on a 96-well plate (3.10^5^ cells/cm²). 50 µL of 40 mM luminol (Merck and Co, USA) was added to each well. After 45 minutes of incubation at room temperature, the oxidative burst was induced by adding 100 µL of zymosan (Merck and Co, USA) at a multiplicity of infection (MOI) of 50:1. The plate was immediately placed in a Berthold Centro XS3 LB 960 luminescence microplate reader (Berthold GmbH, Germany) to measure luminescence emission every 2 minutes for 2 hours.

### Phagocytosis assay

400 µL of filtered, sterile water-washed hemocytes were seeded in a 24-well plate at a concentration of 3.10^5^ cells/cm². After 15 minutes, 50 µL of zymosan was added to the fractions at an MOI of 20:1. For LMG20012^T^ Vibrio, 50 µL of bacteria was added to a total hemolymph at an MOI of 5:1. After 1 hour of contact at room temperature, the cells were resuspended and 200 µL of the suspension was applied to a microscope slide using a Cytospin centrifuge. The slides were then stained with MCDH and observed under a LEICA DMR transmitted light microscope with a 40x magnification objective (Leica AG, Germany) to count phagocytic cells. A total of 2,807 cells were examined in three independent experiments.

### NitroBlueTetrazolium (NBT) staining

ROS production was measured using NitroBlue Tetrazolium reduction after zymosan stimulation. Briefly, 1 mL (1.10^6^ cells) of hemocyte solution was mixed with 50 µL of filtered NBT solution (15 mg/mL in water). Zymosan was added at a 4:1 MOI and the mixture was incubated at room temperature for 10 minutes on a rocking shaker. Then, 50 µL of each sample was plated onto glass coverslips and observed under a transmitted light microscope LEICA DMR with a 40x magnification objective (Leica AG, Germany) to count NBT-positive cells. The positions of positive NBT cells were recorded prior to MCDH staining to identify hemocyte types.

### Rhodanine copper staining of hemocytes

The storage of copper by hemocytes was examined by Rhodanine staining. Briefly, 1.10^5^ hemocytes were plated on Superfrost slides using a cytospin and circled with a hydrophobic pen to retain the staining solution. The Copper Stain Kit (SkyTek, USA) was used to stain the hemocytes. As described by the kit manufacturer, one drop of Rhodanine solution was added to 9 drops of acetate to form the working solution. Five drops were placed on the cytospin cells and a 5 mL Eppendorf tube with the cap removed was placed over the cells to prevent evaporation of the working solution. The slide and the balanced Eppendorf tube were placed in a beaker of boiling distilled water for 20 minutes. The slide was then washed with 5 drops of acetate and 3 drops of hematoxylin were placed on the slide for 1 minute at room temperature. The slides were then washed a final time with acetate and observed under a LEICA DMR transmitted light microscope with a 40x magnification objective (Leica AG, Germany) to count rhodanine-positive cells. The positions of rhodanine-positive cells were recorded prior to MCDH staining to identify hemocyte types.

### Pseudotemporal ordering of cells with Monocle3

Cells from the *C. gigas* dataset were analyzed using Monocle3 (https://github.com/cole-trapnell-lab/monocle3) (*42*). The Monocle3 analysis was performed on the Seurat object following the aforementioned processing steps. Clustering information (features, genes, partitions, clusters and UMAP coordinates) was transferred to a CDS object. The cell trajectory was calculated using the *learn_graph* function. The *choose_graph_segments* function was used to select three lineages. The gene expression along pseudotime data was extracted from the result. Then, the data were used to plot genes along pseudotime in three lineages using ggplot2 v3.4.4 R package and the heatmap was generated using the pheatmap v1.0.12 R package.

### Statistical Analysis

To evaluate differences between samples, a statistical analysis was performed using (version 4.2.1) (R Core Team, 2018) and appropriate packages. All data were examined for normality, and statistical tests were selected accordingly. One-way analysis of variance (ANOVA) was used for normally distributed data. Seven different hemolymph samples were used for cytological analysis. Oysters were provided by our local supplier and approximately 300 hemocytes were counted per sample. Seven independent experiments were performed for Percoll density gradient separation. A Tukey test was used to evaluate the statistical difference between the proportions of hemocyte types. The phagocytic capacity of hemocytes was tested in three independent experiments and statistical differences were evaluated using the Tukey test for both the phagocytic capacity between hemocyte types and the number of particles per phagocyte. Finally, oxidative burst capacity was tested 3 times on Percoll-separated hemocytes. The Tukey test was also used to assess statistical differences between conditions. The null hypothesis was rejected at a significance level of p = 0.05.

## Acknowledgments

We are grateful to the staff of the Ifremer platform of “La Tremblade” for technical support in animal housing. scRNA-seq data generated and used in this work were produced through the MGX platform (University of Montpellier, CNRS,INSERM). We thank the bioinformatic service of Ifremer (SEBIMER) for their help in bioinformatics and the qPCR platform GenSeq (University of Montpellier). We would like to thank Viviane Boulo and Danielle Mello for their enriching discussions, as well as all the members of the 2MAP laboratory team for the fruitful discussions throughout this project.

## Ethics

This work did not require ethical approval from a human subject or animal welfare committee.

## Author contributions

Conceptualization: G.M., D.D.-G., B.G., G.M.C. and E.V.

Methodology: S.d.L.F.D., J.P., G.M., D.D.-G., B.G., G.M.C. and E.V.

Investigation: S.d.L.F.D., J.P., G.M.C. and E.V. Supervision: G.M.C. and E.V.

Writing (original draft): S.d.L.F.D. and E.V.

Writing (review and editing): all authors.

## Declaration of AI use

We have not used AI-assisted technologies in creating this article.

## Data and materials availability

All data needed to evaluate the conclusions in the paper are present in the paper and/or the Supplementary Materials. Raw reads are available at ENA under the project accession number PRJEB74031.

