## Supplemental figures and tables for "Diversity and functional specialization of oyster immune cells uncovered by integrative single cell level investigations"

Sébastien de la Forest Divonne *et al.*

**This PDF file includes:**

Data S1 to S3
Tables S1 to S5
Figs. S1 to S12

**Data S1. Single-cell RNA-seq analysis result file.** CSV file containing the scRNA-seq analysis results with cluster number (<cluster>), gene number (<gene>), chromosome reference where the gene is located (<chromosome>), average expression level (<avg_log2FC>), percentage in the cluster (<pct1>) and in other clusters (<pct2>), pct1 / pct2 ratio (<pct. ratio>), adjusted p-val and p-val values (<p-val> and <p_val_adj>), and gene description (<description>).

**Data S2. Annotation file of CDS extracted from the *Crassostrea gigas* genome file.** CSV file containing the annotation results collected and processed using Blast2GO for annotation mapping and validation.

**Data S3. Compilation file for RBGOA results.** CSV file containing concatenated results of all RBGOA tests used to draw gene ontology analysis heatmaps with cluster number (<cluster_number>), GO term universe (<goterm_universe>), number of good gene candidates (<number_of_good_candidates>), total number of genes in this GOTerm category (<total_number_of_genes_of_this_category>), GO term name (<goterm_name>), adjusted p-value (<pval-adj>) and GO term name variation (<variation>).

| | |
|---|---|
| Number of Reads | 127,959,215 |
| Reads With Valid Barcodes | 97.03 % |
| Sequencing Saturation | 75.63 % |
| Q30 Bases in CB+UMI | 96.14 % |
| Q30 Bases in RNA read | 93.62 % |
| Reads Mapped to Genome: Unique+Multiple | 89.22 % |
| Reads Mapped to Genome: Unique | 82.01 % |
| Reads Mapped to Gene: Unique+Multiple Gene | 72.26 % |
| Reads Mapped to Gene: Unique Gene | 72.26 % |
| Estimated Number of Cells | 2937 |
| Unique Reads in Cells Mapped to Gene | 67,950,796 |
| Fraction of Unique Reads in Cells | 73.48 % |
| Mean Reads per Cell | 23,136 |
| Median Reads per Cell | 18,145 |
| UMIs in Cells | 16,306,374 |
| Mean UMI per Cell | 5552 |
| Median UMI per Cell | 4412 |
| Mean Gene per Cell | 1578 |
| Median Gene per Cell | 1434 |
| Total Gene Detected | 23,841 |

**Table S1. STARsolo summary metrics report.** Metrics dashboard obtained after the STARsolo step, describing the quality of the sequencing and the various characteristics of the cells detected after aligning the reads to the *C. gigas* genome from the Roslin Institute.

| Cluster | KEGG Prefix | KEGG Number | KEGG Pathway | Fold Enrichment | P-value (<0.05) | Genes |
|---|---|---|---|---|---|---|
| 1 | crg | 04144 | Endocytosis | 3.75 | 1.77E-06 | G32465, G28245, G29411, G28554, G23342,G27546, G4677, G4248, G23299, G4679<br>G25017, G27834, G23833, G20650, G1765,G2151, G15227, G11863, G17437 |
| 1 | crg | 00030 | Pentose phosphate pathway | 10.19 | 4.25E-05 | G30122, G34645, G20982, G31268, G25073<br>G11958, G17273 |
| 1 | crg | 00190 | Oxidative phosphorylation | 4.45 | 3.29E-04 | G20983, G16716, COX2, COX1, ND1<br>CYTB, G2226, G3525, G3658, ND5 |
| 1 | crg | 01200 | Carbon metabolism | 3.54 | 1.77E-03 | G30122, G18442, G34645, G20982, G31268<br>G16805, G25073, G11958, G17273, G7839 |
| 1 | crg | 00010 | Glycolysis / Gluconeogenesis | 4.94 | 6.58E-03 | G34645, G20982, G11958, G23780, G7839<br>G13028 |
| 1 | crg | 00620 | Pyruvate metabolism | 5.26 | 1.38E-02 | G18442, G22426, G22427, G23780<br>G7839 |
| 1 | crg | 03266 | Virion - Herpesvirus | 8.74 | 4.39E-02 | G22034, G32778<br>G32842 |
| 2 | crg | 03010 | Ribosome | 11.87 | 5.45E-27 | G5571, G25192, G5077, G3494, G33210,G26985, G26901, G23358, G1703, G1900<br>G1543, G2172, G12279, G5185, G26391,G10525, G4452, G11817, G25420, G2853<br>G4635, G3304, G25620, G5649, G3526,G740, G24712, G26976, G25626, G15236<br>G34618, G34615, G18702 |
| 2 | crg | 04814 | Motor proteins | 3.64 | 3.62E-04 | G32475, G24506, G4572, G3088, G1531<br>G15286, G1299, G153, G22991, G1471<br>G10110, G1470 |
| 2 | crg | 00980 | Metabolism of xenobiotics by cytochrome P450 | 4.61 | 2.17E-02 | G5106, G26659, G18623, G18624<br>G14458 |
| 3 | crg | 00190 | Oxidative phosphorylation | 8.03 | 1.47E-12 | G31970, G3073, G5034, G3298, G5653,G27276, G25266, G650, G331, G23645<br>G1779, G16296, G23701, G13193, G27816,G10181, G15131, G21371, G2133, G34736 |
| 3 | crg | 01200 | Carbon metabolism | 5.42 | 4.82E-08 | G26194, G5034, G4037, G3500, G20764,G16296, G23701, G14096, G22912, G15171<br>G27816, G21794, G19054, G15078, G16455,G34649, G9692 |
| 3 | crg | 00010 | Glycolysis / Gluconeogenesis | 7.41 | 5.01E-06 | G26194, G14096, G22912, G21794, G19054<br>G15078, G4037, G3500, G34649, G12763 |
| 3 | crg | 01230 | Biosynthesis of amino acids | 5.77 | 1.41E-05 | G26194, G14096, G22912, G21794, G4037,G25300, G34649, G7042, G25304, G9692<br>G20764 |
| 3 | crg | 04144 | Endocytosis | 3.2 | 3.22E-05 | G10612, G34830, G4232, G31680, G28015,G28554, G2650, G35288, G23530, G22892<br>G3216, G2800, G22524, G25627, G21676,G20683, G1765, G11863 |
| 3 | crg | 00020 | Citrate cycle (TCA cycle) | 8.23 | 1.48E-04 | G16296, G23701, G5034, G15171, G27816<br>G19054, G16455 |
| 3 | crg | 01100 | Metabolic pathways | 1.27 | 1.48E-02 | G13605, G5034, G3012, G3298, G650,G331, G22961, G5758, G14096, G27816<br>G13000, G2033, G4037, G3500, G34394,G23387, G25763, G20764, G1779, G23701<br>G13193, G21533, G15171, G19054, G15131,G1375, G21371, G12861, G9692, G3073<br>G8883, G31970, G30520, G25177, G27276,G5653, G16296, G2133, G16455, G34649<br>G7042, G9342, G26194, G23482, G6556,G23164, G5106, G25300, G25266, G25304<br>G23645, G25904, G22912, G21794, G15078,G10181, G22919, G16488, G34736, G12763<br>G2280 |
| 3 | crg | 00250 | Alanine, aspartate and glutamate metabolism | 4.26 | 2.79E-02 | G23164, G25300, G25763, G25304<br>G20764 |
| 3 | crg | 00430 | Taurine and hypotaurine metabolism | 5.05 | 4.23E-02 | G23482, G34394, G25763<br>G9342 |
| 4 | crg | 03010 | Ribosome | 11.18 | 2.38E-62 | G32202, G25192, G4582, G5077, G3494,G33210, G33255, G5115, G5555, G11948<br>G31071, G23694, G4502, G732, G1703,G1900, G21588, G18212, G1543, G28190<br>G12279, G13687, G28274, G10525, G7023,G27586, G3387, G11817, G26894, G4635<br>G3304, G2853, G24036, G5649, G24712,G26976, G1172, G12546, G27592, G11965<br>G5571, G4085, G5375, G5652, G23471,G32582, G26985, G26901, G23358, G18432<br>G16773, G21360, G10473, G2172, G35349,G5185, G26391, G4452, G4972, G3885<br>G25420, G25620, G3526, G740, G2636,G25626, G16242, G15236, G34618, G1350<br>G34615, G18702 |
| 4 | crg | 03050 | Proteasome | 9.09 | 3.16E-14 | G11447, G11879, G9178, G28086, G6441,G27078, G6503, G27105, G4954, G21626<br>G1667, G22517, G11085, G13469, G32646,G1140, G29381, G12565, G30766, G34613 |
| 4 | crg | 03040 | Spliceosome | 4.6 | 1.62E-13 | G30069, G23072, G3035, G27237, G4644,G67, G3357, G28826, G21225, G22448<br>G21463, G20152, G31757, G12256, G16599,G2099, G16114, G11265, G19821, G5284<br>G4177, G23063, G3388, G35242, G23100,G2896, G5803, G21555, G22119, G18742<br>G13575, G14401, G10466 |
| 4 | crg | 00190 | Oxidative phosphorylation | 3.14 | 3.88E-04 | G31970, G3077, G3298, G26024, G34551,G11926, G27610, G650, G2799, G17080<br>G1779, G1500, ND3, G31998 |
| 4 | crg | 03015 | mRNA surveillance pathway | 3.18 | 1.10E-03 | G21555, G1768, G21245, G23072, G3035,G1892, G27237, G22452, G12561, G11265<br>G19821, G15119 |
| 4 | crg | 03013 | Nucleocytoplasmic transport | 2.44 | 9.12E-03 | G21555, G1768, G23072, G4818, G3035,G2485, G31807, G27237, G12561, G11265<br>G19821, G27008 |
| 4 | crg | 04141 | Protein processing in endoplasmic reticulum | 2.06 | 1.28E-02 | G11932, G29430, G10911, G22082, G26491,G29962, G2451, G2694, G6732, G4985<br>G20713, G204, G20332, G2609, G10994 |
| 5 | crg | 03010 | Ribosome | 7.41 | 1.68E-12 | G27592, G32202, G5185, G5375, G33210,G27586, G11948, G26985, G3885, G26894<br>G3304, G2853, G18212, G18432, G16773,G1543, G16242, G21360, G13687, G2172<br>G12279 |
| 5 | crg | 00190 | Oxidative phosphorylation | 4.59 | 6.31E-04 | G3430, COX2, COX1, ND1, CYTB<br>ND3, ND2, ND5, ND4 |
| 5 | crg | 04137 | Mitophagy - animal | 4.65 | 8.54E-03 | G30487, G25784, G29415, G3710, G22696<br>G12500 |
| 6 | crg | 03010 | Ribosome | 16.48 | 2.26E-55 | G32202, G25192, G4582, G3494, G5115,G5555, G11948, G4502, G732, G1703<br>G21588, G18212, G1543, G28190, G12279,G13687, G10525, G7023, G27586, G3387<br>G11817, G26894, G4635, G3304, G2853,G24036, G5649, G24712, G26976, G27592<br>G5571, G4085, G5652, G26985, G26901,G23358, G18432, G16773, G21360, G10473<br>G2172, G5185, G26391, G3885, G25420,G25620, G740, G2636, G25626, G16242<br>G15236, G34618, G34615, G18702 |
| 7 | crg | 04144 | Endocytosis | 4.04 | 4.91E-09 | G10612, G23299, G27538, G22524, G12273,G16951, G2151, G3185, G6211, G13317<br>G2450, G31680, G4232, G28015, G2650,G35288, G23342, G20567, G17254, G18762<br>G19412, G21991, G1774, G1993, G30759 |
| 7 | crg | 04137 | Mitophagy - animal | 4.99 | 3.50E-04 | G18587, G25784, G11088, G2334, G30759<br>G3710, G27737, G30997, G12500 |
| 7 | crg | 04145 | Phagosome | 2.82 | 1.88E-03 | G3185, G22181, G3430, G28015, G24310,G17795, G14056, G2026, G1780, G30759<br>G32778, G32842, G90 |
| 7 | crg | 04140 | Autophagy - animal | 2.97 | 3.47E-03 | G20344, G5397, G25608, G25784, G11385,G2024, G11088, G30759, G10907, G21965<br>G12500 |
| 7 | crg | 04310 | Wnt signaling pathway | 2.9 | 6.82E-03 | G24405, G23180, G11208, G35235, G16231<br>G2024, G14136, G2334, G1780, G2797 |
| 7 | crg | 04142 | Lysosome | 1.92 | 4.51E-02 | G11645, G5397, G22181, G3430, G2026,G27524, G18705, G26324, G16202, G2976<br>G27526, G23337 |

**Table S2. Table presenting the result of KEGG analysis performed using DAVID Bioinformatics Resources.** GO term enrichment analysis was conducted on specifically overexpressed genes in each cluster obtained after scRNA-seq processing (genes with Log2FC > 0.25 and significant p-value < 0.001) to highlight the most relevant GO terms associated with a given gene list. The visualization of the different pathways can be obtained from the KEGG website using the KEGG prefix and KEGG number (https://www.genome.jp/kegg/pathway.html)

| Target | Primer name | Sequence |
|---|---|---|
| Laccase 24 | LACC24-F | CCT-TGA-TTC-TTC-TTG-CCA-TCC-G |
| | LACC24-R | AAA-GCT-TGC-GAT-CTT-TGG-CAA |
| C-type lectin domain-containing protein | CLEC-F | ATC-GGC-TTC-TAC-ATG-GAC-TGA-C |
| | CLEC-R | GTG-TCT-AAA-GCT-GCG-CCG-AT |
| Putative modulator of levamisole receptor-1 | LEVAR-F | GTG-ACA-GAC-TTC-CCT-CAC-CCT |
| | LEVAR-R | GCA-CTG-AGT-CGA-GTC-GTA-TGT |
| EGF-like domain-containing protein 8 | EGFL-F | GAG-TGT-TTG-ACA-GGA-CGA-AGC |
| | EGFL-R | CAT-CAT-CGT-TTC-CAA-CTG-AGG-C |
| X-Box binding protein-like domain | XBOX-F | GGG-TCA-ACA-GTG-CTA-GGC-AAT |
| | XBOX-R | GTA-AGC-CAC-CAT-CCC-TAC-CAC |
| Transglutaminase-like domain containing protein | TGC-F | CTA-CAA-GCT-GGA-CAC-CAC-CAA |
| | TGC-R | GCA-TTG-ACC-AGT-GAC-ACA-GTC |
| MD-2-related lipid-recognition domain containing protein | MLDP-F | CTT-GGA-CCT-CGT-TAT-CTT-CGC |
| | MLDP-R | CTC-CCT-CTG-GTC-CAC-AAA-CAA |
| High mobility group protein B1 | HMGB1 | GCC-CAC-GCT-GAA-CTA-TAC-AAG |
| | HMGB1 | CAC-CTT-GTA-GTC-CCT-GAG-TGG |
| Galectin | GAL-F | CCA-CAG-TAT-CAA-CGA-CCC-TCC |
| | GAL-R | TCA-CTA-CCG-TCA-TAG-GGA-CCG |
| Cubilin | CUBN-F | TAA-GTT-CAC-TCT-GGC-CCA-AGG |
| | CUBN-R | GCT-CAT-GAT-CGT-AGT-GGT-GCT |
| Natterin-1 | NAT1-F | CCG-TAC-GAT-GGT-GAG-GAG-AAA |
| | NAT1-R | CCG-TCC-CAC-ATA-CTT-GTC-GTT |
| Caveolin | CAV-F | TCC-AAA-TGA-CCA-TGA-CCC-AGA |
| | CAV-R | ACT-CTA-TTC-TTG-GTC-GCC-TGG |
| G Protein receptor F1-2 domain-containing protein | GPROT-F | CGA-ACG-CCT-GCT-TCT-GAT-ATG |
| | GPROT-R | TCC-ACA-TCG-AAT-GCT-CTG-TCT |
| DBH-like monooxygenase protein 1 | MOX-F | CCT-CCG-CAG-CAA-GAA-GAA-GTA |
| | MOX-R | TTC-TGT-TTC-GTC-CTC-TCC-ACG |
| Ribosomal protein S6 | RPS6-F | CAG-AAG-TGC-CAG-CTG-ACA-GTC |
| | RPS6-R | AGA-AGC-AAT-CTC-ACA-CGG-AC |
| Big Defensin 1&2 | BigDef1-2-F | TTC-GCC-TGC-TTC-CAT-ACT-GG |
| | BigDef1-2-R | GTC-ATG-GTC-ACT-CCT-TAT-TC |
| Hemocyte defensin | HemDef-F | CTA-CCA-GTT-GTT-CAT-ACA-GAG |
| | HemDef-R | TCT-TGG-TCA-GAT-TCA-GTC-TGG |
| Bactericidal Permeability Increasing Protein | BPI | GGA-GGC-GGA-AAT-GGA-TTA-CT |
| | BPI | TGG-TTG-ACA-TCG-TTG-CTG-AC |

**Table S3. Sequences of primers used in this study.** Name of the transcript targeted by the primer pair, name of the primer used in this study and nucleotide sequence of each primer.

| Fraction | Cells | Average | Std.Dev. | Std.Error | Min | Max |
|---|---|---|---|---|---|---|
| 1 | H | 46.43 | 9.47 | 4.73 | 37.50 | 59.63 |
| 1 | ML | 20.26 | 9.09 | 4.55 | 7.34 | 28.38 |
| 1 | BBL | 14.47 | 2.29 | 1.15 | 12.16 | 16.50 |
| 1 | ABL | 15.50 | 4.94 | 2.47 | 8.11 | 18.35 |
| 1 | SGC | 0.35 | 0.44 | 0.22 | 0.00 | 0.92 |
| 1 | BGC | 0.38 | 0.75 | 0.38 | 0.00 | 1.50 |
| 1 | VC | 2.62 | 1.97 | 0.98 | 0.92 | 5.41 |

| Fraction | Cells | Average | Std.Dev. | Std.Error | Min | Max |
|---|---|---|---|---|---|---|
| 5 | H | 4.29 | 4.41 | 2.21 | 0.76 | 10.45 |
| 5 | ML | 19.67 | 12.04 | 6.02 | 3.03 | 30.88 |
| 5 | BBL | 3.56 | 1.91 | 0.95 | 1.49 | 6.06 |
| 5 | ABL | 3.35 | 2.12 | 1.06 | 0.75 | 5.88 |
| 5 | SGC | 40.92 | 7.31 | 3.66 | 33.82 | 49.24 |
| 5 | BGC | 4.50 | 2.98 | 1.49 | 1.47 | 8.33 |
| 5 | VC | 23.71 | 4.48 | 2.24 | 17.91 | 28.79 |

| Fraction | Cells | Average | Std.Dev. | Std.Error | Min | Max |
|---|---|---|---|---|---|---|
| 2 | H | 20.17 | 5.26 | 2.63 | 14.10 | 26.56 |
| 2 | ML | 27.64 | 4.48 | 2.24 | 22.79 | 33.33 |
| 2 | BBL | 27.21 | 9.94 | 4.97 | 16.22 | 39.71 |
| 2 | ABL | 20.17 | 2.36 | 1.18 | 16.91 | 22.52 |
| 2 | SGC | 1.14 | 0.79 | 0.39 | 0.00 | 1.80 |
| 2 | BGC | 1.03 | 1.26 | 0.63 | 0.00 | 2.56 |
| 2 | VC | 2.64 | 1.56 | 0.78 | 0.74 | 4.50 |

| Fraction | Cells | Average | Std.Dev. | Std.Error | Min | Max |
|---|---|---|---|---|---|---|
| 6 | H | 3.57 | 3.82 | 1.91 | 0.65 | 9.09 |
| 6 | ML | 13.95 | 4.91 | 2.45 | 7.10 | 18.18 |
| 6 | BBL | 1.19 | 1.38 | 0.69 | 0.00 | 2.58 |
| 6 | ABL | 1.54 | 0.65 | 0.33 | 0.65 | 2.17 |
| 6 | SGC | 68.00 | 10.11 | 5.06 | 58.70 | 78.46 |
| 6 | BGC | 1.69 | 1.99 | 1.00 | 0.00 | 3.87 |
| 6 | VC | 10.06 | 5.30 | 2.65 | 3.08 | 15.94 |

| Fraction | Cells | Average | Std.Dev. | Std.Error | Min | Max |
|---|---|---|---|---|---|---|
| 3 | H | 12.49 | 3.78 | 1.89 | 10.08 | 18.04 |
| 3 | ML | 28.26 | 8.90 | 4.45 | 17.12 | 38.66 |
| 3 | BBL | 21.18 | 3.11 | 1.55 | 16.89 | 24.32 |
| 3 | ABL | 17.71 | 5.78 | 2.89 | 11.86 | 25.21 |
| 3 | SGC | 7.96 | 5.30 | 2.65 | 3.09 | 14.41 |
| 3 | BGC | 4.71 | 4.16 | 2.08 | 0.00 | 10.14 |
| 3 | VC | 7.68 | 2.87 | 1.43 | 4.20 | 10.81 |

| Fraction | Cells | Average | Std.Dev. | Std.Error | Min | Max |
|---|---|---|---|---|---|---|
| 7 | H | 2.43 | 3.04 | 1.52 | 0.00 | 6.31 |
| 7 | ML | 8.50 | 6.35 | 3.18 | 1.93 | 17.05 |
| 7 | BBL | 1.67 | 1.51 | 0.75 | 0.00 | 3.38 |
| 7 | ABL | 0.36 | 0.72 | 0.36 | 0.00 | 1.45 |
| 7 | SGC | 81.18 | 8.99 | 4.49 | 68.18 | 88.41 |
| 7 | BGC | 2.68 | 1.50 | 0.75 | 0.90 | 4.55 |
| 7 | VC | 3.18 | 2.92 | 1.46 | 0.00 | 6.82 |

| Fraction | Cells | Average | Std.Dev. | Std.Error | Min | Max |
|---|---|---|---|---|---|---|
| 4 | H | 7.82 | 5.97 | 2.99 | 2.20 | 14.29 |
| 4 | ML | 32.69 | 13.33 | 6.66 | 19.34 | 50.82 |
| 4 | BBL | 9.64 | 3.40 | 1.70 | 4.76 | 12.30 |
| 4 | ABL | 5.37 | 0.97 | 0.48 | 4.42 | 6.67 |
| 4 | SGC | 18.29 | 10.39 | 5.19 | 5.74 | 30.94 |
| 4 | BGC | 3.90 | 1.51 | 0.76 | 2.20 | 5.71 |
| 4 | VC | 22.29 | 8.25 | 4.12 | 11.48 | 30.77 |

**Table S4. Hemocyte composition of the 7 Percoll fractions used for qPCR analysis.** For each cell type in each fraction, the table presents the average percentage, standard deviation, standard error, minimum, maximum, and median count values.

| | | | Blast alignment results | | | | | | *Homo sapiens* Homologs | |
|---|---|---|---|---|---|---|---|---|---|---|
| Gene Number | Annotation Name | TF Name | Best Hit Blast Name | Specie | Accession number | Bit Score | Percentage Identity | Coverage | Name | Uniprot Number |
| G29966 | Krueppel-like factor 15 | CgKLF6 | Krueppel-like factor 6 | *C.gigas* | XP_011428991 | 659.448 | 100 | 100 | HsKLF8 | O95600 |
| G30997 | BHLH domain-containing protein | CgTFEB | Transcription factor EC isoform X1 | *C.gigas* | XP_011447558 | 1089.72 | 100 | 100 | HsTFEB | P19484 |
| G17147 | BZIP domain-containing protein | CgCR3L1 | cAMP-responsive element-binding protein 3-like protein 1 isoform 1 | *H.sapiens* | NP_443086 | 1060.06 | 100 | 100 | HsCR3L2 | Q70SY1 |
| G3043 | Transcriptional regulator | CgMYC | Transcriptional regulator Myc-A-like | *C.angulata* | XP_052688275 | 548.128 | 100 | 100 | HsMYC | P01106 |
| G31054 | GATA-binding factor 3 | CgGATA3 | Transcription factor GATA-3 isoform X1 | *C.gigas* | XP_011412780 | 966.066 | 100 | 100 | HsGATA2 | P23769 |
| G11196 | BZIP domain-containing protein | CgFOS | Proto-oncogene c-Fos | *C.gigas* | XP_011446784 | 664.84 | 100 | 100 | HsATF3 | P18847 |
| G10637 | BZIP domain-containing protein | CgFOS2 | Fos-related antigen 2 | *C.gigas* | XP_011439871 | 271.937 | 100 | 100 | HsATF4 | P18848 |
| G2123 | SPRY domain-containing SOCS box protein 3 | CgSOCS3 | SPRY domain-containing SOCS box protein 3 | *C.gigas* | XP_011413771 | 488.804 | 100 | 100 | HsSPSB4 | Q96A44 |
| G1067 | BZIP domain-containing protein | none | uncharacterized protein LOC105347143 | *C.gigas* | XP_019930445.2 | 1419.35 | 100 | 100 | none | none |
| G1708 | SoxE | CgSOX8 | Transcription factor Sox-8-like | *C.gigas* | NP_001295801 | 947.192 | 99.8 | 100 | HsSOX9 | P48436 |
| G3506 | BZIP domain-containing protein | CgATF5 | cAMP-dependent transcription factor ATF-4 | *C.gigas* | XP_011439092 | 566.614 | 85.9 | 100 | HsATF5 | Q9Y2D1 |
| G27827 | BHLH domain-containing protein | CgUSF2 | Upstream stimulatory factor 2-like isoform X4 | *C. angulata* | XP_052673994 | 605.905 | 100 | 99.68 | HsUSF2 | Q15853 |
| G11198 | BZIP domain-containing protein | CgFOS3 | Fos-related antigen 1 | *C.gigas* | XP_034323853 | 456.062 | 100 | 99.62 | none | none |
| G31522 | Helix-loop-helix protein 1 | none | uncharacterized protein LOC105333018 isoform X2 | *C. gigas* | XP_011434110 | 377.096 | 100 | 99.45 | none | none |
| G10636 | BZIP domain-containing protein | CgATF1 | Basic leucine zipper transcriptional factor ATF-like isoform X1 | *C.gigas* | XP_011439867 | 270.011 | 100 | 99.41 | none | none |
| G13555 | BZIP domain-containing protein | CgJUNB | Transcription factor jun-B isoform X1 | *C.gigas* | XP_011440881 | 224.557 | 100 | 99.33 | none | none |
| G21091 | X-box binding protein like protein | CgXBOX | X-box binding protein-like protein | *C.ariakensis* | AEF33390 | 375.491 | 80.5 | 99.12 | none | none |
| G12164 | Estrogen receptor | CgRXRB | Estrogen receptor beta | *C.gigas* | XP_011424805 | 1395.56 | 100 | 98.79 | HsRXRA | P19793 |
| G748 | Interferon regulatory factor | CgIRF1 | Interferon regulatory factor 1 isoform X1 | *C.gigas* | XP_011449592 | 802.742 | 100 | 95.51 | HsIRF1 | P10914 |
| G5204 | ETS-related transcription factor Elf-3 | CgELF3 | ETS homologous factor isoform X1 | *C.gigas* | XP_034310128 | 883.248 | 100 | 95.25 | HsELF3 | P78545 |
| G28398 | ETS domain-containing protein | CgELK3 | ETS domain-containing protein Elk-3 isoform X1 | *C.gigas* | XP_011421195 | 753.051 | 100 | 93.91 | HsELK3 | P41970 |
| G35512 | cAMP-responsive element modulator | CgCREM | cAMP-responsive element modulator isoform X2 | *C.gigas* | XP_011448206 | 572.778 | 100 | 92.79 | HsCREB1 | P16220 |
| G11013 | BZIP domain-containing protein | CgTEF | Thyrotroph embryonic factor-like | *C.virginica* | XP_022332233 | 570.466 | 86.4 | 85.1 | HsTEF | Q10587 |
| G7003 | ETS-related transcription factor Elf-4 | CgELF2 | ETS-related transcription factor Elf-2 | *C.gigas* | XP_011447491 | 523.857 | 99.6 | 84.54 | HsELF1 | P32519 |
| G27920 | LIM domain transcription factor LMO4.1 | CgLMO4 | LIM domain transcription factor LMO4.1 isoform X2 | *C.gigas* | XP_011443581 | 319.316 | 98.2 | 81.86 | HsLMO4 | P61968 |
| G10865 | BZIP domain-containing protein | CgVBP | Transcription factor VBP | *C.gigas* | XP_011457057 | 438.343 | 100 | 77.95 | HsHLF | Q16534 |
| G35439 | BZIP domain-containing protein | CgCEBPG | CCAAT/enhancer-binding protein beta | *C.gigas* | XP_011452433 | 442.195 | 100 | 75.61 | HsCEBPG | P53567 |
| G2334 | Proto-oncogene c-Jun | CgJUN | Transcription factor AP-1 | *C.gigas* | XP_034313954 | 429.098 | 100 | 58.31 | HsJUN | P05627 |
| G26127 | Krueppel-like factor 5 | CgKLF5 | Krueppel-like factor 5 isoform X1 | *C.gigas* | XP_011421316 | 905.975 | 100 | 57.28 | HsKLF7 | O75840 |
| G20682 | cAMP-responsive element-binding protein-like 2 | CgCRBL1 | cAMP responsive element binding protein-like, partial | *C.gigas* | AAU93879 | 318.546 | 99.4 | 56.83 | HsCRBL2 | O60519 |
| G17294 | Runt-related transcription factor 1 | CgRUNX1 | Runt-related transcription factor 1-like isoform X5 | *C.angulata* | XP_052711215 | 1085.86 | 100 | 56.68 | HsRUNX3 | Q13761 |
| G10077 | BZIP domain-containing protein | CgCR3L4 | Clumping factor A isoform X1 | *C.gigas* | XP_011425092 | 775.393 | 100 | 56.11 | HsCR3L3 | Q68CJ9 |
| G2021 | BHLH domain-containing protein | CgTAL1 | T-cell acute lymphocytic leukemia protein 1-like isoform X1 | *C.gigas* | XP_011439324 | 394.045 | 100 | 49.23 | HsTAL2 | Q16559 |
| G2933 | Putative transcription factor SOX-14 | CgSOX11 | Transcription factor Sox-11 | *C.gigas* | XP_011445203 | 424.091 | 100 | 43.97 | HsSOX12 | O15370 |
| G11358 | BZIP domain-containing protein | CgGIANT | Protein giant | *C.gigas* | XP_011417277 | 486.108 | 100 | 38.41 | HsTEF | Q10587 |
| G6354 | ETS domain-containing protein | CgPU.1 | ETS-related transcription factor Elf-3 | *C.gigas* | XP_034320185 | 189.119 | 100 | 36 | HsPU.1 | P17947 |

**Table S5. Transcription factors identified in the scRNA-seq dataset of *Crassostrea gigas* hemocytes.** Transcription factors identified in the scRNA-seq dataset and their homology with human proteins, as indicated by the blast alignment results. Each entry includes the gene identifier, the protein it represents in *C.gigas*, and its human counterpart. The bit score and coverage indicate the strength and extent of the alignment, respectively.

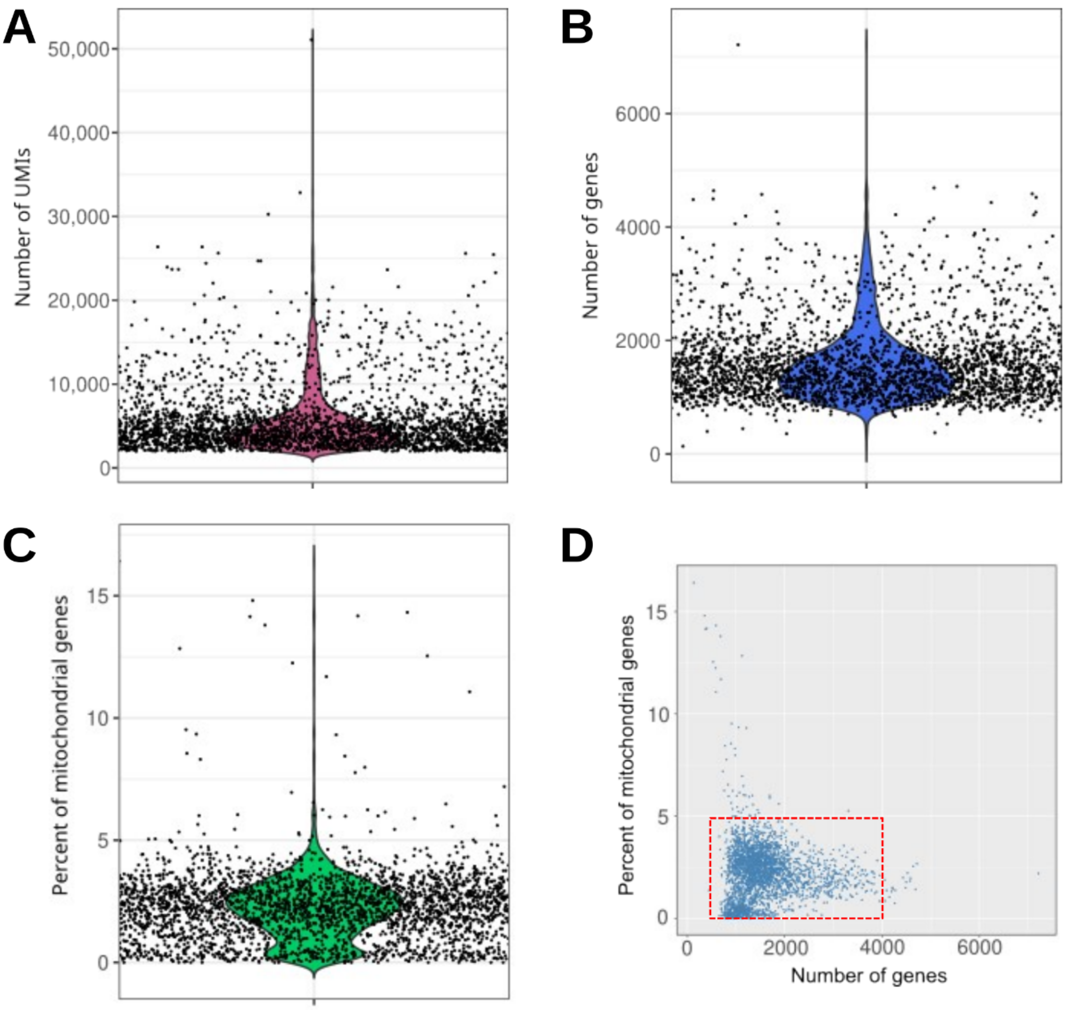


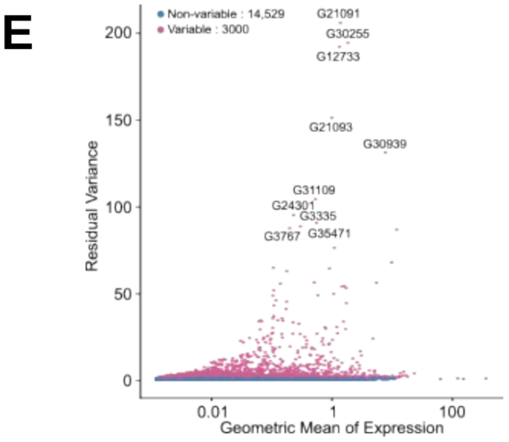


F

| | |
|---|---|
| Number of cells before filtering | 2937 |
| Average number of genes detected by cell | 1578 ± 643 |
| Average number of UMIs detected by cell | 5552 ± 3848 |
| Filtering parameters | |
| Cells with gene between 750 and 4000 | |
| Cells with less than 5 % of mitochondrial genes | |
| Number of cells after filtering | 2817 |
| Average number of genes detected by cell | 1568 ± 590 |
| Average number of UMIs detected by cell | 5432 ± 3462 |

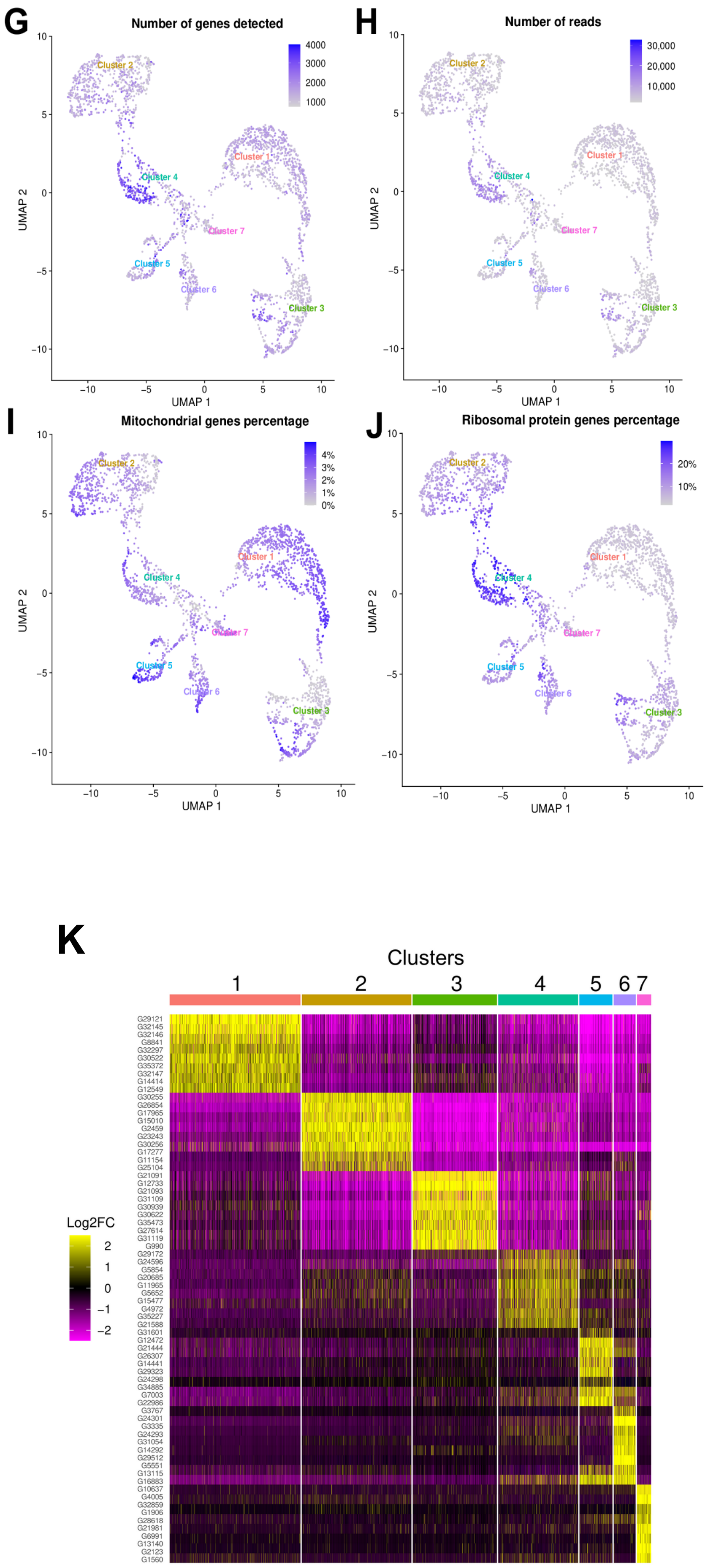
G
Number of genes detected
4000
3000
2000
1000
UMAP 2
UMAP 1
H
Number of reads
30,000
20,000
10,000
I
Mitochondrial genes percentage
4%
3%
2%
1%
0%
J
Ribosomal protein genes percentage
20%
10%
Cluster 1
Cluster 2
Cluster 3
Cluster 4
Cluster 5
Cluster 6
Cluster 7
K
Clusters
1
2
3
4
5
6
7
Log2FC
2
1
0
−1
−2

**Figure S1. ScRNA-seq quality control metrics.** Distribution of **A)** unique molecular identifiers (UMIs), **B)** genes and **C)** percentage of mitochondrial genes detected per cell. Each dot represents one cell. **D)** Plot of the percentage of mitochondrial genes versus the number of genes detected in each cell. The red box represents the cells selected for further analysis (number of genes detected between 750 and 4000 and with a percentage of mitochondrial genes less than 5%) **E)** The *FindVariableFeatures()* function was used to identify features with high cell-to-cell variation in the dataset to highlight the biological signal in the single cell dataset. **F)** Table summarizing some quality control metrics. The table shows the thresholds to remove poor quality cells (doublets or empty droplets). The number of cells, UMIs and genes before and after filtering are shown. **G)** Uniform Manifold Approximation and Projection (UMAP) plot of the cells with the number of expressed genes. **H)** UMAP plot of the number of reads per cell. **I)** UMAP plot of the percentage of mitochondrial genes in each cell. **J)** UMAP graph of the percentage of ribosomal protein transcripts in each cell. **K)** Heatmap showing the top 10 enriched marker genes in each cell per cluster as determined by FindAllMarkers() function in Seurat after secondary bioinformatic processing, corresponding to clusters in UMAP plots from **Fig. 1B**, ranked by log2FC.

**A**

| Stage | Number of CDS | % of CDS |
|---|---|---|
| Input | 30,724 | 100 |
| **Database querying** | | |
| Blast annoted | 30,511 | 99.3 |
| Interpro annoted | 30,718 | 99.9 |
| EggNogg annoted | 12,630 | 41.1 |
| **Blast2Go** | | |
| Mapped | 28,570 | 92.9 |
| Annotated | 22,462 | 73.1 |

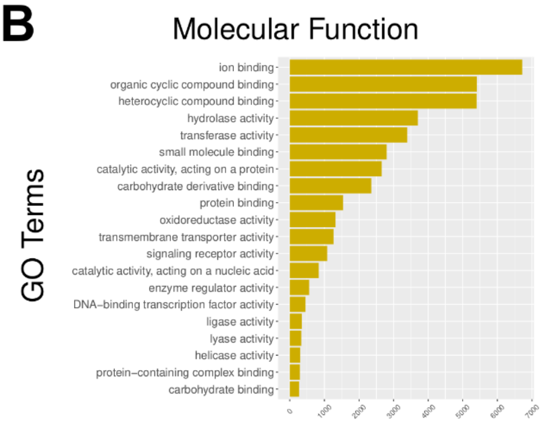


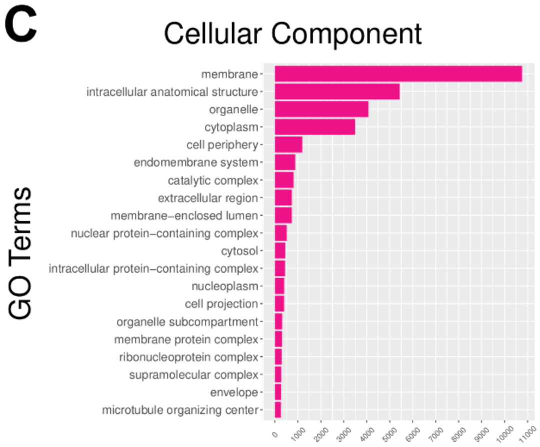


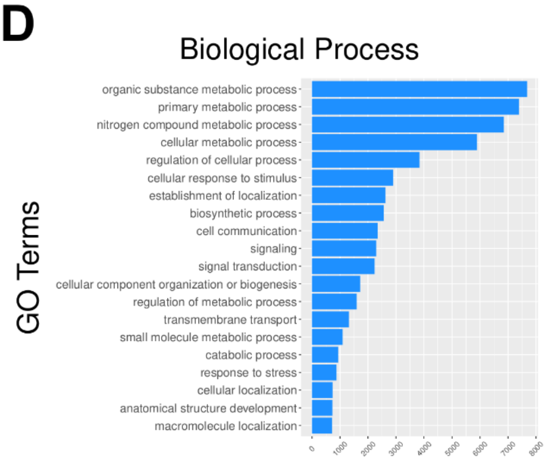


**Figure S2. Results of the *C. gigas* genome re-annotation. A)** The number and percentage of Coding DNA Sequences (CDS) with valid annotation after each annotation step are shown. A BLAST query was performed against the TrEMBL/Uniprot database and InterproScan annotation against Pfam, PrositeSiteProfiles, CDD, TIGRFAM, PRINTS, SMART, SUPERFAMILY and Hamap databases. Blast and InterProScan results were compiled and processed using Blast2Go. A first mapping step was used to enrich the Blast result with GO-terms, and the annotation step was used to optimize and validate the GO-terms annotations. **B) C)** and **D)** show the distribution of the various categories of GO-terms across the three primary domains of Gene Ontology : Molecular Function, Cellular Component and Biological Process, respectively.

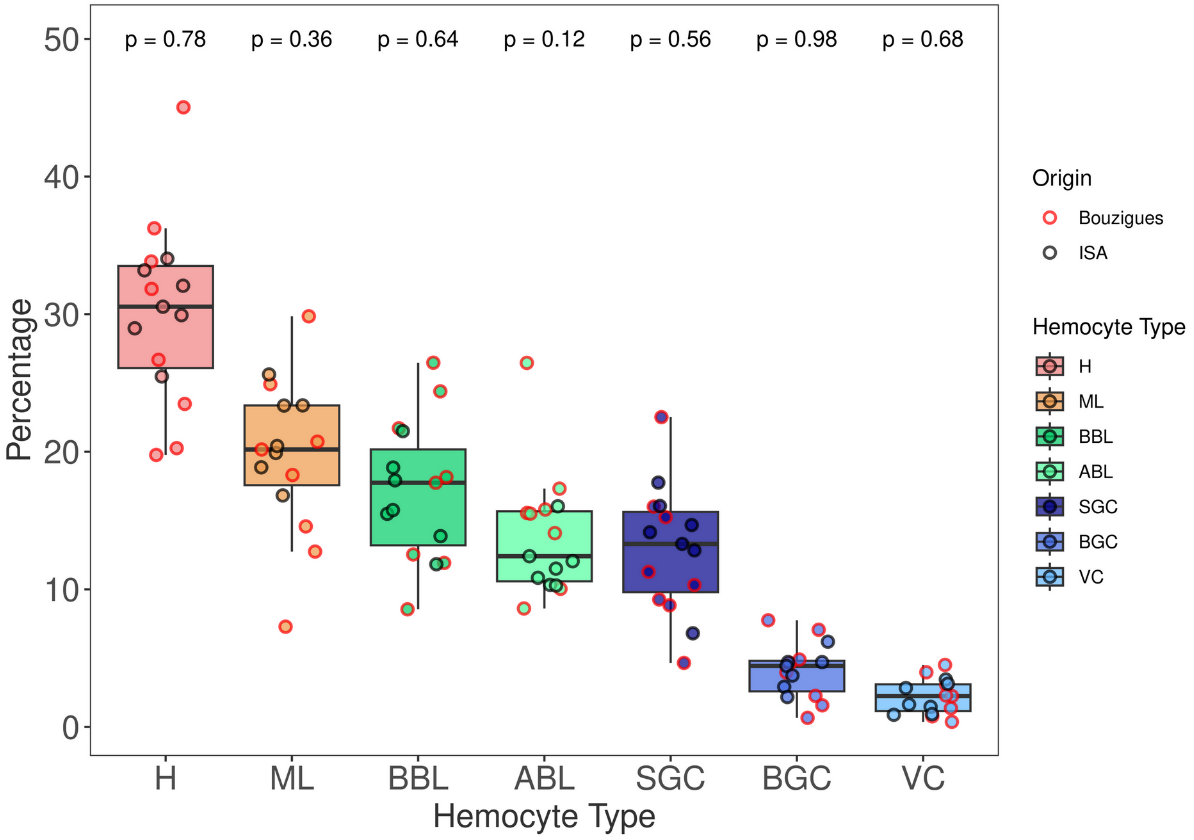


| | Hyalinocyte (H) | Basophilic Blast-Like (BBL) | Acidophilic Blast-Like (ABL) | Macrophage Like (ML) | Small Granule Cells (SGC) | Big Granules Cell (BGC) | Vesicular Cell (VC) |
|---|---|---|---|---|---|---|---|
| **Bouzigues** | 29.64 ± 8.74 % | 17.68 ± 6.34 % | 15.42 ± 5.38 % | 18.57 ± 7.09 % | 12.26 ± 5.50 % | 4.10 ± 2.53 % | 2.32 ± 1.47 % |
| **ISA** | 30.60 ± 2.89 % | 16.46 ± 3.24% | 11.92 ± 1.99 % | 21.19 ± 3.05 % | 13.65 ± 3.45 % | 4.13 ± 1.32 % | 2.05 ± 1.07 % |
| **Student test – pval** | ns - 0.78 | ns - 0.64 | ns - 0.12 | ns - 0.36 | ns - 0.56 | ns - 0.98 | ns - 0.68 |

**Figure S3. Distribution of hemocyte populations in the hemolymph of oysters from ISA and Thau lagoon origins.** Hemolymph was collected from oysters raised in ISA (Ifremer Standardized Animals, La Tremblade, GPS : 45.7981624714465, -1.150171788447683) and Thau lagoon (Bouzigues, GPS : 43.44265228308842, 3.6359883059292057). Hemocytes were plated on slides via cytospin centrifugation and stained using MCDH. The proportions of seven hemocyte types were analyzed: **H** (Hyalinocytes), **ML** (Macrophage-Like cells), **BBL** (Basophilic Blast-Like cells), **ABL** (Acidophilic Blast-Like cells), **SGC** (Small Granule Cells), **BGC** (Big Granule Cells), and **VC** (Vesicular Cells). The data show heterogeneity in the hemocyte composition across individual oysters and cell types. No significant differences were observed between oysters from the two origins (p-value > 0.05). The table below the graph provides a detailed breakdown of the mean percentages and standard deviations of each hemocyte type across the two oyster groups, highlighting the variability observed within and between these populations.

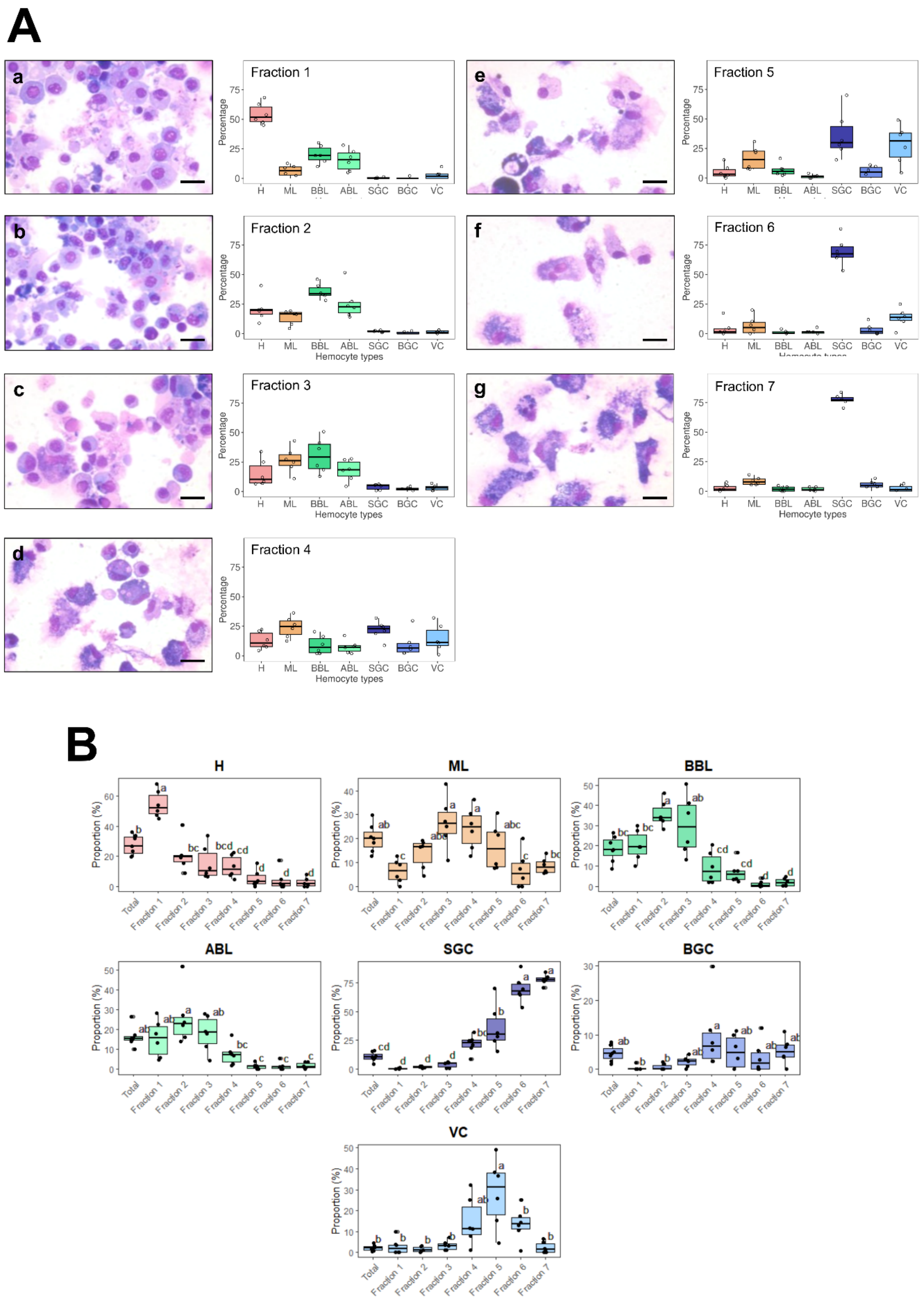


**Figure S4. Statistical significance of enrichment in different hemocyte types in Percoll gradient fractions. (A) a,b,c,d,e,f, and g** : MCDH staining of cells from the 7 fractions isolated from the percoll gradient in (B). Scale bar : 10 µm. **Fraction 1 to 7 :** Quantification of the different types of hemocytes found in each of the 7 fractions from 5 independent fractionation experiments. **(B)** Statistical significance of enrichment of different hemocyte types in Percoll gradient fractions. Results are from six independent experiments. Statistical significance is indicated by letters, as different letters indicate a significant difference between enrichments of cell types within the Percoll density gradient fractions (ANOVA, Tukey's test, p-value <0.05). Hyalinocytes (**H**) were significantly enriched in the first fraction compared to the other fractions and compared to unsorted hemocytes. However, they were significantly depleted in fractions 4, 5, 6 and 7 compared to unsorted hemocytes. Macrophage-like cells (**ML**) were significantly enriched in fractions 3 and 4 compared to fractions 1, 6 and 7. They were depleted in fraction 1 compared to unsorted hemocytes. Acidophilic blasts (**ABL**) were significantly depleted in fractions 4, 5, 6, and 7 compared to unsorted hemocytes. Basophilic blasts (**BBL**) were significantly enriched in fractions 2 and 3 compared to fractions 4, 5, 6, and 7 and in fraction 1 compared to fractions 6 and 7. Compared to unsorted hemocytes, basophilic blasts (**BBL**) were significantly enriched in fraction 2 and depleted in fractions 6 to 7. Small granule cells (**SGC**) were significantly depleted in fractions 1, 2, and 3 compared to fractions 4, 5, 6, and 7, and also significantly depleted in fractions 4 and 5 compared to fractions 6 and 7. In addition, small granule cells (**SGC**) were significantly enriched in fractions 5, 6, and 7 compared to unsorted hemocytes. The distribution of the big granule cells (**BGC**) showed a significant enrichment in fraction 4, compared to fractions 1 and 2, but no significant changes were observed with unsorted hemocytes. Vesicular cells (**VC**) were enriched in fraction 5 compared to fractions 1, 2, 3, 6, 7 and unsorted hemocytes in fractions 1, 2, and 3 compared to fraction 5 and enriched in fraction 5 compared to fraction 7 and unsorted hemocytes.

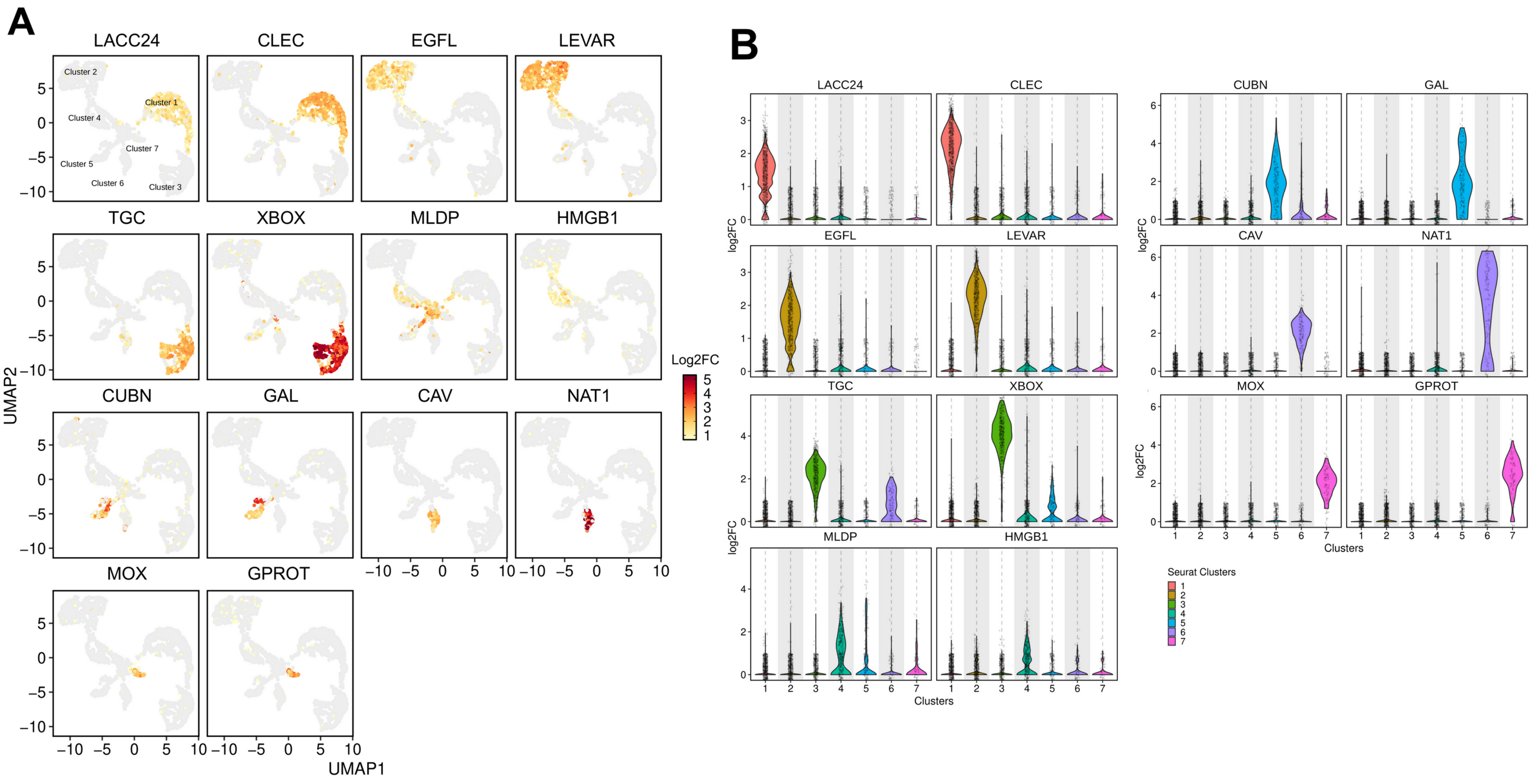


**Figure S5. Cluster specificity and expression level of the 14 selected cluster markers A)** Identification of cells expressing the selected markers on the UMAP plot. Positive cells are colored according to the Log2FC value. LACC24 and CLEC are specific of cluster 1, EGFL and LEVAR of cluster 2, TGC and XBOX of cluster 3, MLDP and HMGB1 of cluster 4, CUBN and GAL of cluster 5, CAV and NAT1 of cluster 6 and MOX and GPROT of cluster 7. **LACC24** : Laccase 24, **CLEC** : C-type lectin domain-containing protein, **EGFL** : EGF-like domain-containing protein 8, **LEVAR** : Putative regulator of levamisole receptor-1, **TGC** : TGc domain-containing protein, **XBOX** : X-box binding protein-like protein, **MLDP** : ML domain-containing protein, **HMGB1** : High mobility group protein B1, **CUBN** : Cubilin, **GAL** : Galectin, **CAV** : Caveolin, **NAT1** : Natterin-1, **MOX** : DBH-like monooxygenase protein 1, **GPROT** : G protein receptor F1-2 domain-containing protein. **B)** Violin graph showing the average expression level (Log2FC) of the 14 selected marker transcripts specific to the different scRNA-seq clusters.

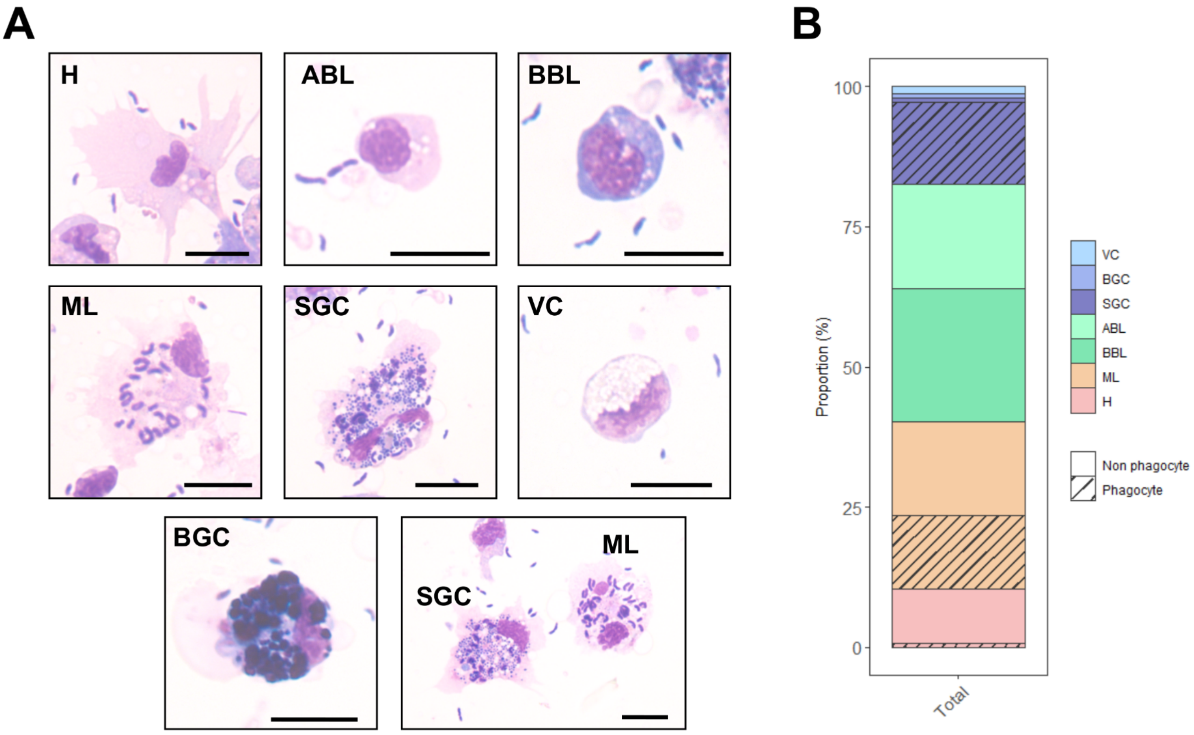


**Figure S6. Measurement of the ability of *C.gigas* hemocytes to phagocytose *Vibrio tasmaniensis* LMG20012$^{T}$.** Oyster hemocytes were challenged with non-pathogenic *Vibrio tasmaniensis* LMG20012$^{T}$ and phagocytosis was measured by observing intracellular bacteria after MCDH staining. **(A)** MCDH staining of hemocytes after phagocytosis assay. Scale bar : 10µm. **(B)** Bar plot showing the proportion of each cell type and the proportion of phagocytic cells. **H** : Hyalinocytes, **ABL** : Acidophilic Blast-Like cells, **BBL** : Basophilic Blast-Like cells, **ML** : Macrophage-Like cells, **SGC** : Small Granule Cells, **VC** : Vesicular Cells and **BGC** : Big Granule Cells.

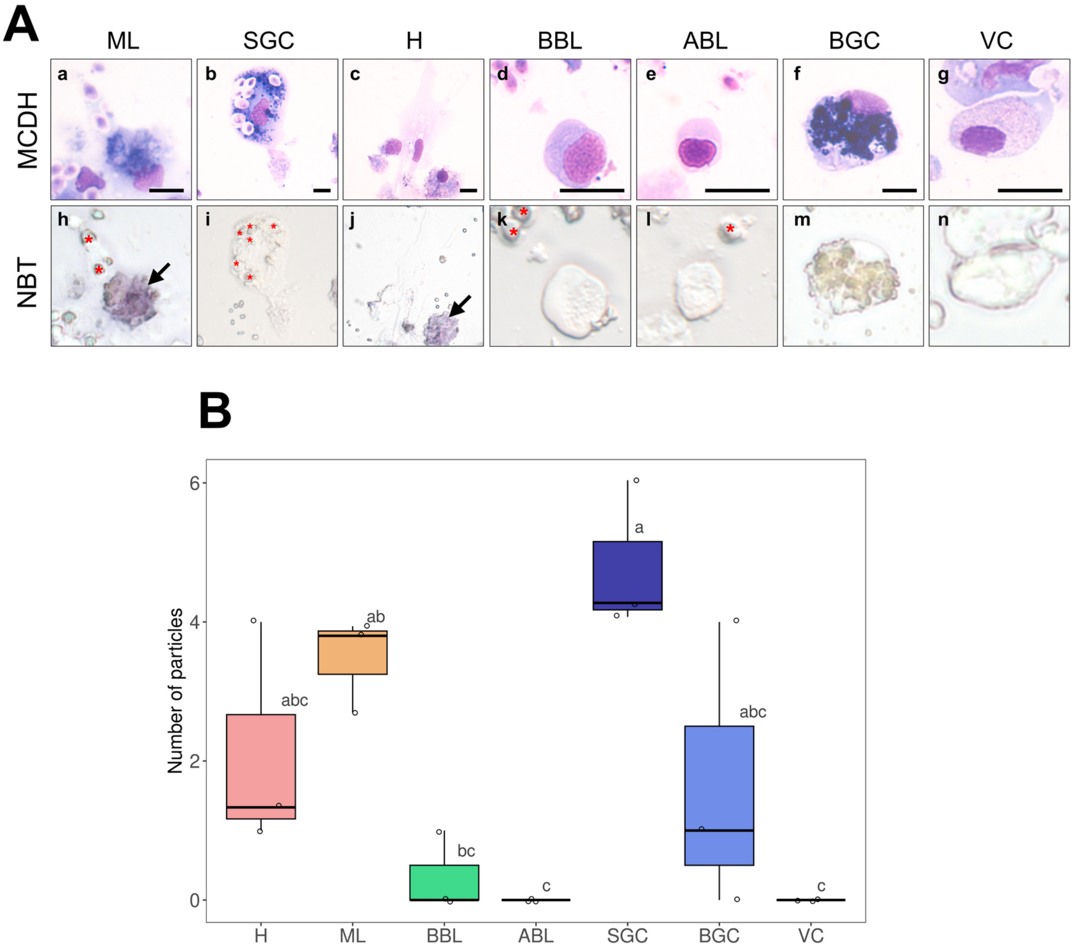


**Figure S7. NBT (NitroBlueTetrazolium) staining of oyster hemolymph exposed to zymosan particles. (A)** Hemocyte morphology after MCDH staining : Macrophage Like **(a)**, Small Granule Cells **(b)**, Hyalinocyte **(c)**, Basophilic **(d)** and Acidophilic **(e)** Blast cells, Big Granule Cells **(f)** and Vesicular Cells **(g)**. NBT staining of the different hemocyte types **(h-n)**. Red stars show zymosan and bacteria particles. Black arrows identify Macrophage-Like cells. Scale bar : 10 µm. **(B)** Results of quantification of the phagocytic activity of each cell type and number of zymosan particles per cell type. The graph shows the result of 3 independent experiments.

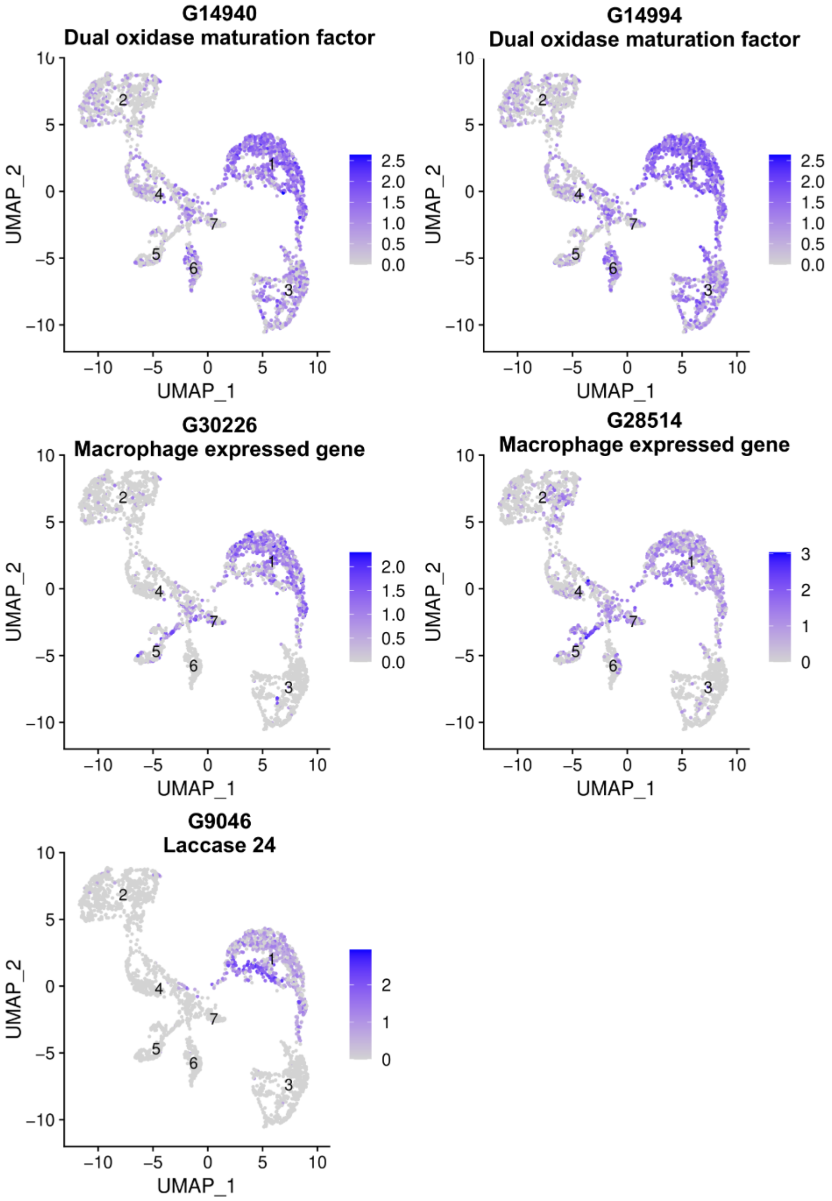


**Figure S8. Uniform Manifold Approximation and Projection (UMAP) plots of cells expressing Macrophage-Like markers.** Cluster numbers are indicated on each cluster. Each point in the UMAP plot represents a single hemocyte, and the clustering of these points reveals the distinct transcriptional profiles of macrophage-like specific markers within the hemocyte population.

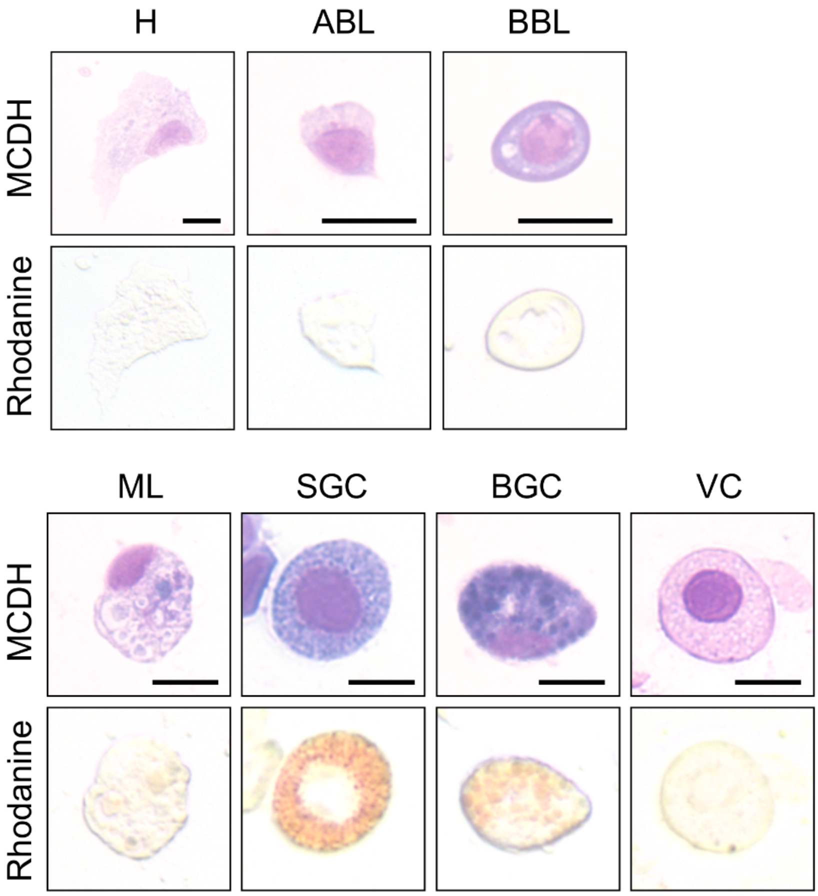


**Figure S9. Labeling of intracellular copper stores in C.gigas hemocytes.** MCDH (upper panels) and rhodanine (lower panels) staining of oyster hemocytes to reveal copper accumulation. Cells were first processed for copper staining and then stained according to MCDH protocol. **H** : Hyalinocytes, **ABL** : Acidophilic Blast-Like cells, **BBL** : Basophilic Blast-Like cells, **ML** : Macrophage-Like cells, **SGC** : Small Granule Cells, **VC** : Vesicular Cells and **BGC** : Big Granule Cells. Bar : 10µm.

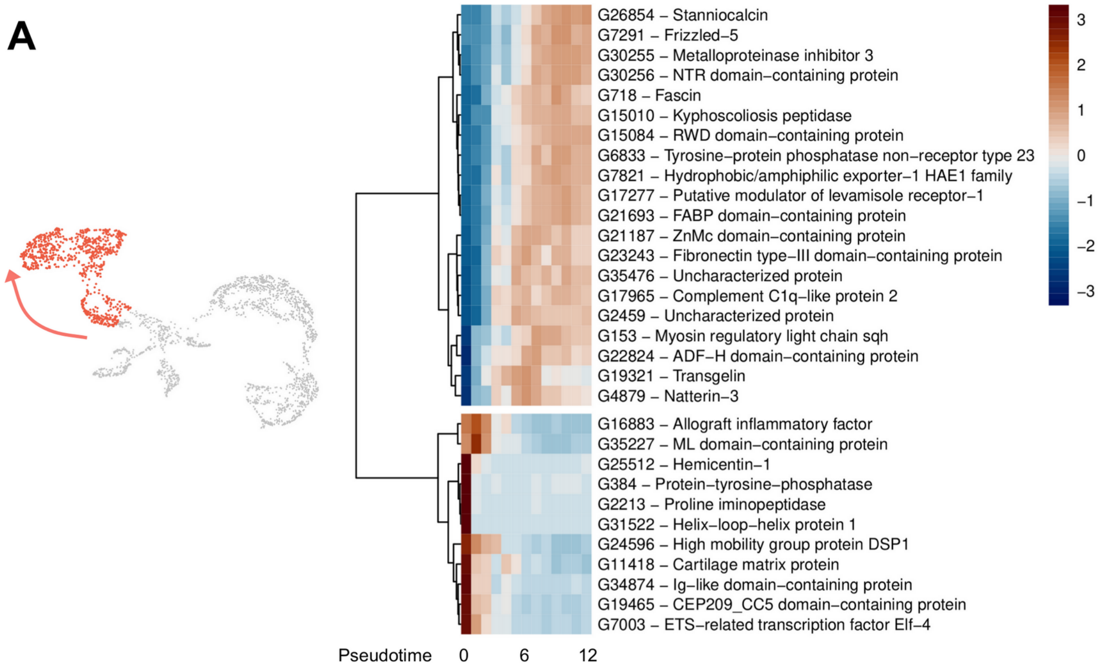
A
G26854 – Stanniocalcin
G7291 – Frizzled–5
G30255 – Metalloproteinase inhibitor 3
G30256 – NTR domain–containing protein
G718 – Fascin
G15010 – Kyphoscoliosis peptidase
G15084 – RWD domain–containing protein
G6833 – Tyrosine–protein phosphatase non–receptor type 23
G7821 – Hydrophobic/amphiphilic exporter–1 HAE1 family
G17277 – Putative modulator of levamisole receptor–1
G21693 – FABP domain–containing protein
G21187 – ZnMc domain–containing protein
G23243 – Fibronectin type–III domain–containing protein
G35476 – Uncharacterized protein
G17965 – Complement C1q–like protein 2
G2459 – Uncharacterized protein
G153 – Myosin regulatory light chain sqh
G22824 – ADF–H domain–containing protein
G19321 – Transgelin
G4879 – Natterin–3
G16883 – Allograft inflammatory factor
G35227 – ML domain–containing protein
G25512 – Hemicentin–1
G384 – Protein–tyrosine–phosphatase
G2213 – Proline iminopeptidase
G31522 – Helix–loop–helix protein 1
G24596 – High mobility group protein DSP1
G11418 – Cartilage matrix protein
G34874 – Ig–like domain–containing protein
G19465 – CEP209_CC5 domain–containing protein
G7003 – ETS–related transcription factor Elf–4
Pseudotime 0 6 12
3
2
1
0
−1
−2
−3

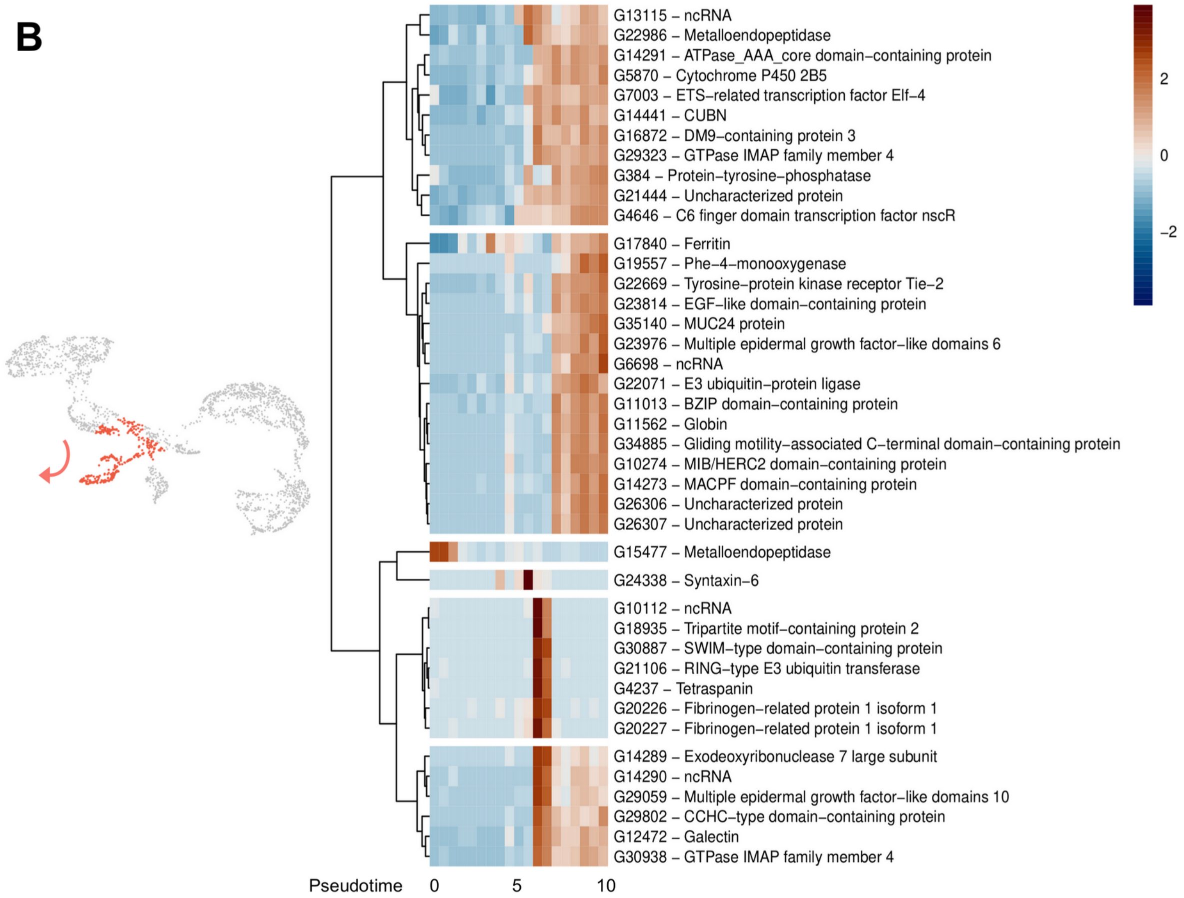
B
G13115 – ncRNA
G22986 – Metalloendopeptidase
G14291 – ATPase_AAA_core domain–containing protein
G5870 – Cytochrome P450 2B5
G7003 – ETS–related transcription factor Elf–4
G14441 – CUBN
G16872 – DM9–containing protein 3
G29323 – GTPase IMAP family member 4
G384 – Protein–tyrosine–phosphatase
G21444 – Uncharacterized protein
G4646 – C6 finger domain transcription factor nscR
G17840 – Ferritin
G19557 – Phe–4–monooxygenase
G22669 – Tyrosine–protein kinase receptor Tie–2
G23814 – EGF–like domain–containing protein
G35140 – MUC24 protein
G23976 – Multiple epidermal growth factor–like domains 6
G6698 – ncRNA
G22071 – E3 ubiquitin–protein ligase
G11013 – BZIP domain–containing protein
G11562 – Globin
G34885 – Gliding motility–associated C–terminal domain–containing protein
G10274 – MIB/HERC2 domain–containing protein
G14273 – MACPF domain–containing protein
G26306 – Uncharacterized protein
G26307 – Uncharacterized protein
G15477 – Metalloendopeptidase
G24338 – Syntaxin–6
G10112 – ncRNA
G18935 – Tripartite motif–containing protein 2
G30887 – SWIM–type domain–containing protein
G21106 – RING–type E3 ubiquitin transferase
G4237 – Tetraspanin
G20226 – Fibrinogen–related protein 1 isoform 1
G20227 – Fibrinogen–related protein 1 isoform 1
G14289 – Exodeoxyribonuclease 7 large subunit
G14290 – ncRNA
G29059 – Multiple epidermal growth factor–like domains 10
G29802 – CCHC–type domain–containing protein
G12472 – Galectin
G30938 – GTPase IMAP family member 4
Pseudotime 0 5 10
2
0
−2

**C**

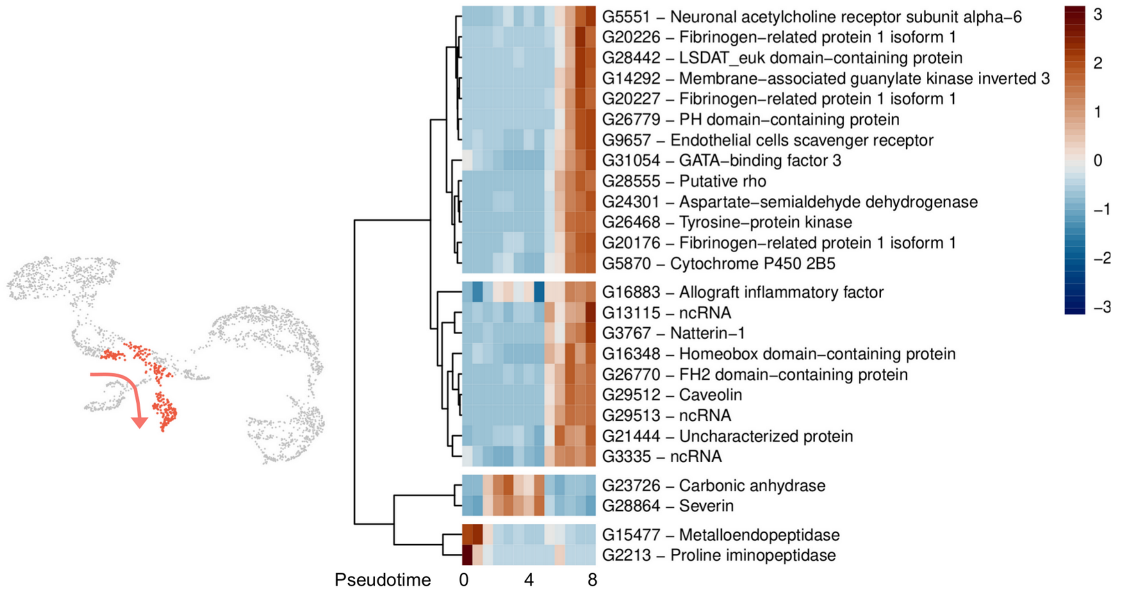
G5551 – Neuronal acetylcholine receptor subunit alpha–6
G20226 – Fibrinogen–related protein 1 isoform 1
G28442 – LSDAT_euk domain–containing protein
G14292 – Membrane–associated guanylate kinase inverted 3
G20227 – Fibrinogen–related protein 1 isoform 1
G26779 – PH domain–containing protein
G9657 – Endothelial cells scavenger receptor
G31054 – GATA–binding factor 3
G28555 – Putative rho
G24301 – Aspartate–semialdehyde dehydrogenase
G26468 – Tyrosine–protein kinase
G20176 – Fibrinogen–related protein 1 isoform 1
G5870 – Cytochrome P450 2B5
G16883 – Allograft inflammatory factor
G13115 – ncRNA
G3767 – Natterin–1
G16348 – Homeobox domain–containing protein
G26770 – FH2 domain–containing protein
G29512 – Caveolin
G29513 – ncRNA
G21444 – Uncharacterized protein
G3335 – ncRNA
G23726 – Carbonic anhydrase
G28864 – Severin
G15477 – Metalloendopeptidase
G2213 – Proline iminopeptidase
Pseudotime 0 4 8
3
2
1
0
–1
–2
–3

**D**

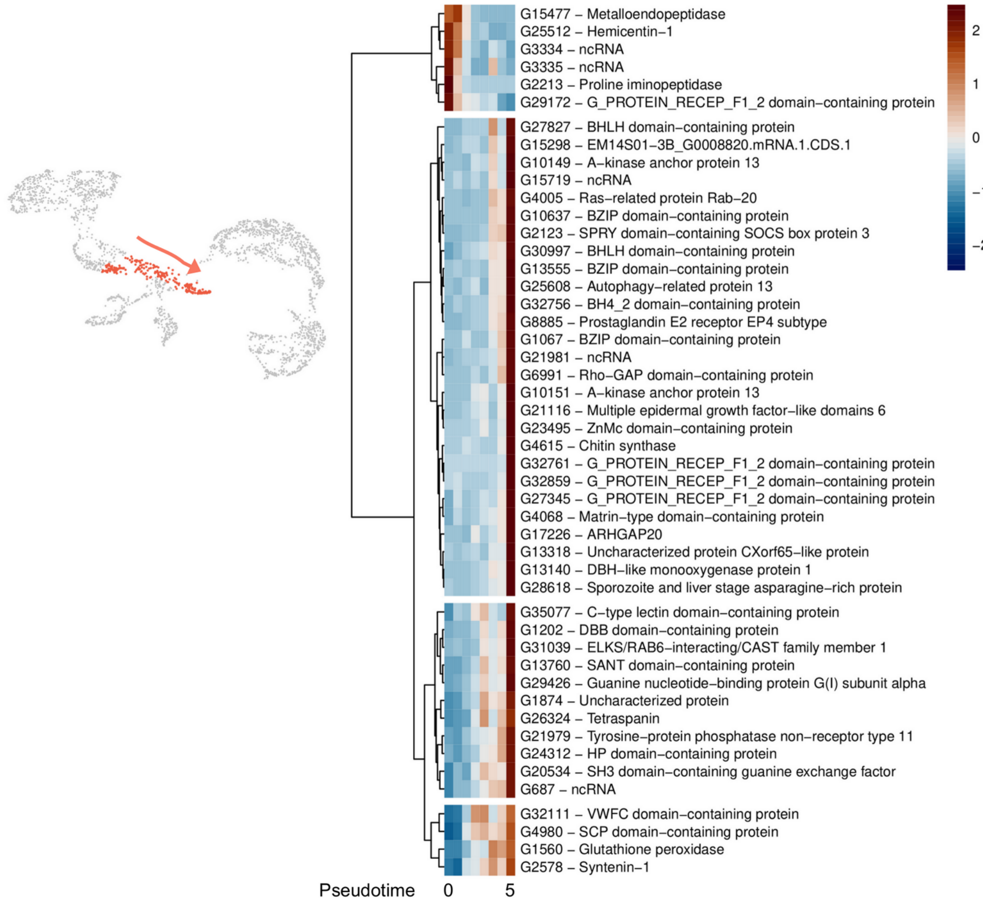
G15477 – Metalloendopeptidase
G25512 – Hemicentin–1
G3334 – ncRNA
G3335 – ncRNA
G2213 – Proline iminopeptidase
G29172 – G_PROTEIN_RECEP_F1_2 domain–containing protein
G27827 – BHLH domain–containing protein
G15298 – EM14S01–3B_G0008820.mRNA.1.CDS.1
G10149 – A–kinase anchor protein 13
G15719 – ncRNA
G4005 – Ras–related protein Rab–20
G10637 – BZIP domain–containing protein
G2123 – SPRY domain–containing SOCS box protein 3
G30997 – BHLH domain–containing protein
G13555 – BZIP domain–containing protein
G25608 – Autophagy–related protein 13
G32756 – BH4_2 domain–containing protein
G8885 – Prostaglandin E2 receptor EP4 subtype
G1067 – BZIP domain–containing protein
G21981 – ncRNA
G6991 – Rho–GAP domain–containing protein
G10151 – A–kinase anchor protein 13
G21116 – Multiple epidermal growth factor–like domains 6
G23495 – ZnMc domain–containing protein
G4615 – Chitin synthase
G32761 – G_PROTEIN_RECEP_F1_2 domain–containing protein
G32859 – G_PROTEIN_RECEP_F1_2 domain–containing protein
G27345 – G_PROTEIN_RECEP_F1_2 domain–containing protein
G4068 – Matrin–type domain–containing protein
G17226 – ARHGAP20
G13318 – Uncharacterized protein CXorf65–like protein
G13140 – DBH–like monooxygenase protein 1
G28618 – Sporozoite and liver stage asparagine–rich protein
G35077 – C–type lectin domain–containing protein
G1202 – DBB domain–containing protein
G31039 – ELKS/RAB6–interacting/CAST family member 1
G13760 – SANT domain–containing protein
G29426 – Guanine nucleotide–binding protein G(I) subunit alpha
G1874 – Uncharacterized protein
G26324 – Tetraspanin
G21979 – Tyrosine–protein phosphatase non–receptor type 11
G24312 – HP domain–containing protein
G20534 – SH3 domain–containing guanine exchange factor
G687 – ncRNA
G32111 – VWFC domain–containing protein
G4980 – SCP domain–containing protein
G1560 – Glutathione peroxidase
G2578 – Syntenin–1
Pseudotime 0 5
2
1
0
–1
–2

**E**

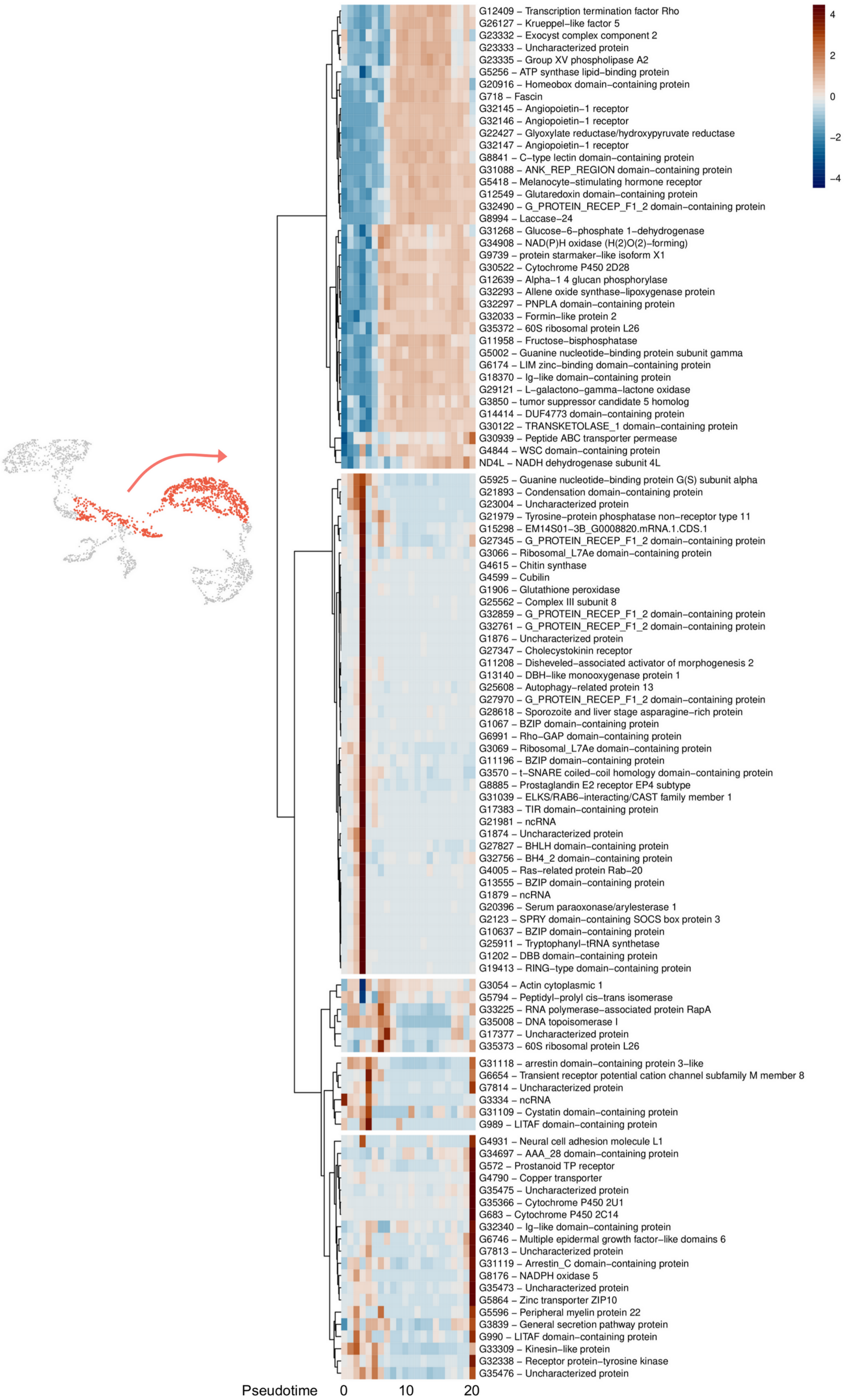
G12409 – Transcription termination factor Rho
G26127 – Krueppel–like factor 5
G23332 – Exocyst complex component 2
G23333 – Uncharacterized protein
G23335 – Group XV phospholipase A2
G5256 – ATP synthase lipid–binding protein
G20916 – Homeobox domain–containing protein
G718 – Fascin
G32145 – Angiopoietin–1 receptor
G32146 – Angiopoietin–1 receptor
G22427 – Glyoxylate reductase/hydroxypyruvate reductase
G32147 – Angiopoietin–1 receptor
G8841 – C–type lectin domain–containing protein
G31088 – ANK_REP_REGION domain–containing protein
G5418 – Melanocyte–stimulating hormone receptor
G12549 – Glutaredoxin domain–containing protein
G32490 – G_PROTEIN_RECEP_F1_2 domain–containing protein
G8994 – Laccase–24
G31268 – Glucose–6–phosphate 1–dehydrogenase
G34908 – NAD(P)H oxidase (H(2)O(2)–forming)
G9739 – protein starmaker–like isoform X1
G30522 – Cytochrome P450 2D28
G12639 – Alpha–1 4 glucan phosphorylase
G32293 – Allene oxide synthase–lipoxygenase protein
G32297 – PNPLA domain–containing protein
G32033 – Formin–like protein 2
G35372 – 60S ribosomal protein L26
G11958 – Fructose–bisphosphatase
G5002 – Guanine nucleotide–binding protein subunit gamma
G6174 – LIM zinc–binding domain–containing protein
G18370 – Ig–like domain–containing protein
G29121 – L–galactono–gamma–lactone oxidase
G3850 – tumor suppressor candidate 5 homolog
G14414 – DUF4773 domain–containing protein
G30122 – TRANSKETOLASE_1 domain–containing protein
G30939 – Peptide ABC transporter permease
G4844 – WSC domain–containing protein
ND4L – NADH dehydrogenase subunit 4L
G5925 – Guanine nucleotide–binding protein G(S) subunit alpha
G21893 – Condensation domain–containing protein
G23004 – Uncharacterized protein
G21979 – Tyrosine–protein phosphatase non–receptor type 11
G15298 – EM14S01–3B_G0008820.mRNA.1.CDS.1
G27345 – G_PROTEIN_RECEP_F1_2 domain–containing protein
G3066 – Ribosomal_L7Ae domain–containing protein
G4615 – Chitin synthase
G4599 – Cubilin
G1906 – Glutathione peroxidase
G25562 – Complex III subunit 8
G32859 – G_PROTEIN_RECEP_F1_2 domain–containing protein
G32761 – G_PROTEIN_RECEP_F1_2 domain–containing protein
G1876 – Uncharacterized protein
G27347 – Cholecystokinin receptor
G11208 – Disheveled–associated activator of morphogenesis 2
G13140 – DBH–like monooxygenase protein 1
G25608 – Autophagy–related protein 13
G27970 – G_PROTEIN_RECEP_F1_2 domain–containing protein
G28618 – Sporozoite and liver stage asparagine–rich protein
G1067 – BZIP domain–containing protein
G6991 – Rho–GAP domain–containing protein
G3069 – Ribosomal_L7Ae domain–containing protein
G11196 – BZIP domain–containing protein
G3570 – t–SNARE coiled–coil homology domain–containing protein
G8885 – Prostaglandin E2 receptor EP4 subtype
G31039 – ELKS/RAB6–interacting/CAST family member 1
G17383 – TIR domain–containing protein
G21981 – ncRNA
G1874 – Uncharacterized protein
G27827 – BHLH domain–containing protein
G32756 – BH4_2 domain–containing protein
G4005 – Ras–related protein Rab–20
G13555 – BZIP domain–containing protein
G1879 – ncRNA
G20396 – Serum paraoxonase/arylesterase 1
G2123 – SPRY domain–containing SOCS box protein 3
G10637 – BZIP domain–containing protein
G25911 – Tryptophanyl–tRNA synthetase
G1202 – DBB domain–containing protein
G19413 – RING–type domain–containing protein
G3054 – Actin cytoplasmic 1
G5794 – Peptidyl–prolyl cis–trans isomerase
G33225 – RNA polymerase–associated protein RapA
G35008 – DNA topoisomerase I
G17377 – Uncharacterized protein
G35373 – 60S ribosomal protein L26
G31118 – arrestin domain–containing protein 3–like
G6654 – Transient receptor potential cation channel subfamily M member 8
G7814 – Uncharacterized protein
G3334 – ncRNA
G31109 – Cystatin domain–containing protein
G989 – LITAF domain–containing protein
G4931 – Neural cell adhesion molecule L1
G34697 – AAA_28 domain–containing protein
G572 – Prostanoid TP receptor
G4790 – Copper transporter
G35475 – Uncharacterized protein
G35366 – Cytochrome P450 2U1
G683 – Cytochrome P450 2C14
G32340 – Ig–like domain–containing protein
G6746 – Multiple epidermal growth factor–like domains 6
G7813 – Uncharacterized protein
G31119 – Arrestin_C domain–containing protein
G8176 – NADPH oxidase 5
G35473 – Uncharacterized protein
G5864 – Zinc transporter ZIP10
G5596 – Peripheral myelin protein 22
G3839 – General secretion pathway protein
G990 – LITAF domain–containing protein
G33309 – Kinesin–like protein
G32338 – Receptor protein–tyrosine kinase
G35476 – Uncharacterized protein
Pseudotime
0
10
20
4
2
0
−2
−4

**F**

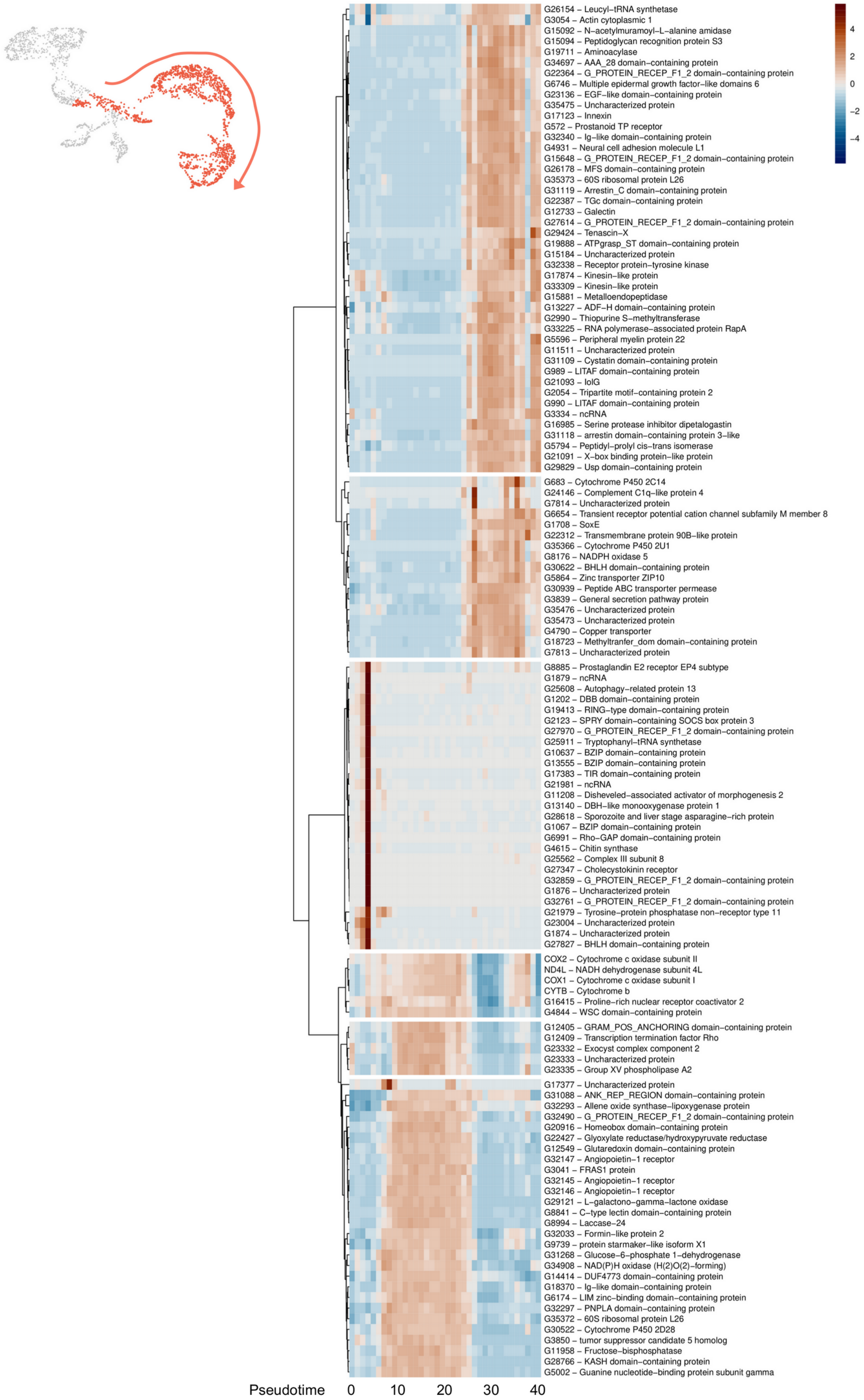
G26154 – Leucyl–tRNA synthetase
G3054 – Actin cytoplasmic 1
G15092 – N–acetylmuramoyl–L–alanine amidase
G15094 – Peptidoglycan recognition protein S3
G19711 – Aminoacylase
G34697 – AAA_28 domain–containing protein
G22364 – G_PROTEIN_RECEP_F1_2 domain–containing protein
G6746 – Multiple epidermal growth factor–like domains 6
G23136 – EGF–like domain–containing protein
G35475 – Uncharacterized protein
G17123 – Innexin
G572 – Prostanoid TP receptor
G32340 – Ig–like domain–containing protein
G4931 – Neural cell adhesion molecule L1
G15648 – G_PROTEIN_RECEP_F1_2 domain–containing protein
G26178 – MFS domain–containing protein
G35373 – 60S ribosomal protein L26
G31119 – Arrestin_C domain–containing protein
G22387 – TGc domain–containing protein
G12733 – Galectin
G27614 – G_PROTEIN_RECEP_F1_2 domain–containing protein
G29424 – Tenascin–X
G19888 – ATPgrasp_ST domain–containing protein
G15184 – Uncharacterized protein
G32338 – Receptor protein–tyrosine kinase
G17874 – Kinesin–like protein
G33309 – Kinesin–like protein
G15881 – Metalloendopeptidase
G13227 – ADF–H domain–containing protein
G2990 – Thiopurine S–methyltransferase
G33225 – RNA polymerase–associated protein RapA
G5596 – Peripheral myelin protein 22
G11511 – Uncharacterized protein
G31109 – Cystatin domain–containing protein
G989 – LITAF domain–containing protein
G21093 – IolG
G2054 – Tripartite motif–containing protein 2
G990 – LITAF domain–containing protein
G3334 – ncRNA
G16985 – Serine protease inhibitor dipetalogastin
G31118 – arrestin domain–containing protein 3–like
G5794 – Peptidyl–prolyl cis–trans isomerase
G21091 – X–box binding protein–like protein
G29829 – Usp domain–containing protein
G683 – Cytochrome P450 2C14
G24146 – Complement C1q–like protein 4
G7814 – Uncharacterized protein
G6654 – Transient receptor potential cation channel subfamily M member 8
G1708 – SoxE
G22312 – Transmembrane protein 90B–like protein
G35366 – Cytochrome P450 2U1
G8176 – NADPH oxidase 5
G30622 – BHLH domain–containing protein
G5864 – Zinc transporter ZIP10
G30939 – Peptide ABC transporter permease
G3839 – General secretion pathway protein
G35476 – Uncharacterized protein
G35473 – Uncharacterized protein
G4790 – Copper transporter
G18723 – Methyltranfer_dom domain–containing protein
G7813 – Uncharacterized protein
G8885 – Prostaglandin E2 receptor EP4 subtype
G1879 – ncRNA
G25608 – Autophagy–related protein 13
G1202 – DBB domain–containing protein
G19413 – RING–type domain–containing protein
G2123 – SPRY domain–containing SOCS box protein 3
G27970 – G_PROTEIN_RECEP_F1_2 domain–containing protein
G25911 – Tryptophanyl–tRNA synthetase
G10637 – BZIP domain–containing protein
G13555 – BZIP domain–containing protein
G17383 – TIR domain–containing protein
G21981 – ncRNA
G11208 – Disheveled–associated activator of morphogenesis 2
G13140 – DBH–like monooxygenase protein 1
G28618 – Sporozoite and liver stage asparagine–rich protein
G1067 – BZIP domain–containing protein
G6991 – Rho–GAP domain–containing protein
G4615 – Chitin synthase
G25562 – Complex III subunit 8
G27347 – Cholecystokinin receptor
G32859 – G_PROTEIN_RECEP_F1_2 domain–containing protein
G1876 – Uncharacterized protein
G32761 – G_PROTEIN_RECEP_F1_2 domain–containing protein
G21979 – Tyrosine–protein phosphatase non–receptor type 11
G23004 – Uncharacterized protein
G1874 – Uncharacterized protein
G27827 – BHLH domain–containing protein
COX2 – Cytochrome c oxidase subunit II
ND4L – NADH dehydrogenase subunit 4L
COX1 – Cytochrome c oxidase subunit I
CYTB – Cytochrome b
G16415 – Proline–rich nuclear receptor coactivator 2
G4844 – WSC domain–containing protein
G12405 – GRAM_POS_ANCHORING domain–containing protein
G12409 – Transcription termination factor Rho
G23332 – Exocyst complex component 2
G23333 – Uncharacterized protein
G23335 – Group XV phospholipase A2
G17377 – Uncharacterized protein
G31088 – ANK_REP_REGION domain–containing protein
G32293 – Allene oxide synthase–lipoxygenase protein
G32490 – G_PROTEIN_RECEP_F1_2 domain–containing protein
G20916 – Homeobox domain–containing protein
G22427 – Glyoxylate reductase/hydroxypyruvate reductase
G12549 – Glutaredoxin domain–containing protein
G32147 – Angiopoietin–1 receptor
G3041 – FRAS1 protein
G32145 – Angiopoietin–1 receptor
G32146 – Angiopoietin–1 receptor
G29121 – L–galactono–gamma–lactone oxidase
G8841 – C–type lectin domain–containing protein
G8994 – Laccase–24
G32033 – Formin–like protein 2
G9739 – protein starmaker–like isoform X1
G31268 – Glucose–6–phosphate 1–dehydrogenase
G34908 – NAD(P)H oxidase (H(2)O(2)–forming)
G14414 – DUF4773 domain–containing protein
G18370 – Ig–like domain–containing protein
G6174 – LIM zinc–binding domain–containing protein
G32297 – PNPLA domain–containing protein
G35372 – 60S ribosomal protein L26
G30522 – Cytochrome P450 2D28
G3850 – tumor suppressor candidate 5 homolog
G11958 – Fructose–bisphosphatase
G28766 – KASH domain–containing protein
G5002 – Guanine nucleotide–binding protein subunit gamma
4
2
0
−2
−4
Pseudotime
0
10
20
30
40

**Figure S10. Results of scRNA-seq trajectory analysis using Monocle3.** Analysis was performed using cluster 4 as the zero pseudotime. **(A) to (F)** Heatmaps representing the level of normalized expression of genes along the trajectory. **(A)** Lineage from cluster 4 cells to hyalinocytes, **(B)** to cluster 5 cells, **(C)** to cluster 6 cells, **(D)** to vesicular cells, **(E)** to macrophage-like cells and **(F)** to small granule cells. For each lineage, cell trajectories are shown in red and heat maps of pseudo time-dependent genes are shown. Blue indicates low expression, and red indicates high expression. Pseudotime flows from left to right.

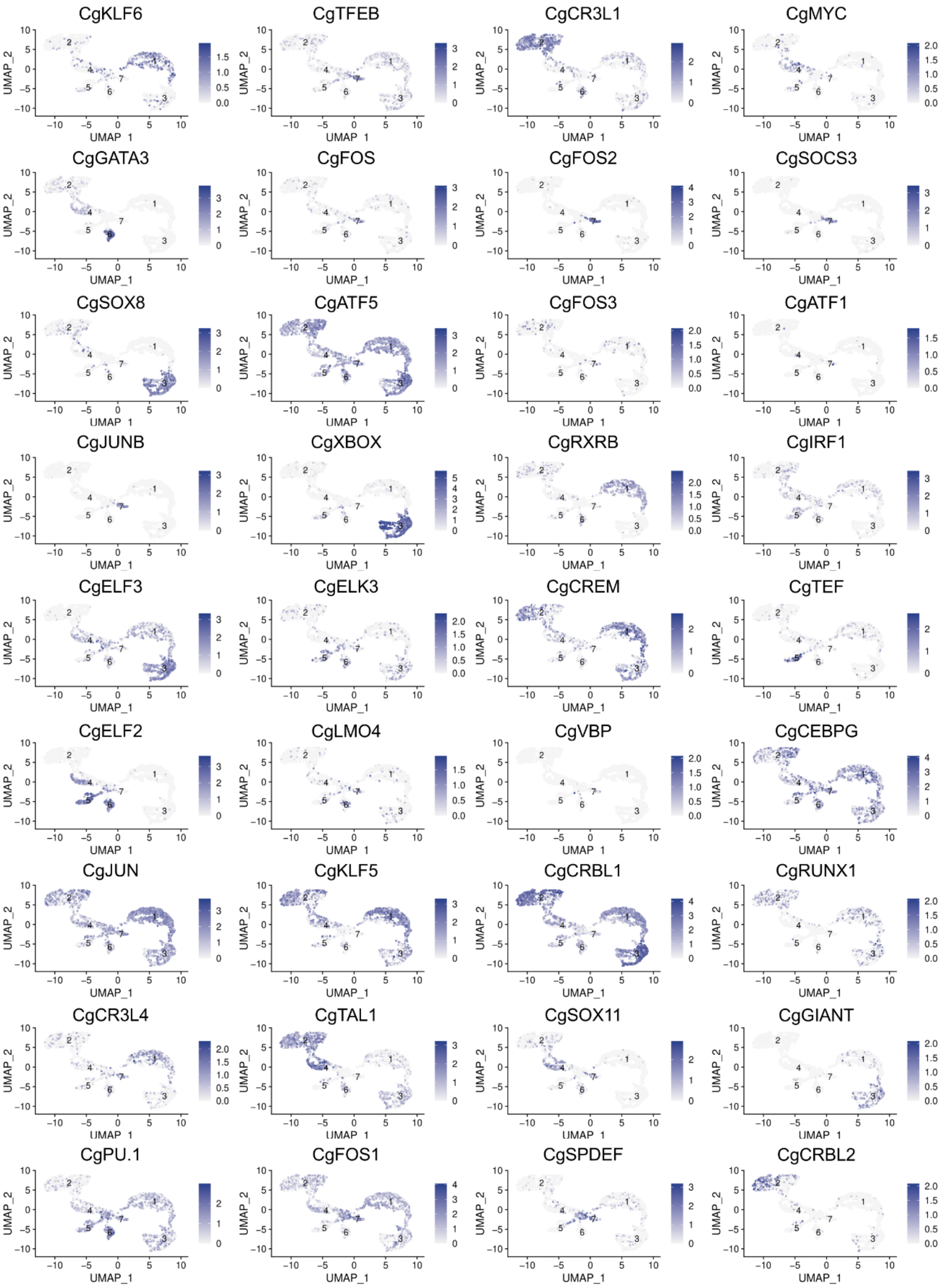


**Figure S11. Uniform Manifold Approximation and Projection (UMAP) plots of cells expressing transcription factors.** 28 UMAP representation for the transcription factors identified in the scRNA-seq dataset. Each UMAP plot shows cells expressing the transcription factor in purple. Log2FC expression level is also reported.

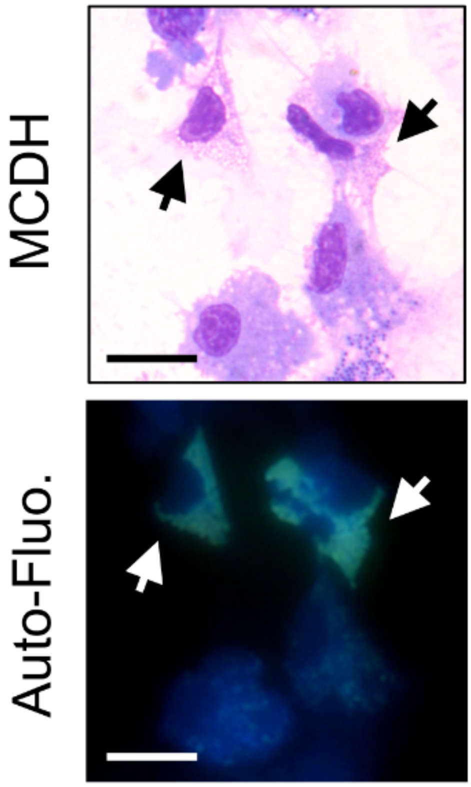


**Figure S12. Observation of autofluorescence of vesicular cells in hemolymph.** Freshly punctured total hemolymph was cytospun, directly observed under a microscope using a DAPI filter set (DAPI Blue ex : 350/50 nm, DC : 400 nm and em : 460/50 nm) and then processed for MCDH staining. Arrows indicate autofluorescent cells. Scale bar : 10 µm
